# High-Throughput, Lysis-free Screening for Sulfatase Activity Using *Escherichia coli* Autodisplay in Microdroplets

**DOI:** 10.1101/479162

**Authors:** Bert van Loo, Magdalena Heberlein, Philip Mair, Anastasia Zinchenko, Jan Schüürmann, Bernard D. G. Eenink, Josephin M. Holstein, Carina Dilkaute, Joachim Jose, Florian Hollfelder, Erich Bornberg-Bauer

## Abstract

Directed evolution of enzymes toward improved catalytic performance has become a powerful tool in protein engineering. To be effective, a directed evolution campaign requires the use of high-throughput screening. In this study we describe the development of a high-throughput lysis-free procedure to screen for improved sulfatase activity by combining microdroplet-based single-variant activity sorting with *E. coli* autodisplay. For the first step in a 4-step screening procedure we quantitatively screened >10^5^ variants of the homodimeric arylsulfatase from *Silicibacter pomeroyi* (*Sp*AS1), displayed on the *E. coli* cell surface, for improved sulfatase activity using fluorescence activated droplet sorting. Display of the sulfatase variants on living *E. coli* cells ensured the continuous linkage of genotype and phenotype during droplet sorting and allowed for direct recovery by simple regrowth of the sorted cells. The use of autodisplay on living cells simplified and reduced the degree of liquid handling during all steps in the screening procedure to the single event of simply mixing substrate and cells. The percentage of apparent improved variants was enriched >10-fold as a result of droplet sorting. We ultimately identified 25 *Sp*AS1-variants with improved performance toward 4-nitrophenyl sulfate (up to 6.2-fold) and/or fluorescein disulfate (up to 30-fold). In *Sp*AS1 variants with improved performance toward the bulky fluorescein disulfate, many of the beneficial mutations occur in residues that form hydrogen bonds between α-helices in the C-terminal oligomerization region, suggesting a non-trivial, previously unknown role for the dimer interface in shaping the substrate binding site of *Sp*AS1.

**Table of contents graphic:** 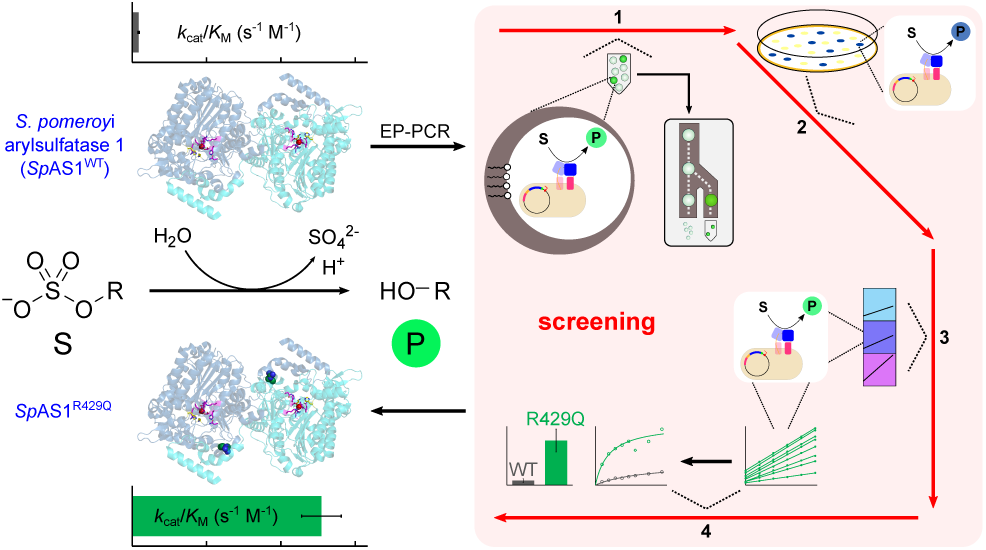

## Introduction

Directed evolution comprises repeated cycles of mutation,^1–4^ followed by selection of variants with improved desired function.^5,6^ Directed evolution has been used successfully to improve several enzyme properties such as enantioselectivity,^7–9^ operational stability,^10–12^ and catalytic efficiency.^13,14^ Effective enzyme engineering by directed evolution requires fast, sensitive and reliable screening of large libraries of enzyme variants in order to select the very few improved variants present in these libraries. Current methods for large-scale library testing such as colorimetric colony screening^8,15^ or growth selection-based systems^16,17^ suffer from narrow dynamic ranges, i.e. they are either too sensitive (almost all variants appear equally active) or too selective (almost all variants appear inactive), limiting the degree of improvement that can be reached. Microtiterplate-based screening methods have a broader dynamic range,^18^ but require extensive instrumentation and consumables.

Over the last decade, water-in-oil emulsion microdroplets have been established as a cost and resource effective alternative for efficient high-throughput screening. Microdroplets are *in vitro* compartments and are, in essence, miniaturized reaction vessels that can be created at high frequency (>8 kHz) and in high numbers (∼ 10^7^ per day).^19–22^ Each droplet contains, as a result of a controllable Poisson distribution, on average less than one library variant (in this study λ = 0.35). In such a library, the corresponding protein is produced *in vitro* from a single plasmid copy,^23,24^ or by single cells, typically bacteria^21,25^ or yeast,^19^ each harboring one library variant. Reaction progress in microdroplets can be monitored at high frequencies (>2 kHz), using either commercial fluorescence-activated cell sorters^22^ (for sorting water-in-oil-in-water double emulsion droplets), or custom-made on-chip sorters.^19,21,25^ For the latter a range of optical signals reporting on reaction progress is now available,^26^ based on fluorescence,^27^ absorbance,^28^ or anisotropy.^29^

In many cases, directed evolution campaigns that screened for single variants required an intrinsic physical link between genotype and phenotype,^30^ e.g. using phage display for antibody evolution. Such display methods can also be adapted to screen for single turnover reactions,^31–34^ but screening for multiple turnover catalysis, the hallmark of efficient enzymes, is not possible with these methods. In droplet-based screening procedures, the phenotype can be the result of, e.g. a fluorescent reaction product formed by multiple turnovers of substrate. During droplet sorting this phenotype stays linked to its genotype (the variant DNA) due to the boundary of the micro-droplet. For both display and many of the droplet-based methods, the genotypes need assistance in order to be recovered after droplet sorting: e.g. for phage display transfection of fresh *E. coli* is required and for lysis-based screening with single *E. coli* cells the plasmid DNA needs to be re-transformed.

In our study we combine a self-replicating genotype, i.e. a living *E. coli* cell, that autodisplays our enzyme of interest on its surface, with activity-based droplet sorting to select for improved performance of multiple turnover reactions. This combination has thus far only been reported for yeast.^19^ Autodisplay uses the β-barrel and the linker domain of a natural type V autotransporter protein for displaying the passenger, i.e. the enzyme, to the cell surface (Figure S1).^35,36^ Anchoring of the enzyme within the outer membrane by the β-barrel results in its mobility on the cell surface. Thus, we present a system that can be used in an easy-to-use lab organism (*E. coli*, as compared to yeast) that can deal with homo-oligomeric proteins such as *Sp*AS1.^37^ Thus far, autodisplay has been used to screen enzyme libraries by binding artificial substrates at the cell surface,^38,39^ but has not yet been used for direct detection of enzymatic turnover. We describe a screening procedure for selecting improved variants of the well-characterized arylsulfatase from *Silicibacter pomeroyi* (*Sp*AS1).^37,40^ We screened two separate error-prone PCR-generated mutant libraries based on *Sp*AS1^WT^ for improved sulfatase activity using a lysis-free 4-step screening procedure (Figure 1). We identified in total 25 *Sp*AS1 variants with significantly improved activity toward fluorescein disulfate (1.3-to up to 30-fold improved *k*_cat_/*K*_M_) and/or 4-nitrophenyl sulfate (1.7-to 6.2-fold improved *k*_cat_/*K*_M_), despite the latter substrate only being used for screening in the final 2 steps. The droplet-based screening for improved activity was successfully used to enrich the libraries for improved activity toward the fluorescein disulfate as well as enriching the library for improved sulfatase activity in general.

**Figure 1:**
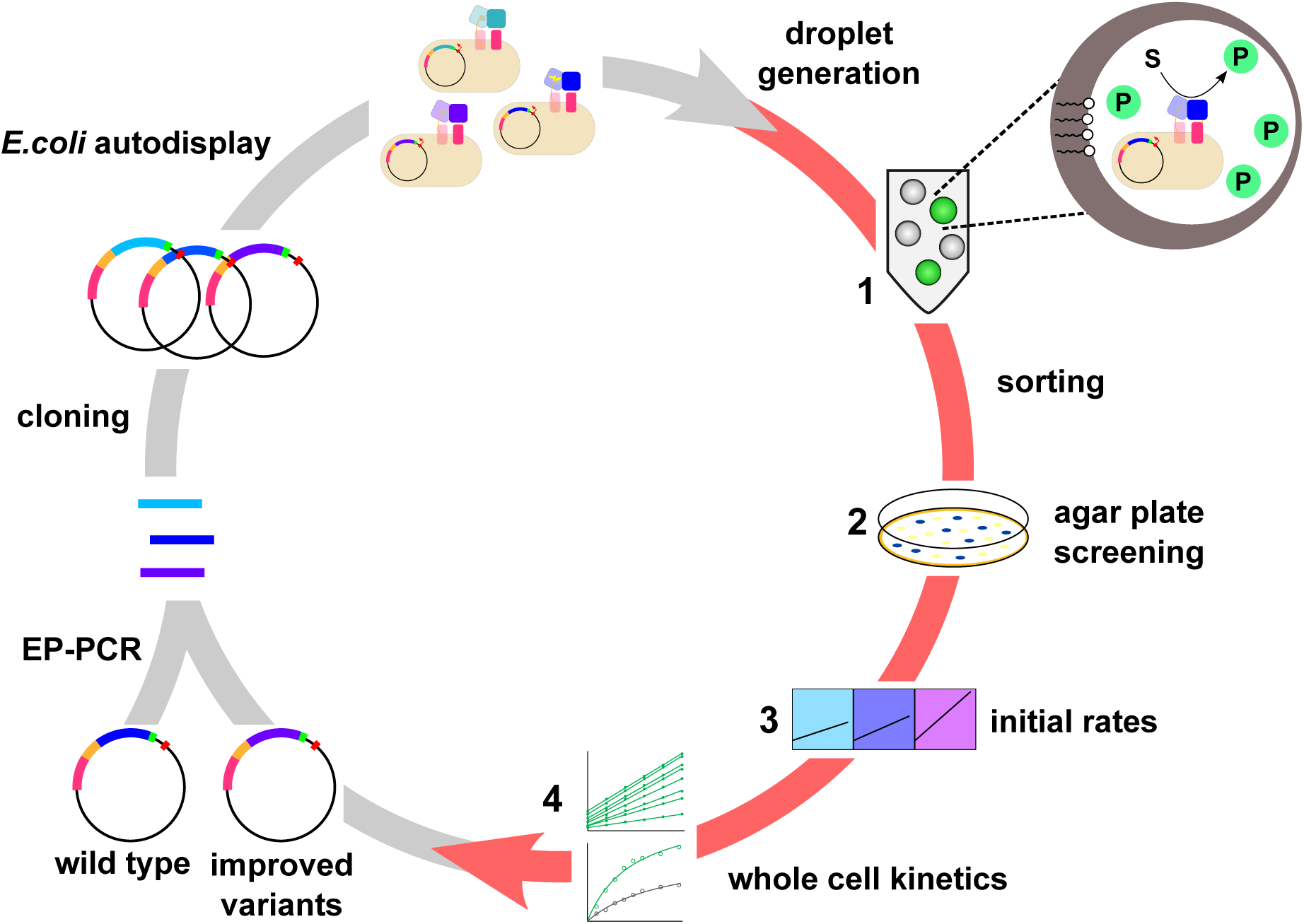
Directed evolution workflow for screening for improved catalytic activity of an autodisplayed dimeric arylsulfatase. A library of arylsulfatase mutants, generated by error-prone PCR (EP-PCR), is displayed on the surface of *E. coli* (grey arrow). Mutants with improved arylsulfatase activity were selected by a 4-step screening procedure (red arrow): **1**) droplet generation and fluorescence assisted droplet sorting (FADS), followed by **2**) agarplate activity screening, testing of initial reaction rates and **4**) whole cell-based approximation of Michaelis-Menten kinetics. For a more detailed description of all 4 step see Figure 2 and 3.

## Results and Discussion

### Display of active *Sp*AS1 on the surface of *E. coli*

The dimeric arylsulfatase *Sp*AS1^37,40^ was fused to a gene III secretion signal (N-terminus) and an autotransporter protein (C-terminus) (Figure 2 and Figure S1), as described previously for a wide variety of other proteins.^35,41,42^ Successful display of active *Sp*AS1 on the surface of *E. coli* was first shown by testing whole cells expressing the autodisplay construct for activity toward sulfate monoesters **2a** and **3a** (Figure 3). Cells displaying the catalytically inactive *Sp*AS1 C53A variant (*k*_cat_/*K*_M_ *∼* 10^5^-fold below wild type) and cells containing the empty autodisplay expression vector showed no detectable hydrolysis of any of the sulfate monoesters used in this study. SDS-PAGE analysis of the membrane proteins of cells expressing the *Sp*AS1-autotransporter construct showed that the latter is attached to *E. coli* membrane. The same analysis after treatment of cells expressing the *Sp*AS1-autotransporter complex with a protease showed removal of the *Sp*AS1-domain, indicating that *Sp*AS1 is indeed displayed toward the outside of the *E. coli* cells (Figure S2).

**Figure 2:**
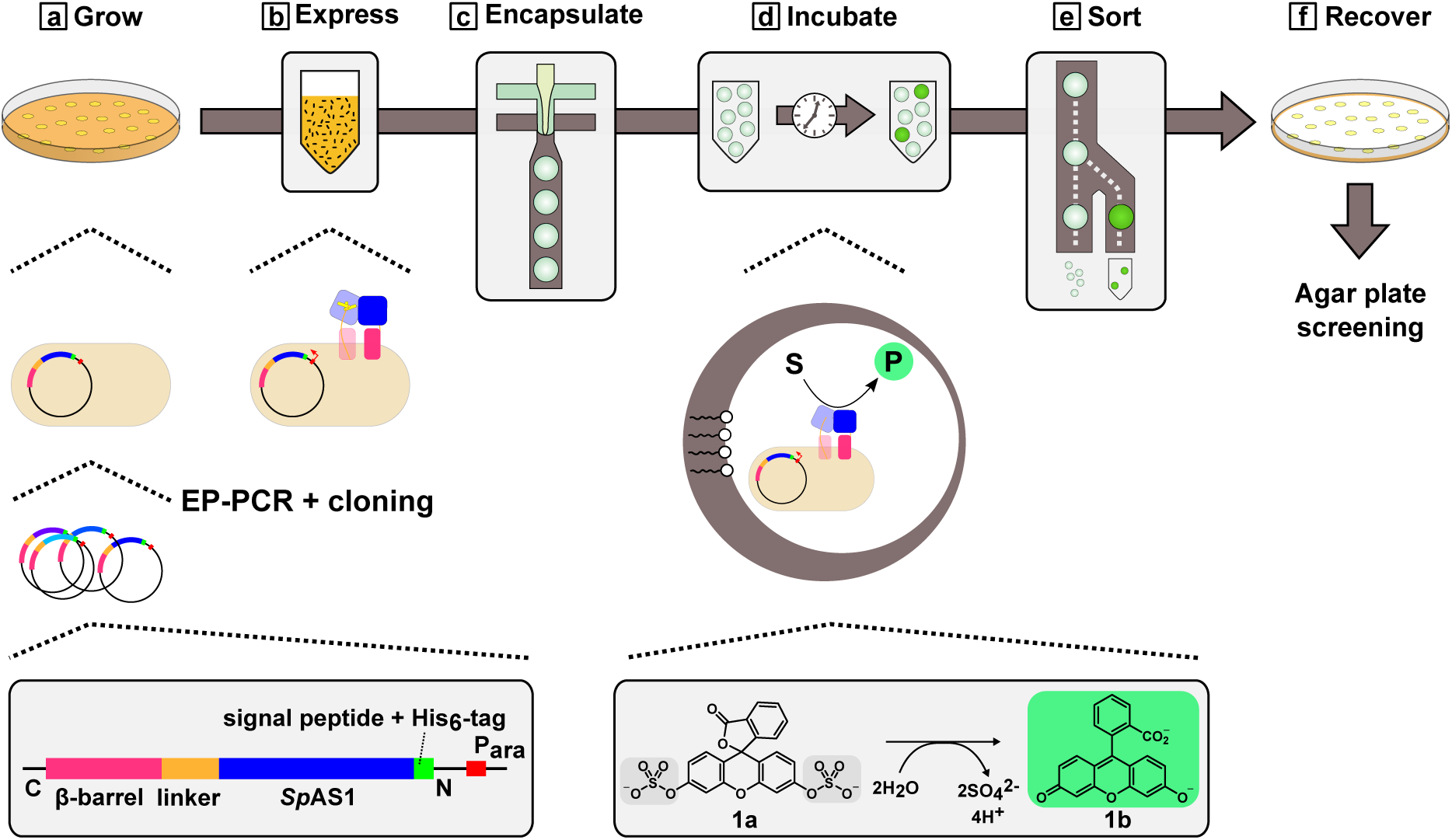
Screening of a library of autodisplayed *Sp*AS1-variants for improved sulfatase activity in microdroplets (step 1 in Figure 1). (**a**) *E. coli* cells containing an error-prone PCR generated mutant library of autodisplayed *Sp*AS1 are grown into full-sized colonies. (**b**) The resulting colonies are resuspended in liquid medium and expression of the autodisplayed *Sp*AS1 variants is induced in bulk overnight. (**c**) The cells are washed and resuspended in reaction buffer and are encapsulated in water-in-oil microdroplets, each containing 5 µM fluorescein disulfate **1a**) and a single *E. coli* cell (the latter in ∼ 1 in every 3 droplets). (**d**) The droplets are left in the dark at room temperature overnight and (**e**) are subsequently sorted using a custom-built fluorescence activated droplet sorter (FADS). Droplets with a fluorescence signal above the cut-off value are collected directly into a vial containing a mixture of oil, surfactant and recovery medium. (**f**) Once the desired number of sorted events is reached, additional recovery medium containing 0.5% (w/v) pyruvate is added and the cells are allowed to recover for two hours. The recovered cells were plated onto a nitrocellulose filter sitting on top of solid medium and were left to grow into full-sized colonies overnight, ready for screening step 2 (Figure 3A). Details regarding screening step 1, library sizes and coverage can be found in the supporting information (Figure S3-S5, Table S1).

**Figure 3:**
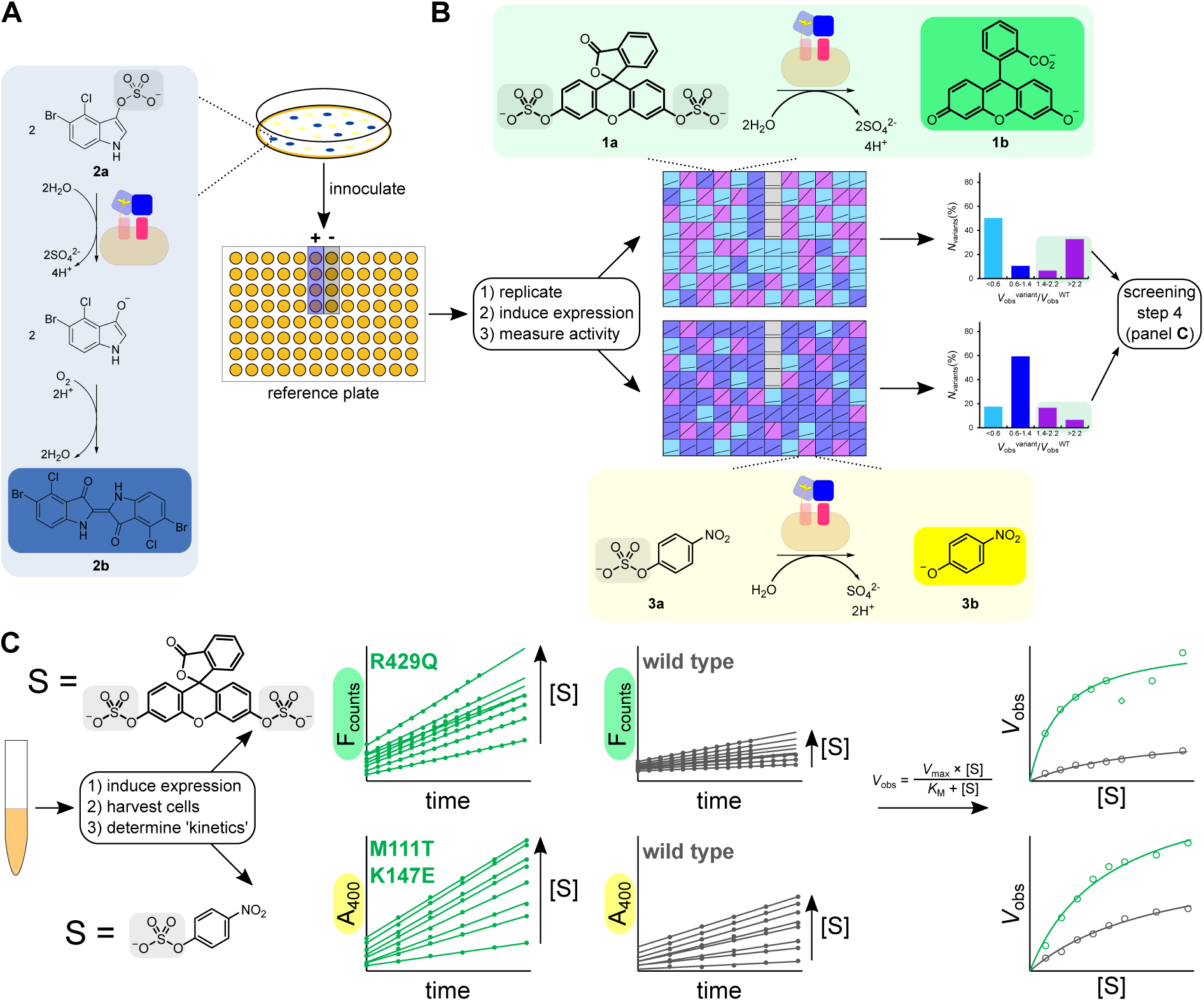
Screening of autodisplayed *Sp*AS1 variants for improved sulfatase activity (steps 2-4 in Figure 1). (**A**) Screening step 2: colonies sitting on top of a nitrocellulose filter expressing active autodisplayed *Sp*AS1 variants are tested for activity toward X-sulfate **2a**. Colonies displaying active variants that turned blue, due to the formation of chromophore **2b**, within 30 minutes were used to inoculate an individual well in a reference microtiter plate containing growth medium. After overnight growth the reference plate was replicated into another microtiterplate in which expression of the selected *Sp*AS1 variants was induced overnight. (**B**) Screening step 3: activity toward fluorescein disulfate **1b** and 4-nitrophenyl sulfate **3a** were determined side-by-side using fluorescence and absorbance respectively, corrected for the measured cell density (OD_600_). Wild-type and an inactive *Sp*AS1-variant were used for reference (+) and as a negative control (-) respectively. Given the considerable variation in the wild-type control (± 30%), only variants with >1.4-fold improved activity were carried over to the final screening step. (**C**) Screening step 4: approximation of Michaelis-Menten parameters of autodisplayed *Sp*AS1 variants for fluorescein disulfate **1b** and 4-nitrophenyl sulfate **3a**. Initial rates of product formation (*V*_obs_) catalyzed by cells displaying *Sp*AS1 variants were determined for 8 different substrate concentrations ([S]). The resulting data were fitted to Michaelis-Menten kinetics and the resulting parameters *V*_max_, *K*_M_ and *V*_max_/*K*_M_ were compared to those of autodisplayed *Sp*AS1^WT^ (see Figure S6-S8 for details)

### Screening of an error-prone-PCR-generated library of displayed *Sp*AS1 variants

We created two different libraries of autodisplayed *Sp*AS1 variants, containing mutations introduced using error-prone PCR with mutagenic nucleotides dPTP and 8-oxo-dGTP respectively (Table S1). Expression of both libraries was induced in *E. coli* in a high cell density bulk solution (Figure 2). The cells displaying the *Sp*AS1 variants were encapsulated in water-in-oil microdroplets containing fluorescein disulfate (sulfate monoester **1a**, Figure 2), according to a Poisson distribution (with on average 0.35 cells per droplet, e.g. λ *≈* 0.35). Hydrolysis of fluorescein disulfate **1a** into the fluorophore fluorescein (product **1b**, Figure 2) by an active autodisplayed *Sp*AS1 variant allows for fluorescence activated droplet sorting (FADS).^21,25,27^ We sorted through >10^6^ microdroplets until ∼ 4000 microdroplets with at least 2-fold higher fluorescence signal than wild type were collected (Table S1, Figure S5). The sorted positive cells were re-grown into full-sized colonies on a nitrocellulose filter placed on top of solid growth medium (Figure 2).

The regrown colonies were tested in a second screening step for activity toward sulfate **2a** (Figure 3A). This so-called blue-white screening was done to weed out false positives caused by sorting artifacts and completely inactive clones that ‘hitchhike’ in multiple (mostly double) occupancy droplets. The latter are unavoidable, due to the nature of the Poisson distribution of the cells encapsulated in water-in-oil microdroplets. Colonies in which *Sp*AS1-catalyzed conversion of sulfate monoester **2a** resulted in formation of the blue chromophore **2b** (Figure 3A, Figure S6) within 30 minutes, were selected for the third screening step.

For screening step 3, we tested for improved activity toward both fluorescein disulfate (**1a**) and 4-nitrophenyl sulfate (**3a**) in a side-by-side fashion, i.e. activity measurements were done on the same day with the same overnight expression experiment carried out in microtiterplates (Figure 3B and S7). All *Sp*AS1 variants that showed >1.4-fold increased activity toward either sulfate monoester (**1a** or **3a**) relative to *Sp*AS1^WT^ (Figure S9), were selected for the fourth and final step of the screening procedure.

In the fourth and final step we determined the Michaelis-Menten parameters toward fluorescein disulfate **1a** and 4-nitrophenyl sulfate **3a** for the autodisplayed *Sp*AS1 variants (Figure 3C and S8). Altogether, 40 *Sp*AS1 variants with improved whole cell second order rates (*V*_max_/*K*_M_) toward fluorescein disulfate **1a** and/or 4-nitrophenyl sulfate **3a** were selected and their mutations were determined by DNA sequencing. Out of those 40 selected variants, 3 were found twice and 10 variants contained only silent mutations (Figure S10). These silent mutations could possibly enhance expression levels. Retesting of these variants could not unambiguously rule out that they were expressed at significantly higher levels than autodisplayed *Sp*AS1^WT^, for which a ±50% variation in *V*_max_/*K*_M_ was observed (Table S2-S5). The 27 unique *Sp*AS1 variants that each contain at least 1 amino acid substitution were characterized further (see below for details).

The micro-droplet sorting step, during which droplets with a fluorescence signal 2-fold higher than wild-type are selected (Figure S5B and C), is expected to enrich for *E. coli* cells displaying highly active *Sp*AS1 variants. In order to assess the effectiveness of the droplet-sorting-based enrichment we also performed the colony-based activity screening (step 2 in Figure 3A) on the ‘naive’ library, i.e. the library prior to microdroplet-based screening. We compared the number of colonies scoring as positive with this assay, i.e. turn blue 30 minutes after exposure to sulfate monoester **2a**), before and after droplet sorting. Prior to sorting, ∼ 28% of the colonies in the dPTP-generated library turned blue after 30 minutes, whereas after sorting this percentage rises to ∼ 42% (∼ 1.5-fold enrichment). For the ‘naive’ 8-oxo-dGTP-generated library fewer colonies scored as positive before sorting (∼ 20%) compared to the dPTP library. Furthermore, the 8-oxo-dGTP library shows no apparent enrichment of the percentage of *Sp*AS1 variants that are active toward sulfate monoester **2a** at a level above the threshold after sorting. Like with the actual screening procedure, we also tested the blue colonies from the ‘naive’ library for activity toward fluorescein disulfate **1a** and 4-nitrophenyl sulfate **3a** in a side-by-side comparison (Figure 3, Figure S7).

The correlation between the activities of the *Sp*AS1 variants toward both fluorescein disulfate **1a** and 4-nitrophenyl sulfate **3a** are positively correlated for both the ‘naive’ and the ‘sorted’ libraries (Figure 4). However, the inclusion of droplet sorting for improved activity toward fluorescein disulfate **1a** improves this correlation considerably. Furthermore, both ‘sorted’ libraries show a clear enrichment of the fraction of autodisplayed *Sp*AS1 variants with improved activity toward fluorescein disulfate **1a** and/or 4-nitrophenyl sulfate **3a** (Figure 5). In particular for the dPTP-generated library, the enrichment for fluorescein disulfate **1a** is stronger than for 4-nitrophenyl sulfate **3a** (Figure 5B), as would be expected since the sorted library has been screened for improved performance toward fluorescein disulfate **1a**. For the 8-oxo-dGTP-generated library, the enrichment of autodisplayed *Sp*AS1 variants with improved activity toward 4-nitrophenyl sulfate **3a** appears to be much stronger than the enrichment of autodisplayed *Sp*AS1 variants with improved activity toward fluorescein disulfate **1a** (Figure 5D). However, for the latter the number of improved variants in the tested ‘naive’ library was so low (1 count for each category, Figure 5C), that the actual level of enrichment could easily be only half of what we observe. Taking the latter into account, together with the more detrimental mutational load used for the 8-oxo-dGTP-generated library (Table S1), the droplet sorting procedure may in this case simply enrich for *Sp*AS1 variants that are generally active.

**Figure 4:**
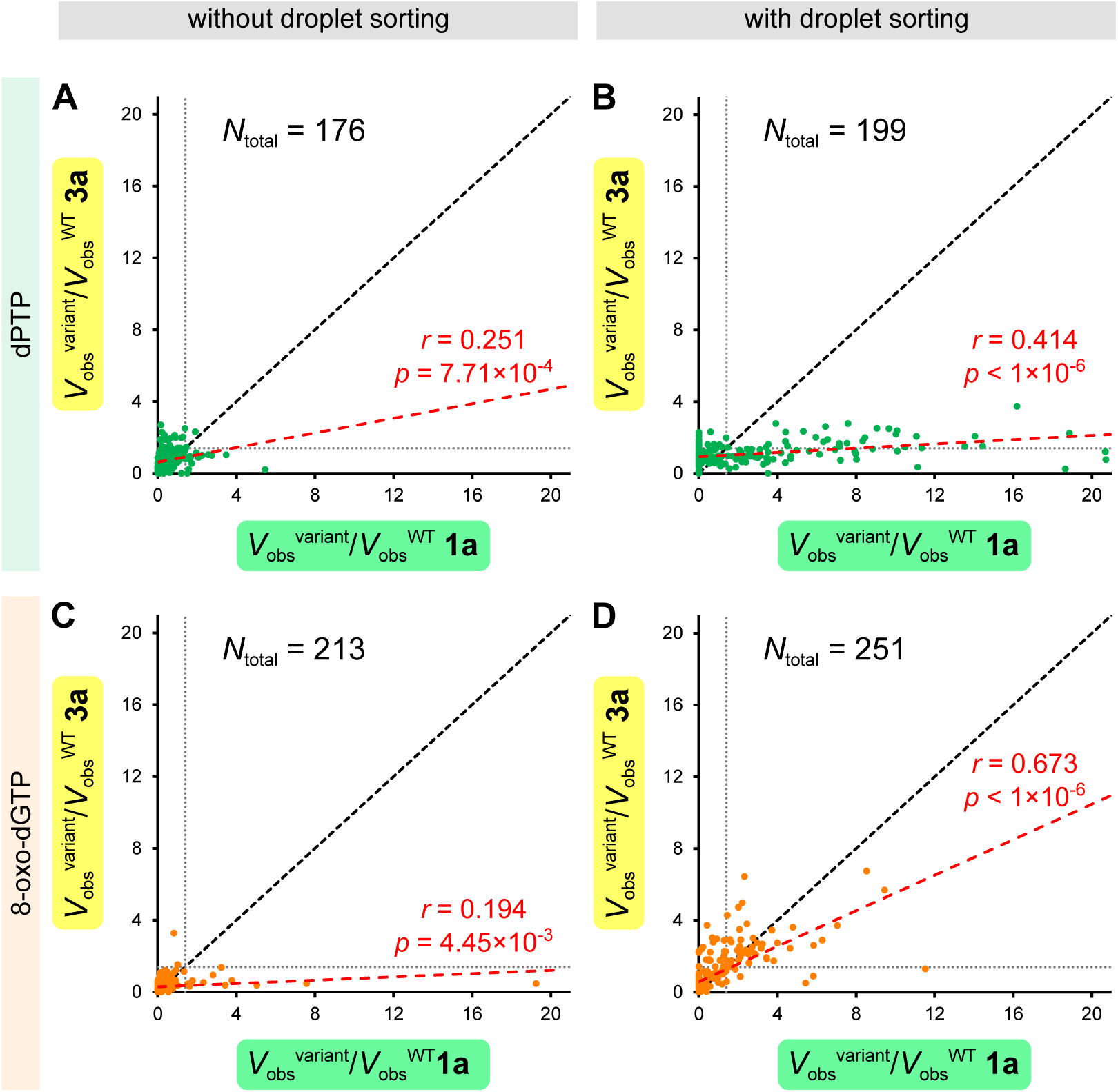
Cross-correlation for whole cell activity measurements for *Sp*AS1 variants toward fluorescein disulfate **1a** and 4-nitrophenyl sulfate **3a**. Both the dPTP and the 8-oxo-dGTP generated libraries show a significant positive correlation between both activities (*p* > 5 ×10^-2^). The black dashed diagonals represent the theoretical situation in which all mutations would have an equal effect on the catalytic performance toward both substrates. The dotted vertical and horizontal lines indicate *V*_obs_^variant^/*V*_obs_^WT^ = 1.4 for each substrate. The red dashed lines indicate the linear correlations for which the correlation coefficient (*r*) and *p*-value are indicated.

**Figure 5:**
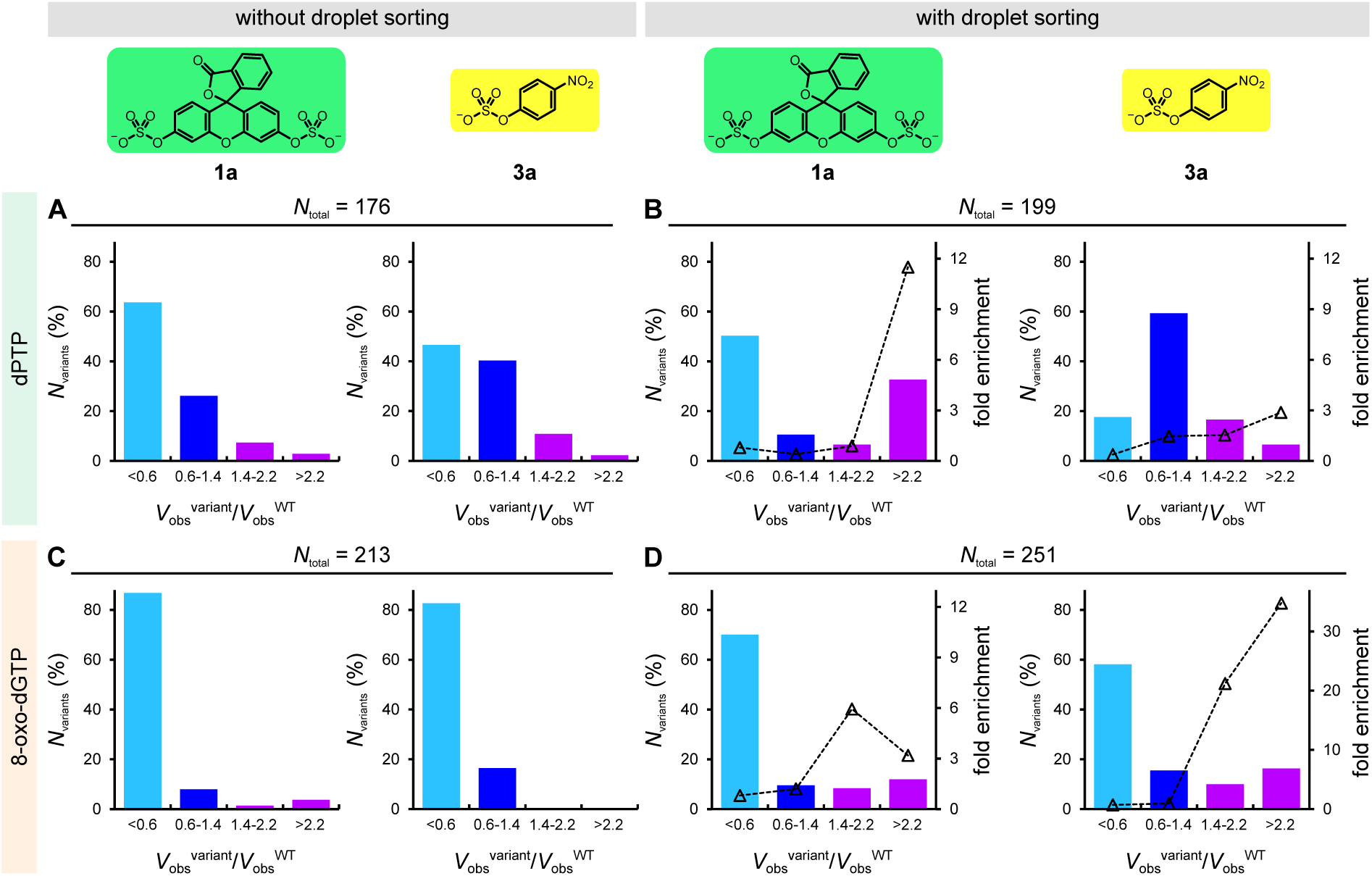
Assessment of the effect of screening for improved activity toward fluorescein disulfate (sulfate monoester **1a**), using fluorescence assisted droplet sorting (FADS), on the distribution of active *Sp*AS1-variants. For all libraries completely inactive variants were removed by only including *Sp*AS1 variants that showed activity toward sulfate monoester **2a** (screening step 2, Figure 3A and S6). We tested all libraries side-by-side for activity toward fluorescein disulfate **1a** and 4-nitrophenylsulfate **3a**. The activity data, normalized to wild-type controls recorded in the same experiment, are grouped into 4 categories, i.e. decreased activity (cyan bars), wild-type-like (blue), moderately improved and strongly improved (both purple). Comparison of the distributions with and without the inclusion of droplet sorting for improved activity toward fluorescein disulfate (**1a**) showed up to 40-fold enrichment of *Sp*AS1 variants with strongly improved activity toward either one of the two substrates. All variants showing >1.4-fold increased activity compared to wild-type for any of the two substrates (purple bars in panel **B** and **D**) were selected for the further screening in step 4 (see also Figure S9).

### Characterization of improved variants

All 27 unique *Sp*AS1 variants that contained at least 1 amino acid substitution were sub-cloned into a vector for cytosolic overexpression (see SI for details), produced in *E. coli* and purified to homogeneity. For all purified *Sp*AS1 variants the catalytic efficiency (*k*_cat_/*K*_M_) toward of fluorescein disulfate **1a** and 4-nitrophenyl sulfate **3a** was determined (Figure 6, Table S6 and S7). Out of the 27 purified *Sp*AS1 variants, 7 mutants showed improved catalytic efficiency only toward fluorescein disulfate **1a**, 7 mutants showed improved catalytic efficiency only toward 4-nitrophenyl sulfate **3a**, whereas 11 *Sp*AS1 variants showed improved catalytic performance toward both substrates (Figure S18). For two variants, the improvements in activity toward either substrate relative to *Sp*AS1^WT^ were within experimental error.

**Figure 6:**
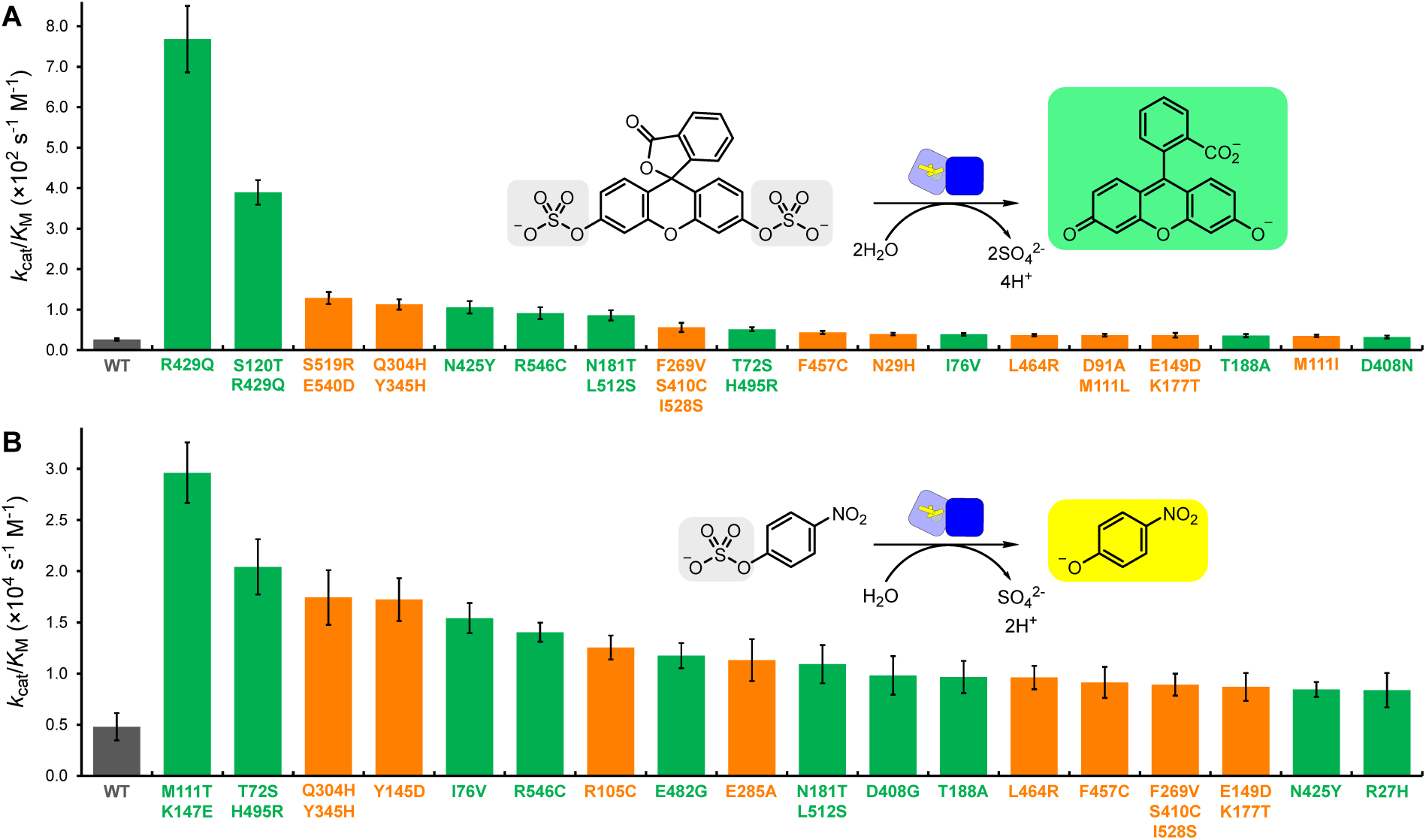
Catalytic efficiency (*k*_cat_/*K*_M_) toward fluorescein disulfate (**A**) and 4-nitrophenyl sulfate (**B**) of purified *Sp*AS1-variants. Mutants indicated in green and orange were generated using dPTP and 8-oxo-dGTP mutagenic nucleotides respectively. Only variants with significantly improved catalytic efficiency relative to wild type are depicted here. The individual kinetic parameters (*k*_cat_, *K*_M_, *K*_SI_ and *k*_cat_/*K*_M_) for all variants toward fluorescein disulfate (**1a**) and 4-nitrophenyl sulfate (**3a**) are listed in Tables S6 and S7 respectively.

The fusion of *Sp*AS1 variants to the autotransporter might affect their catalytic performance in such a way that the improvements in activity observed during step 4 of the screening procedure do not correlate accordingly with the improvements in catalytic efficiencies of the purified versions of the respective *Sp*AS1 variants. We obtained kinetic parameters for two substrates for 27 *Sp*AS1 variants, both in their autodisplayed and purified form (Figure S13-S16). Therefore, we are able to determine the correlation between the kinetic parameters of the autodisplayed and purified versions of the selected *Sp*AS1 variants (Figure S17). We observe positive and significant correlations between the second order rate constants of the autodisplayed (*V*_max_/*K*_M_) and purified (*k*_cat_/*K*_M_) *Sp*AS1 variants for both fluorescein disulfate **1a** and 4-nitrophenyl sulfate **3a** (Figure S17). Although the improvements do not translate directly, mostly likely due to differences in expression levels between the autodisplayed *Sp*AS1 variants, the improvement in *V*_max_/*K*_M_ for a given autodisplayed *Sp*AS1 variant is a good proxy for improvement of catalytic efficiency (*k*_cat_/*K*_M_) in the corresponding purified *Sp*AS1 variant.

In all 25 mutants that show significant improvement in catalytic efficiency toward fluorescein disulfate **1a** and/or 4-nitrophenyl sulfate **3a** we found in total 31 mutated positions. Out of these 31 positions, 6 were only found in mutants with improved activity toward fluorescein disulfate **1a** and 6 positions were exclusively associated with improvements in activity toward 4-nitrophenyl sulfate **3a**. Only 2 out of these 31 positions are conserved in a multiple sequence alignment of *Sp*AS1 with 94 closely related dimeric arylsulfatases,^37,43^ while all other positions are moderately to highly variable in the multiple sequence alignment (Figure S20). For the positions analogous to M111 in *Sp*AS1, the amino acid found in *Sp*AS1^WT^ is identical to the consensus residue. The conserved amino acid at the position analogous to Y345 is a histidine (Figure S20). Therefore, mutation Y345H is a so-called back-to-consensus mutation.^44^ Such mutations are likely to have a stabilizing effect.^45^

The top *Sp*AS1 variants for fluorescein disulfate **1a** (*Sp*AS1^R429Q^) and 4-nitrophenyl sulfate **3a** (*Sp*AS1^M111T/K147E^) both show no significant change in catalytic efficiency (*k*_cat_/*K*_M_) toward the respective other substrate (Table S6 and S7). The latter suggests that large improvements in catalytic efficiencies (*k*_cat_/*K*_M_) toward both substrates are difficult to achieve. Indeed, a significant negative correlation (*r* = −0.448; *p* = 1.93*×*10^-2^) between the activities toward fluorescein disulfate **1a** and 4-nitrophenyl sulfate **3a** (log *vs*. log) for all 27 selected *Sp*AS1 variants indicate that improved catalytic efficiency toward one substrate often comes at the cost of catalytic efficiency toward the other (Figure S18 and S19).

We found 18 *Sp*AS1 variants with 1.7-to 6.2-fold improved catalytic efficiencies toward 4-nitrophenyl sulfate **3a** (Figure 6B). The top 5 *Sp*AS1 variants for 4-nitrophenyl sulfate conversion (>3-fold improved *k*_cat_/*K*_M_) all contained at least one residue relatively close to the active site (within 8.2 Å). In particular mutations in T72 and I76, which could play a role in positioning conserved active site residue N74, and mutations observed in Y145, K147 and E149, which are all found in a loop region partially covering the active site entrance, could be classified as so-called ‘second shell mutations’ (Figure S21). The observation of larger effects on catalytic performance and/or specificity by mutations closer to the active site^46,47^ has been observed previously in many directed evolution campaigns involving screening of error-prone PCR-generated libraries. However, even the improvement in catalytic performance of the top performer for this substrate, *Sp*AS1^M111T/K147E^, does not exceed an order of magnitude, i.e. the improvements in performance toward 4-nitrophenyl sulfate **3a** are fairly modest.

In total 18 *Sp*AS1 variants with improved activity toward fluorescein disulfate **1a** were found. The improvements in performance relative to wild type were considerably larger than for 4-nitrophenyl sulfate **3a**: up to 30-fold improved catalytic performance was observed. The larger improvement found is most likely due to the fact that the first screening step is expected to enrich for variants with improved performance toward fluorescein disulfate **1a**. Many of the mutations found in the *Sp*AS1 variants with improved catalytic performance toward fluorescein disulfate **1a** are located in the largely α-helical C-terminal region involved in the formation of the oligomeric interface of the *Sp*AS1 dimer (amino acid number >420 aa). Four out of the five residues that are mutated in this region appear to form hydrogen bonds with the side-chain of a residue in another α-helix, either within the same protomer (*E518*-R546 and S519-E540) or between α-helices from different protomers ((R429-*D513*, Figure 7B). Mutations in the S519-E540 pair even occur within the same *Sp*AS1 variant. For all three pairs of residues, the observed mutations still allow the formation of a hydrogen bond (R429Q, S519R, E540D and R546C), possibly with the same partner as in the wild-type. However, the distance and/or angle of the ‘new’ hydrogen bond donor-acceptor pairs will no longer be optimal. The latter could result either in *i*) loss of that particular hydrogen bond interaction (and possible formation of another hydrogen bond with a different residue), or *ii*) movement of the respective α-helices in order to re-optimize hydrogen bonding. Both effects are expected to alter the shape of the putative substrate binding pocket and as a result improve the catalytic performance toward fluorescein disulfate **1a**. Furthermore, for all three hydrogen bonding pairs that apparently influence substrate binding when altered, one of the contributing residues of each hydrogen bonding pair is located in the penultimate α-helix of *Sp*AS1. The latter α-helix makes direct contact with the other protomer, suggesting a critical role of the oligomerization interface in determining substrate specificity.

**Figure 7:**
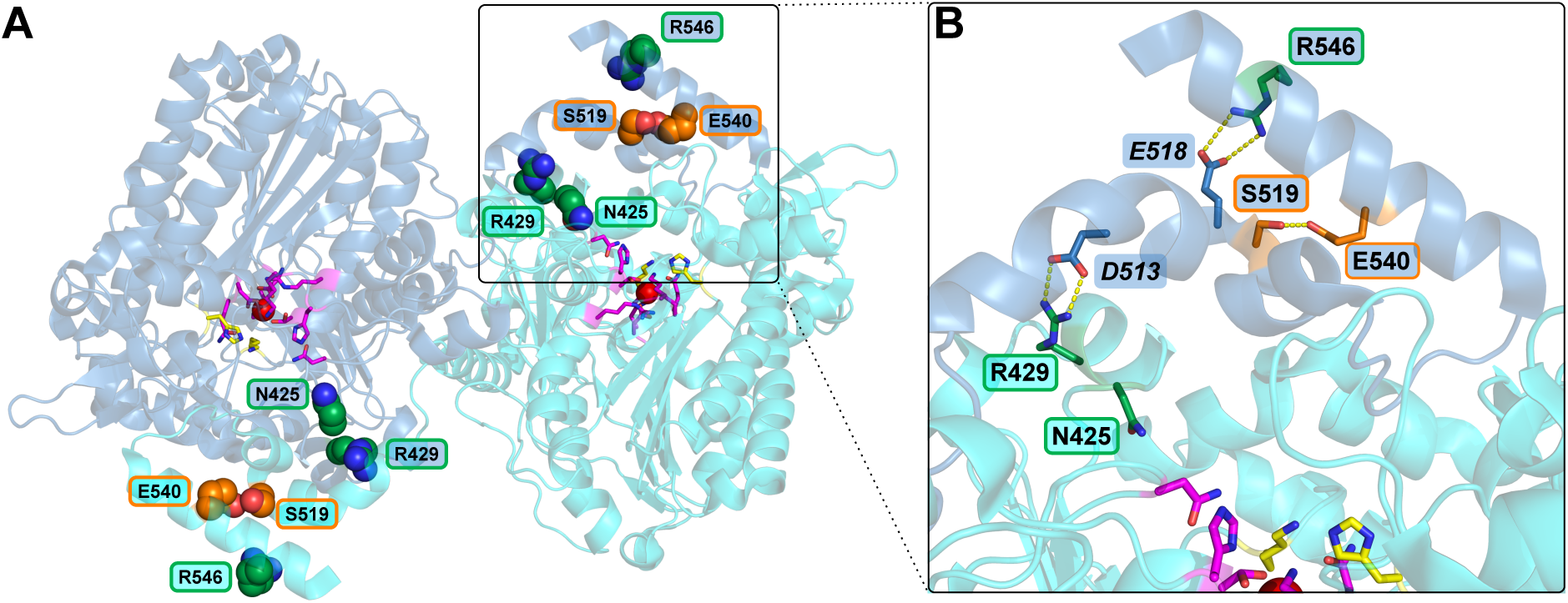
Location of the residues in the C-terminal region of dimeric *Sp*AS1 in which mutations have a >3.5-fold positive effect of the catalytic activity toward fluorescein disulfate **1a** (PDB ID for monomer: 4UPI). Mutations indicated in green and orange were introduced during mutagenic PCRs with nucleotide analogs dPTP and 8-oxo-dGTP respectively. The conserved active site groups^37,40^ are indicated in yellow (putative leaving group stabilizing residues H234 and K341), magenta (all other putative conserved active site residues), and red (active site metal ion). (**A**) Positions of the residues that are mutated in the C-terminal region of *Sp*AS1 that are involved in the formation of the oligomerization interface. (**B**) Enlarged view of the region indicated in panel A. All but one (N425) of the five residues in which mutations were found appear to form hydrogen bonds with the side-chain of a residue in another α-helix, either within the same protomer (*E518*-R546 and S519-E540) or between α-helices from different protomers, which are part of the oligomeric interface (R429-*D513*). Mutations in the S519-E540 pair occur within the same *Sp*AS1 variant.

### Advantages of bacterial autodisplay

Robustness and throughput of the screening procedure are most important for the success of a directed evolution campaign. Throughput determines the likelihood of finding an improved variant and thereby determines the time and effort needed to accomplish a certain degree of improvement. Medium to high throughput detection of sulfatase activity has been accomplished previously, using either conventional agarplate-based and microtiterplate-based methods only^48^ or a combination thereof with microdroplets.^21,22,25^ For all previous campaigns the sulfatase variants were expressed in the cytosol of *E. coli*. Since negatively charged sulfate esters cannot pass the negatively charged head groups of the phospholipids of the cell membranes, cell lysis was required prior to activity measurements.^21,22,25,48^ As a control, we showed that intact *E. coli* cells overexpressing *Sp*AS1 wild-type in their cytosol indeed do not show detectable sulfatase activity.

Displaying a library of enzyme variants on the surface of a cell has several advantages over cell lysis: *i*) it is possible to avoid interfering endogenous activities of cytosolic enzymes present in the host *ii*) in essence only one liquid handling step (mixing of cells and substrate) is required *vs*. at least three steps when using cell lysis (see SI for details) and, *iii*) in particular for microdroplet-based screening, selected genotypes can be directly recovered by cell growth of selected clones, instead of the alternative, recovery of genetic diversity by purification and subsequent re-transformation of plasmid DNA, which is required when cells are lysed during the droplet screening procedure.

The fraction of genetic diversity that can be maximally recovered by plasmid re-transformation is dependent on *i*) the copy number of the plasmid, i.e. how many plasmid molecules can be maximally recovered from the single cell lysate and *ii*) how efficiently the plasmid is retransformed, i.e. which fraction of the isolated plasmids are successfully transformed and result in the formation of a colony. For small, high copy-number plasmids (∼ 300 plasmids/cell) that transform efficiently (1 in 18 plasmid molecules is successfully transformed), the genotypic diversity can be recovered multiple times (maximum ∼ 1670%, see SI for details). However, for larger, low copy-number plasmids (∼ 20 copies/cell) that transform less efficiently (1 in 3661), such as the pBAD-AT-His_6_-*Sp*AS1 plasmid we are using here, the maximum recovery of genetic diversity is 0.55% (see SI for details). The live cell re-growth method we used recovered 12% of the sorted diversity, thereby outperforming re-transformation at least >20-fold (see SI for details).

Another advantage, compared e.g. to yeast display, is that our autodisplay system can be used to display libraries of homodimeric enzymes. In yeast display a single protein molecule of interest is linked to either agglutinin or floculation domains which are in turn (covalently) linked to the rigid yeast cell wall.^49^ This means that the displayed protein cannot move freely and form homodimers. The autotransporter we fused our libraries of *Sp*AS1-variants to can move through the outer membrane of *E. coli*, allowing individual *Sp*AS1 monomers to interact and form homodimers. Furthermore, the use of bacterial autodisplay can be easily implemented to replace any lysis-based screening system for which the widely-used laboratory work horse *E. coli* was previously used, all without drastically changing standard experimental procedures for cell growth and genetic manipulation.

## Concluding Remarks

In this study we show, for the first time, the combination of microdroplet-based single variant screening with *E. coli* autodisplay. Using this system we quantitatively screened 10^5^-10^6^ *Sp*AS1 variants for improved sulfatase activity toward fluorescein disulfate within several hours. The mobile β-barrel anchor of the autotransporter system facilitated screening a library of random variants of the *homodimeric* sulfatase *Sp*AS1 we used as a model system. Such a screen is currently not possible with yeast display, due to the immobility of the ‘anchors’ that are attached to the displayed protein subunits. Using living *E. coli* cells during the screening steps enabled us to *i*) recover the genotypic diversity after droplet sorting >20-fold more efficient compared to re-transformation-based recovery and *ii*) test for sulfatase activity in microdroplets without cell lysis. Avoiding cell lysis before activity measurements also simplified all subsequent screening steps.

The lysis-free 4-step screening procedure ultimately resulted in the identification of 25 unique *Sp*AS1 variants with up to 30-fold improved catalytic performance toward fluorescein disulfate and up-to 6.2-fold improved activity toward 4-nitrophenyl sulfate after a single round of mutagenesis. All 31 positions in which mutations were found were non-obvious, i.e. none were previously described conserved active site amino acids.^37,40^ In particular mutations in residues that form hydrogen bonds between various α-helices in the C-terminal oligomerization region appear to be important for the accommodation of the large fluorescein disulfate substrate. The latter region is highly variable between the dimeric sulfatases that *Sp*AS1 is closely related to.^37,43^ This high sequence variability would make (semi)-rational prediction of beneficial mutations in this C-terminal region especially tricky, which means the ability to screen large random mutagenesis-based libraries of enzyme variants continues to be essential to reach large improvements in enzyme performance. Therefore, the development of high-throughput screening methods such as the one we describe in this paper is essential for successful enzyme improvements.

## Methods

### Construction of plasmid vectors for *E. coli* autodisplay of *Sp*AS1

The previously described dimeric arylsulfatase 1 from *Silicibacter pomeroyi* DSS-3^37,40^ (*Sp*AS1) was cloned into autodisplay vector pBAD-AT^42^ using standard restriction endonuclease-based cloning using restriction sites *Xho*I and *Kpn*I. Exact experimental details regarding PCR amplification and molecular cloning can be found in the expanded methods.

Initial testing if *Sp*AS1 was displayed as active enzyme was done by testing *E. coli* cells expressing the His_6_-*Sp*AS1-autotransporter construct for the ability to catalyze the hydrolysis of 4-nitrophenyl sulfate (sulfate monoester **3a**). As a negative control we used a His_6_-*Sp*AS1-autotransporter construct of an inactive *Sp*AS1 variant (C53A, *k*_cat_/*K*_M_ ∼ 10^5^-fold below wild type). Bacterial culture conditions for expressing the His_6_-*Sp*AS1-autotransporter construct are described in detail in the expanded methods. Additional assessment of correct autodisplay of *Sp*AS1 was done essentially as described previously.^50,51^ In short, we expressed the His_6_-*Sp*AS1-autotransporter construct at 80 mL scale. Half of the cells were treated with proteinase K while the other half was untreated. We subsequently isolated all outer membrane proteins for both treatments and analyzed the protein extracts using SDS-PAGE. Details regarding the bacterial culture conditions, proteinase K treatment and total membrane protein isolation are described in the expanded methods.

### Generation of mutant libraries

Genotypic diversity in the *Sp*AS1 libraries was created by two separate error-prone PCR reactions using nucleotide analogs and 2’-deoxy-P-nucleoside-5’-triphosphate (dPTP) and 8-oxo-2’-deoxyguanosine-5’-triphosphate (8-oxo-dGTP) respectively,^52^ both in combination with the non-proofreading *Taq* DNA polymerase (Go*Taq*, Promega). The primers used for this PCR reaction anneal outside the His_6_-*Sp*AS1 open reading frame (Table S8). The resulting PCR-products were purified and used as a template in an non-mutagenic PCR in order to remove any nucleotide analogs incorporated in the DNA. The non-mutagenic PCR was done essentially as described above, except in this case we used a proofreading polymerase (Herculase II) and only conventional nucleotides (dTNPs). Cloning of the mutant library into the pBAD-AT vector was essentially done as described above. For each library we sequenced 12 randomly picked library variants to asses the mutation frequencies, which were 2.3±1.7 (dPTP) and 3.7±2.6 (8-oxo-dGTP) non-synonymous base pair substitutions per gene respectively (Table S1).

### Library screening procedures

The two different error-prone libraries generated as described above were screened for *Sp*AS1 variants with improved sulfatase activity in four subsequent steps. For all these steps *Sp*AS1 was displayed on the outer membrane of *E. cloni* 10G. During the first step we screened the displayed *Sp*AS1 library for improved activity toward fluorescein disulfate **1a** using a fluorescence activated droplet sorter (FADS, Figure 2 and S3-S4). The cells displaying *Sp*AS1 variants with improved sulfatase activity were regrown to fully sized colonies on a nitrocellulose filter sitting on top of solid medium. For the second step these colonies were tested for improved activity toward 5-bromo-4-chloro-3-indolyl sulfate (X-sulfate **2a**, Figure 3A and S6). All variants that turned visibly blue within 30 minutes after exposure to X-sulfate **2a** were scored as positive. In the third step these positive clones were tested for improved turnover rates of fluorescein disulfate **1a** and 4-nitrophenyl sulfate **3a** at sub-saturating substrate concentrations (i.e. at [S]≪*K*_M_) (Figure 3B and S7). Displayed *Sp*AS1 variants that showed >1.4-fold improved activity toward either of the two substrates (compared to wild type) were chosen for further testing. In the fourth and final step we approximated Michaelis-Menten parameters *V*_max_, *K*_M_ and *V*_max_/*K*_M_ for the displayed *Sp*AS1 variants for both fluorescein disulfate **1a** and 4-nitrophenyl sulfate **3a** (see Figure 3C and S8 and expanded methods for details). Mutants that showed significant improvement to either substrate relative to wild-type in a side-by-side comparison, were chosen for further characterization.

### Characterization of *Sp*AS1 variants

All 27 unique autodisplayed *Sp*AS1 variants selected after the fourth screening step that contain at least one *non*-synonymous mutation were selected for more detailed characterization. The corresponding *Sp*AS1 variant-encoding genes were cloned into the pASKIBA5^+^ vector. The resulting N-terminally strep-tagged *Sp*AS1 variants were produced in *E. coli* TOP10 and purified to homogeneity. Detailed experimental procedures for cloning, protein expression, purification and the determination of kinetic parameters (*k*_cat_, *K*_M_, *K*_SI_ and *k*_cat_/*K*_M_) toward fluorescein disulfate **1a** and 4-nitrophenyl sulfate **3a** are described in the expanded methods.

## Associated content

Supporting information containing further details about the used methods and microfluidic devices, additional discussion, and all kinetic data is available.

## Supporting information

Supportinginformation

## Acknowledgements

This research was funded by the Human Frontier Science Program (to F.H. and E.B.B.; grant number RGP0006/2013). P.M. holds a studentship from the Engineering and Physical Sciences Research Council (EP/L015889/1). A.Z. was supported by the BBSRC, the Cambridge Home and EU Scholarship Scheme (CHESS) and the EU Marie-Curie networks PhosChemRec (FP7-PEOPLE ITN-2009-238679) and ENEFP (FP7-PEOPLE-2007-1-1-ITN-215560). J.S. was funded by a fellowship of the Federal Ministry of Education and Research as part of a project within “Bioindustrie 2021” (funding reference: 0316163B). B.D.G.E holds a fellowship from the European Community’s Innovative Training Network ES-cat (722610), J.M.H. is supported by the EU (metafluidics, 685474). F.H. is an ERC Advanced Investigator (695669).

