## Supportinginformation for "High-Throughput, Lysis-free Screening for Sulfatase Activity Using *Escherichia coli* Autodisplay in Microdroplets"

#### Contents

**1 Expanded methods** . . . **S3**
1.1 Materials . . . S3
1.1.1 Reagents . . . S3
1.1.2 Microfluidic chips . . . S3
1.2 Construction of plasmid vectors . . . S3
1.2.1 *E. coli* autodisplay: pBAD-AT-$His_6$-*Sp*AS1 . . . S3
1.2.2 Protein production for purification: pASKIBA5$^+$-*Sp*AS1 . . . S3
1.3 Verification of *E. coli* autodisplay of *Sp*AS1 . . . S4
1.3.1 Activity testing . . . S4
1.3.2 Isolation and analysis of total membrane protein . . . S4
1.4 Error-prone PCR . . . S5
1.5 Screening step 1: microdroplets . . . S5
1.5.1 Preparation of *E. coli* cells for microdroplet encapsulation. . . . S5
1.5.2 Generation of droplets . . . S5
1.5.3 Fluorescence activated droplet sorting (FADS) . . . S6
1.6 Screening step 2: agar plate activity screening . . . S6
1.7 Screening step 3: microtiter plate activity screening . . . S7
1.7.1 Expression of autodisplayed protein in microtiterplates . . . S7
1.7.2 Activity testing: fluorescein disulfate . . . S7
1.7.3 Activity testing: 4-nitrophenyl sulfate . . . S8
1.8 Screening step 4: approximation of catalytic efficiency measurements with whole cells . . . S8
1.9 Protein production and purification . . . S10
1.10 Measurement of kinetic parameters for purified *Sp*AS1 variants . . . S10
1.11 Calculation of the expected rate constant of uncatalyzed conversion of fluorescein disulfate **1a** . . . S12
1.12 Conversion of transformation efficiencies to suitable units for comparison between plasmids of different sizes . . . S12

2 **Expanded discussion** **S14**
2.1 Comparison cell lysis *vs.* autodisplay . . . . . . . . . . . . . . . . . . . . . . . . . S14
2.1.1 Recovery of genetic diversity after microdroplet sorting . . . . . . . . . . . S14
2.1.2 Recovery and number of experimental steps during agar plate-based screening . . . . . . . . . . . . . . . . . . . . . . . . . . . . . . . . . . . . S15
2.1.3 Number of experimental steps during microtiterplate screening . . . . . . S15

3 **Supporting Figures** **S17**

4 **Supporting Tables** **S39**

5 **Supplementary references** **S47**

### 1 Expanded methods

#### 1.1 Materials

##### 1.1.1 Reagents

Sulfate monoester **1a** was synthesized as described previously.[1] Sulfate monoester **2a** was from Gold Biotechnology. Sulfate monoester **3a** was from Acros. Autodisplay vector pBAD-AT was constructed as described previously.[2] All restriction enzymes and T4 DNA ligase were from Thermo Scientific. Overexpression vector pASKIBA5$^+$ and Strep-tactin resin were from IBA. Herculase II was from Agilent. Nucleotide analogs 8-oxo-dGTP (8-Oxo-2'-deoxyguanosine-5'-Triphosphate) and dPTP (2'-Deoxy-P-nucleoside-5'-Triphosphate) were from Jena Biosciences. Go*Taq* DNA Polymerase was from Promega.

##### 1.1.2 Microfluidic chips

The microfluidic devices used for droplet generation and droplet sorting were fabricated following classical soft-lithography procedures by using high-resolution acetate masks (Microlithography Services Ltd) and SU-8 photoresist patterning. The patterned poly(dimethyl siloxane) (PDMS, Dow Corning) was plasma bonded to a microscope glass slide in a low-pressure oxygen plasma generator (Femto, Diener Electronics), flushed with 1% (v/v) trichloro(1*H*,1*H*,2*H*,2*H*-perfluorooctyl)silane (Sigma-Aldrich) in HFE-7500 (3M), and left at 65 °C for 30 minutes.

#### 1.2 Construction of plasmid vectors

##### 1.2.1 *E. coli* autodisplay: pBAD-AT-His$_6$-*Sp*AS1

The *Sp*AS1-encoding gene was amplified by PCR using the previously described pASKIBA5$^+$-*Sp*AS1$^{WT}$ construct[3,4] as a template (10-20 ng plasmid DNA). Forward and reverse primers (Table S8) were used at 0.4 µM in a reaction with 0.2 mM dNTPs and Herculase II DNA polymerase (Agilent). The N-terminal primer includes a poly-histidine tag (His$_6$). The temperature cycling program consisted of 5 minutes at 95 °C, followed by 30 cycles of 1 minute at 95 °C, 45 seconds at 68 °C - 0.5 °C cycle$^{-1}$ (each cycle the temperature of this segment was lowered by 0.5 °C), 3 minutes at 72 °C, and finished with 10 min at 72 °C. The PCR products were digested with *Xho*I and *Kpn*I and subsequently ligated into *Xho*I-*Kpn*I-digested pBAD-AT[2] plasmid DNA using T4 DNA ligase. The ligation mixture was transformed to *E. coli* (*E. cloni* 10G, Lucigen). Cells were plated on LB-agar containing ampicillin (50-100 mg L$^{-1}$) and resulting colonies were checked for insert using a PCR reaction with Go*Taq* DNA polymerase and colony material as the template. Plasmid DNA was extracted from positive clones and the correct insertion of the *Sp*AS1 gene was confirmed by sequencing.

##### 1.2.2 Protein production for purification: pASKIBA5$^+$-*Sp*AS1

PCR amplification of improved *Sp*AS1-variants for subcloning into the pASKIBA5$^+$ vector for cytosolic overexpression was done essentially as described above using the appropriate cloning primers (see Table S8 for details) to introduce the translational fusion to the strep-tactin tag

(N-terminus) and re-introduction of the stop-codon (C-terminus). Mutant *Sp*AS1$^{R546C}$ required the use of an alternative reverse cloning primer, since using the standard primer would revert the R546C mutant to wild type. The resulting PCR product was ligated into *Xho*I-*Pst*I digested pASKIBA5$^{+}$ plasmid DNA using T4 DNA ligase. The ligation mixture was transformed into *E. coli* (*E. coli* TOP10, Invitrogen). Cells were plated on LB-agar containing ampicillin (50-100 mg $L^{-1}$) and resulting colonies were checked for insert using a PCR reaction with Go*Taq* DNA polymerase and colony material as the template. Plasmid DNA was extracted from positive clones and the correct insertion of the *Sp*AS1 variant was confirmed by DNA sequencing.

#### 1.3 Verification of *E. coli* autodisplay of *Sp*AS1

##### 1.3.1 Activity testing

*E. coli* cells containing the pBAD-AT-His$_6$-*Sp*AS1$^{WT}$ plasmid construct were grown at 37 °C in 3 mL LB-medium containing ampicillin (50-100 mg $L^{-1}$) until clearly turbid ($OD_{600}$ ∼0.5-0.8), at which point the growth temperature was lowered to 25 °C and cells were left to grow for an hour. Expression of the autodisplay construct was subsequently induced by the addition of up to 0.02% (v/v) L-arabinose followed by overnight growth at 25 °C. For activity testing the cells were mixed with 1 mM 4-nitrophenylsulfate **3a** and the appearance of nitrophenolate **3b** resulted in an increase of absorbance at $\lambda$ = 400 nm ($A_{400}$) over time when the *Sp*AS1 variant was active. An autodisplay construct of *Sp*AS1$^{C53A}$, a variant of *Sp*AS1 with ∼$10^5$-fold lowered $k_{cat}/K_M$, was used as a negative control.

##### 1.3.2 Isolation and analysis of total membrane protein

Expression of autodisplayed *Sp*AS1 was performed with a larger culture volume (80 mL), but otherwise the growth conditions were as described above for the activity testing. After overnight expression of the autodisplayed *Sp*AS1 construct the culture was split into two equal portions. For both samples the cells were harvested by centrifugation (3850×g, 10 minutes). One of the two cell pellets was resuspended in 1 mL PBS buffer (10 mM $Na_2HPO_4$; 2 mM $KH_2PO_4$; 137 mM NaCl; 2.7 mM KCl, pH 7.4) and up to 62 µg $mL^{-1}$ proteinase K was added. The mixture was incubated at 37 °C for one hour and the reaction was stopped by adding 5 mL 200 mM Tris-HCl pH 8.0 containing 10% fetal calf serum (FCS). The mixture was centrifuged (3850 × *g*, 10 minutes) and resulting cell pellet was used for further experiments.

Both the proteinase K-treated and untreated cell pellets were resuspended in 5 mL 60 mM Tris-HCl pH 8.0 containing 20 mM sucrose, 0.2 mM EDTA and 0.2 mg $mL^{-1}$ lysozyme and incubated at room temperature for 10 minutes. Both samples were mixed in a 1:1 ratio with 1 mM phenylmethylsulfonyl fluoride (PMSF), 20 µg $mL^{-1}$ aprotinin and 10 µg $mL^{-1}$ DNAse in extraction buffer (2% (v/v) Triton X-100, 50 mM Tris-HCl pH 8.0, 10 mM $MgCl_2$) and incubated on ice for 30 minutes. The samples were centrifuged for 5 minutes at 3850 × *g*. The resulting supernatant was subsequently centrifuged at high speed (10 minutes, 38700 × *g*, 4 °C). The resulting pellets, which contain all membrane-bound proteins, were washed twice with ultrapure water, resuspended in ultrapure water and analyzed using SDS-PAGE.

#### 1.4 Error-prone PCR

The two mutagenic PCRs, using 8-oxo-2'-deoxyguanosine-5'-triphosphate (8-oxo-dGTP) and 2'-deoxy-P-nucleoside-5'-triphosphate (dPTP) (Jena Biosciences) respectively to introduce mutations, were performed using 30-35 ng pBAD-AT-His$_6$-*Sp*AS1 plasmid DNA as a template, 0.5 µM forward and reverse primers that anneal outside the *Sp*AS1 coding sequence in the pBAD-AT-His$_6$-*Sp*AS1 construct (Table S8), 0.25 mM of each dNTP, 2 mM $MgCl_2$ and Go*Taq* DNA Polymerase (Promega). Additionally. The mutagenic oligonucleotides were used at 40 µM (8-oxo-dGTP) and 1.0 µM (dPTP) respectively. The temperature cycling program consisted of 10 minutes at 95 °C, followed by 13 cycles of 1 minute at 95 °C, 1 minute 45 seconds at 68 °C - 1.0 °C cycle$^{-1}$ (each cycle the temperature of this segment was lowered by 1.0 °C), 1 minute 45 seconds at 72 °C, followed by 19 cycles of 1 minute at 95 °C, 45 seconds at 56 °C, 1 minute 45 seconds at 72 °and finished with 10 min at 72 °C. In order to remove the wild-type template material after thermal cycling, we added 20 units of *Dpn*I (Thermoscientific) to the reaction mixture and incubated at 37 °C for 2 hours. The reaction mixtures were loaded onto an 0.8 % (w/v) agarose gel and the fragment of the correct size was isolated and purified using standard techniques. In order to remove the incorporated mutagenic bases, 30-35 ng of the purified error-prone PCR product was amplified in a regular PCR as described above.

#### 1.5 Screening step 1: microdroplets

A graphical representation of the microdroplet screening procedure is depicted in Figure S3.

##### 1.5.1 Preparation of *E. coli* cells for microdroplet encapsulation.

We plated $\sim 10^5$-$10^6$ autodisplayed *Sp*AS1-variants propagating in *E. cloni* 10G onto LB-agar containing 50-100 mg L$^{-1}$ ampicillin followed by overnight growth at 37 °C. The resulting colonies were resuspended into LB medium containing 15% (v/v) glycerol. The $OD_{600}$ was measured and the cell suspension was divided into aliquots each containing $\sim 10^{10}$ cells (assuming $OD_{600}$ = 1 to correspond to $5 \times 10^8$ *E. coli* cells mL$^{-1}$). These aliquots could be stored at -80 °C or used immediately. Expression of the resuspended *Sp*AS1 mutant library was induced by diluting a single $10^{10}$-cell aliquot into a total volume of 5 mL LB medium containing ampicillin (100 mg L$^{-1}$) and L-arabinose (0.2% (v/v)). After overnight expression of the *Sp*AS1-library at 20-25 °C/200 rpm, the cells were harvested by centrifugation and washed three times in SID buffer (44 mM Succinic acid, 33 mM Imidazole, 33 mM Diethanolamine) pH 7.5 containing 1% (v/v) glycerol. The washed cells were resuspended in 1 mL SID buffer pH 7.5 containing 1% (v/v) glycerol and the $OD_{600}$ was determined. The cell suspension was diluted down to $4 \times 10^8$ cells mL$^{-1}$ ($OD_{600}$ = 0.8) in SID buffer pH 7.5 containing 1% (v/v) glycerol prior to droplet generation.

##### 1.5.2 Generation of droplets

Droplets containing *E. cloni* cells expressing the autodisplayed *Sp*AS1-library and 5 µM fluorescein disulfate **1a** were produced in a double flow-focusing junction chip. At the first junction, the cell suspension was mixed 1:1 with a solution of 10 µM fluorescein disulfate **1a** in SID buffer pH 7.5. At the second junction, the mixed aqueous phase was dispersed into the fluorous oil HFE-7500 (3M) containing 1% (v/v) fluorosurfactant-008 (RAN Biotechnologies) generating 20 µm droplets with an expected cell occupancy of λ = 0.35 (assuming Poisson distribution). The solutions were injected using syringe pumps (Nemesys) and gas-tight syringes which were connected to the chip *via* PTFE tubing; the flow rates were 50 µL min$^{-1}$ for both aqueous phases and 500 µL min$^{-1}$ for the oil/surfactant phase. The droplet formation was monitored on an inverted microscope (SP981, Brunell Microscopes) equipped with a high-speed camera (Miro ex4, Phantom Research).The droplets were collected and stored overnight in a dark place at room temperature.

##### 1.5.3 Fluorescence activated droplet sorting (FADS)

Droplets were sorted according to their fluorescence intensity using a custom-built fluorescence activated droplet sorter (FADS), based on previously used droplet sorter designs.[5] Droplets were injected into a sorting chip at 5-15 µL min$^{-1}$ and spaced out with oil (HFE7500, 3M) at 150 µL min$^{-1}$, resulting in a sorting frequency of 1.0-1.5 kHz. Fluorescence of fluorescein **1b** was measured by excitation at λ = 488 nm (laser 85-BCD-30, Melles-Griot) and detecting emission at λ = 497-553 nm using a photo mutliplier tube (PMT, PM002, Thorlabs). Sorting was performed using a field-programmable gate array (FPGA, PCIe-7841R, National Instruments) which monitored and recorded the signal using custom LabView software. If a droplet fulfilled the operator-defined sorting criteria, the FPGA triggered a pulse generator (PG, TGP110, Thurlby Thandar Instruments) and image acquisition by the high-speed camera (Miro Ex4, Phantom Research). The electronic pulse was amplified to 600V and applied to the electrode channels filled with a 4M NaCl solution. The entire process was monitored on an inverted microscope (IX73, Olympus). See Figure S4 for details regarding the set-up of the droplet sorter.

Positive droplets were collected in a microvial containing 45 µL HFE-7500, 5 µL per-fluorooctanol and 100 µL Lucigen recovery medium. After the sorting was finished the final sorted droplets were washed out of the tubing with HFE-7500 and 500 µL lucigen recovery medium was added. The medium phase was supplemented with up to 0.5% (w/v) sodium pyruvate and the entire mixture was incubated at 37 °C under mild agitation for two hours. The medium phase was plated onto nitrocellulose filters sitting on top of LB-agar containing ampicillin (100 mg L$^{-1}$) and the plates were incubated overnight at 37 °C.

#### 1.6 Screening step 2: agar plate activity screening

After overnight growth at 37 °C, the nitrocellulose filters containing *E. cloni* cells containing pBAD-AT-His$_6$-*Sp*AS1 variants were transfered to LB-agar plates containing 100 mg L$^{-1}$ ampicillin and 0.02% (w/v) L-arabinose to induce expression of the autodisplay construct. These plates were left to incubate overnight at room temperature. The following day the nitrocellulose filter was transferred to LB-agar containing 100 mg L$^{-1}$ ampicillin, 0.02% (w/v) L-arabinose and 0.005% (w/v) 5-bromo-4-chloro-3-indolyl sulfate (sulfate monoester **2a**). Enzyme-catalyzed hydrolysis of sulfate monoester **2a** will ultimately result in the formation of a blue dye (chromophore **2b**). Each colony that turned blue within 30 minutes after transfer of the nitrocellulose filter to the sulfate monoester **2a**-containing LB-agar plates was used to inoculate a 200 µL microtiter plate-well with LB-medium containing 10% (v/v) glycerol and 100 mg $L^{-1}$ ampicillin. After overnight growth at 37 °C, the resulting microtiter plates were kept at -80 °C for reference and storage. Wells with pBAD-His$_6$-*Sp*AS1$^{WT}$ (for assessment of the catalytic performance of the library variants compared to the wild-type) and pBAD-His$_6$-*Sp*AS1$^{C53A}$ (inactive variant, negative control) were included in quadruplicate on each microtiter plate. A graphical representation of the entire agar plate-based screening step is depicted in Figure S6.

#### 1.7 Screening step 3: microtiter plate activity screening

A graphical representation of the entire microtiter plate-based screening step is depicted in Figure S7. All optical measurements for this step (absorbance and fluorescence) were carried out using a TECAN infinite M200 microtiter plate reader.

##### 1.7.1 Expression of autodisplayed protein in microtiterplates

The reference microtiter plates containing the active *Sp*AS1 variants from the agar plate-based screening (screening step 2) were replicated into microtiter plates with 100 µL LB-medium containing 100 mg $L^{-1}$ ampicillin in each well. The plates were subsequently incubated at 37 °C/180 rpm for 6-8 hours, until each well had become clearly turbid. The temperature was lowered to 25 °C/180 rpm and the cells were left to grow for another hour. At this point we added 100 µL LB-medium containing 100 mg $L^{-1}$ ampicillin and 0.04% (w/v) L-arabinose (final concentration = 0.02% (v/v)) to each well and the plates were incubated overnight at 25 °C/180 rpm. The optical density ($OD_{600}$) for each well was determined in order to correct for variations in cell density. For each well the initial rate of substrate conversion was performed for both fluorescein disulfate **1a** and 4-nitrophenyl sulfate **3a**.

##### 1.7.2 Activity testing: fluorescein disulfate

Activity testing was done by monitoring the formation of fluorescein **1b** as a result of whole-cell-catalyzed hydrolysis of fluorescein disulfate **1a** over time. The increase in concentration of fluorescein **1b** over time was determined by measuring fluorescence ($\lambda_{ex}$ = 490 nm; $\lambda_{em}$ = 515 nm) in black opaque microtiterplates. Activity tests were carried out at 0.5 mM fluorescein disulfate **3a**, $\sim$10-fold below the $K_M$ of *Sp*AS1$^{WT}$ toward fluorescein disulfate **1a** to ensure that the observed improvements were representative for improvements of $k_{cat}/K_M$. Calibration measurements with fluorescein in buffer containing varying amounts of *E. cloni* 10G cells expressing autodisplayed *Sp*AS1$^{WT}$ showed that the strength of the fluorescence response ($F_{response}$ in $F_{counts}$ $M^{-1}$) decreases with increasing cell density and is linear within the range of optical densities used (Figure S22). Therefore, the initial rates ($V_{init}$) in $F_{counts}$ $s^{-1}$ were converted to M $s^{-1}$ with the correctly adjusted $F_{response}$ for each individual well based on the measured $OD_{600}$ (using the the calibration line in Figure S22), according to equation 1.

$$V_{\text{init}} = \frac{F_{\text{counts}}\ \text{s}^{-1}}{F_{\text{response}}} \qquad (1)$$

In addition we corrected all measured rates for chemical hydrolysis of fluorescein disulfate **1a** ($V_{\text{chem}}$), the latter calculated according to equation 2, resulting in $V_{\text{corr}}$ in M s$^{-1}$ as calculated according to equation 3

$$V_{\text{chem}} = k_{\text{uncat}} \times [\text{S}] \tag{2}$$

$$V_{\text{corr}} = V_{\text{init}} - V_{\text{chem}} \tag{3}$$

In which $k_{\text{uncat}}$ = 2.4×10$^{-8}$ s$^{-1}$, as calculated from the published temperature dependence data for a range of arylsulfates[6] and assuming a p$K_a$ for the fluorescein leaving group of 6.4 (Figure S22B). Measurement of the background rate of fluorescein disulfate conversion confirmed the expected value to be correct. The corrected reaction rates ($V_{\text{corr}}$ in M s$^{-1}$) were normalized for the cell density ($OD_{600}$) for each individual well. The activity measurements of the wild-type typically showed a variation of ±30%. Therefore only variants with a >1.4-fold increase in activity relative to wild-type were selected for the next screening round.

##### 1.7.3 Activity testing: 4-nitrophenyl sulfate

Activity testing was done by monitoring the formation of 4-nitrophenolate **3b** as a result of whole-cell-catalyzed hydrolysis of 4-nitrophenyl sulfate **3a** over time. The increase in concentration of 4-nitrophenolate **3b** over time was determined by measuring the absorbance at 400 nm. We mixed 50 µL SID buffer pH 7.5 and 70 µL cell suspension from each well and measured the cell density ($OD_{600}$) of each individual well. Activity tests were carried out at 1 mM 4-nitrophenyl sulfate **3a**, ~8-fold below the $K_M$ of *Sp*AS1$^{WT}$ toward 4-nitrophenyl sulfate **3a** to ensure that the observed improvements were representative for improvements of $k_{cat}/K_M$. The observed reaction rates (in $A_{400}$ s$^{-1}$) were normalized for the cell density ($OD_{600}$) for each individual well. The activity measurements of the wild-type typically showed a variation of ±30%. Therefore only variants with a >1.4-fold in crease in activity were selected for the next screening round.

#### 1.8 Screening step 4: approximation of catalytic efficiency measurements with whole cells

All optical measurements for this step (absorbance and fluorescence) were carried out using a TECAN infinite M200 microtiter plate reader. For each of the autodisplayed *Sp*AS1-variants selected after the microtiter plate-based activity screening for improved activity toward fluorescein disulfate **1a** and/or 4-nitrophenyl sulfate **3a**, we inoculated 3 mL LB-medium containing ampicillin (50-100 mg L$^{-1}$) with *E. cloni* 10G containing the respective pBAD-AT-$His_6$-*Sp*AS1 variant. The cultures were grown at 37 °C to an $OD_{600}$ ~ 0.5, at which point the cultures were cooled to 25 °C. Once the culture reached the desired temperature, expression of the autodisplayed *Sp*AS1 variant was induced by adding up to 0.02% (w/v) L-arabinose, followed by overnight growth at 25 °C. Cells were harvested by centrifugation and resuspended in 1 mL SID buffer pH 7.5. In each well we mixed 30 µL of this cell suspension with 90 µL SID buffer pH 7.5, 8 wells for each 'whole cell Michaelis-Menten curve', and measured the $OD_{600}$. We prepared the same mixture in each well of a black opaque microtiterplate.

To the cells in the black opaque microtiteplate we subsequently added 80 µL substrate solution (1-10 mM fluorescein disulfate **1a**, final substrate concentration ([S]) ranging 0.4-4.0 mM) and measured the increase in fluorescence ($\lambda_{ex}$ = 490 nm; $\lambda_{em}$ = 515 nm) over time ($V_{init}$ in $F_{counts}$ $s^{-1}$). The corrected rate ($V_{corr}$) of fluorescein formation for each well was calculated using equations 1-3, using the average of the 8 $OD_{600}$ measurements mentioned above for the corrections of $F_{response}$. Subsequently $V_{corr}$ was converted to the number of fluorescein molecules formed ($N_P$) per second using equation 4, in which $Volume$ = $2\times10^{-4}$ L (200 µL), and $N_{av}$ = $6.02\times 10^{23}$ molecules $mol^{-1}$.

$$N_{\mathrm{P}}\ \mathrm{s}^{-1} = V_{\mathrm{corr}} \times Volume \times N_{av} \quad (4)$$

For measuring whole cell kinetics toward 4-nitrophenyl **3a** we added 80 µL 2-20 mM 4-nitrophenyl sulfate **3a** (final concentration of 4-nitrophenyl sulfate **3a** ranging 0.8-8.0 mM) to the wells containing 120 µL cell-buffer mix as prepared above and measured the initial rates of the increase in $A_{400}$ ($V_{init}$ in $A_{400}$ $s^{-1}$). The number of product molecules per second was calculated from the rate of increase in absorbance at 400 nm ($V_{init}$) using equation 5, in which $\epsilon_{400}$ = 14441 $M^{-1}$ $cm^{-1}$, $d$ = 0.58 cm, $Volume$ = $2\times10^{-4}$ L (200 µL), and $N_{av}$ = $6.02\times 10^{23}$ molecules $mol^{-1}$.

$$N_{\mathrm{P}}\ \mathrm{s}^{-1} = \frac{V_{\mathrm{init}}}{\epsilon_{400} \times d} \times Volume \times N_{av} \quad (5)$$

The observed rate of product formation for both substrates ($V_{obs}$ in $N_P$ $cell^{-1}$ $s^{-1}$) was calculated by dividing the number of fluorescent/aborbant molecules formed ($N_P$) per second by the number of cells ($N_{cells}$) present in each well according to equation 6.

$$V_{\mathrm{obs}} = \frac{N_{\mathrm{P}}\ \mathrm{s}^{-1}}{N_{\mathrm{cells}}} \quad (6)$$

The number of cells was calculated from the $OD_{600}$ measured for each well, corrected for the background signal from the microtiter plate ($OD_{600}^{background}$ = 0.0357) using equation 7, in which $d$ = 0.35 cm, $Volume$ = 0.12 mL (120 µL) and $N_{\mathrm{cells\ mL^{-1}}\ at\ \mathrm{OD_{600}}=1}$ = $5\times10^{8}$. For the calculation of the number of cells present during the fluorescence measurement again the average of the 8 respective wells was used.

$$N_{\mathrm{cells}} = \frac{\mathrm{OD}_{600} - 0.0357}{d} \times Volume \times N_{\mathrm{cells\ mL^{-1}}\ at\ \mathrm{OD_{600}}=1} \quad (7)$$

Observed reaction rates ($V_{obs}$ in $N_P$ $cell^{-1}$ $s^{-1}$) were plotted against the concentration of substrate (fluorescein disulfate **1a** or 4-nitrophenyl sulfate **3a**, [S]) and kinetic parameters $V_{max}$ and $K_M$ were determined by fitting the data to equation 8.

$$V_{\mathrm{obs}} = \frac{V_{\mathrm{max}} \times [\mathrm{S}]}{K_{\mathrm{M}} + [\mathrm{S}]} \quad (8)$$

The errors ($\delta$) listed for $V_{max}$ and $K_M$ in Tables S2-S5 correspond to the errors from least-squares fitting of the Michaelis-Menten plots to equation 8. The respective errors for the $V_{max}/K_M$-values are calculated using equation 9.

$$\delta\frac{V_{\text{max}}}{K_{\text{M}}} = \frac{V_{\text{max}}}{K_{\text{M}}} \times \sqrt{(\frac{\delta V_{\text{max}}}{V_{\text{max}}})^2 + (\frac{\delta K_{\text{M}}}{K_{\text{M}}})^2} \quad (9)$$

#### 1.9 Protein production and purification

For production of each strep-tagged *Sp*AS1 variant, *E. coli* TOP10 cells containing the respective pASKIBA5$^{+}$-*Sp*AS1 plasmid constructs were grown in 100-400 mL of 2×YT medium containing ampicillin (50-100 mg $L^{-1}$) at 37 °C to an $OD_{600}$ ~ 0.5, at which point the culture was cooled to 28 °C. Once the culture had reached the desired temperature, overexpression of the respective strep-tagged *Sp*AS1 variants was induced by adding up to 200 µg $L^{-1}$ anhydrotetracycline followed by overnight growth at 28 °C.

Cells were harvested by centrifugation and resuspended in 50 mM Tris-HCl pH 8.0 containing an EDTA-free protease inhibitor cocktail (cOmplete™ EDTA-free, Sigma). Crude cell lysate was obtained by mixing the resuspended cells with an equal volume of 1× BugBuster® and adding Lysonase™ (Promega, 3 µL $g^{-1}$ wet cell weight), followed by gentle mixing at room temperature for 15-30 minutes. The crude lysate was centrifuged at 18,000×g for 60 minutes and the resulting supernatant was passed through a 0.45 µm sterile filter. The filtered cleared cell extract was subsequently loaded onto a 1 mL Strep-Tactin column equilibrated in 100 mM Tris-HCl pH 8.0 + 150 mM NaCl. Unbound protein was removed by washing extensively with 100 mM Tris-HCl pH 8.0 + 150 mM NaCl. The strep-tagged protein was eluted with 2.5 mM *d*-desthiobiotin in 100 mM Tris-HCl pH 8.0 + 150 mM NaCl. Protein containing fractions were either used immediately or pooled and concentrated to 100-200 µM, divided into appropriate aliquots, flash frozen in liquid nitrogen and stored at -20 °C.

#### 1.10 Measurement of kinetic parameters for purified *Sp*AS1 variants

All activity measurements with purified *Sp*AS1 variants were performed in SID buffer pH 7.5 in the presence of 0.6 mM $MnCl_2$ at 30 °C. The rate of enzymatic hydrolysis of fluorescein disulfate **1a** was followed by monitoring the release of fluorescein (**1b**) using fluorescence ($\lambda_{ex}$ = 490 nm; $\lambda_{em}$ = 515 nm) over time in black opaque microtiter plates ($V_{init}$ in $F_{counts}$ $s^{-1}$), using a TECAN infinite M200 microtiter plate reader. These initial rates were converted to the observed rate constant ($k_{obs}$) according to equation 10.

$$k_{\text{obs}} = (\frac{V_{\text{init}}}{F_{\text{response}}} - V_{\text{chem}})/[\text{Enz}] \quad (10)$$

In which the response factor for fluorescein fluorescence ($F_{response}$) under the given reaction conditions (SID buffer at pH 7.5) and detection settings ($\lambda_{ex}$ = 490 nm; $\lambda_{em}$ = 515 nm, gain = 30) was $4.37\times10^{8}$ fluorescent counts $M^{-1}$, and typical enzyme concentrations ([Enz]) ranged from 40-300 nM. The contribution of chemical hydrolysis ($V_{chem}$) for fluorescein disulfate ($k_{uncat}$ = $2.4\times10^{-8}$ $s^{-1}$) is relevant with respect to the enzymatic rate at the used enzyme concentrations. $V_{chem}$ was calculated for each substrate concentration ([S]) according to equation 2.

The rate of enzymatic hydrolysis of 4-nitrophenyl sulfate **3a** was followed by monitoring the release of 4-nitrophenolate **3b** using absorbance (λ= 400 nm) over time, using either a TECAN infinite M200 or Spectramax ABS Plus microtiter plate reader. The resulting initial rates ($V_{\text{init}}$ in $A_{400}$ $s^{-1}$) for formation of nitrophenolate **3b** were converted to the observed rate constants ($k_{\text{obs}}$ in $s^{-1}$) using equation 11.

$$k_{\text{obs}} = \frac{V_{\text{init}}}{\epsilon_{400\text{ nm}} \times d \times [\text{Enz}]} \tag{11}$$

In which the extinction coefficient of 4-nitrophenol in SID at pH 7.5 ($\epsilon_{400\text{ nm}}$) is 14441 $M^{-1}$ $cm^{-1}$ for the TECAN infinite M200 and 14828 $M^{-1}$ $cm^{-1}$ for the Spectramax ABS Plus, the path length in a microtiter plate well containing 200 µL liquid ($d$) is 0.58 cm and the concentration of the respective *Sp*AS1 variants ([Enz] was typically 3-5 nM. Chemical hydrolysis of 4-nitrophenyl sulfate ($k_{\text{uncat}}$ = $4.3\times10^{-10}$ $s^{-1}$ is too low to be relevant in comparison to the *Sp*AS1-catalyzed reaction ($k_{\text{cat}}/K_{\text{M}}$ *Sp*AS1$^{\text{WT}}$ = $4.8\times10^{3}$ $s^{-1}$ $M^{-1}$). Therefore, correction of $k_{\text{obs}}$ for chemical hydrolysis is not needed for determining the kientic parameters of *Sp*AS1-catalyzed 4-nitrophenyl sulfate conversion.

Observed rate constants ($k_{\text{obs}}$'s) were plotted against the concentration of substrate ([S]), and kinetic parameters $k_{\text{cat}}$, $K_{\text{M}}$ and $K_{\text{SI}}$ (substrate inhibition constant) were determined by fitting the data to equation 12 or 13.

$$k_{\text{obs}} = \frac{k_{\text{cat}} \times [\text{S}]}{K_{\text{M}} + [\text{S}]} \tag{12}$$

$$k_{\text{obs}} = \frac{k_{\text{cat}} \times [\text{S}]}{K_{\text{M}} + [\text{S}] + \frac{[\text{S}]^2}{K_{\text{SI}}}} \tag{13}$$

In order to obtain a reliable fit to equation 12 or 13, typically 16 (for fluorescein disulfate **1a**) or 32 (4-nitrophenyl sulfate **3a**) initial rate measurements were performed in such a way that the substrate concentrations used varied from at least 5-fold below the estimated $K_{\text{M}}$-value to at least 5-fold above the estimated $K_{\text{M}}$-value (with increments that increase with increasing substrate concentration), wherever substrate solubility allowed for it. The $K_{\text{M}}$-value was assumed to be similar to the value for *Sp*AS1$^{\text{WT}}$ (Table S6 and S7). If the latter substrate range resulted in difficult to fit data, lower substrate concentrations were used (see Figure S11 and S12 for examples of the Michaelis-Menten plots for several *Sp*AS1-variants).

The errors ($\delta$) listed for $k_{\text{cat}}$, $K_{\text{M}}$ and $K_{\text{SI}}$ in Table S6 and S7 correspond to the errors from least-squares fitting of the data points to equations 12 and 13. The error for $k_{\text{cat}}/K_{\text{M}}$ is calculated using equation 14.

$$\delta\frac{k_{\text{cat}}}{K_{\text{M}}} = \frac{k_{\text{cat}}}{K_{\text{M}}} \times \sqrt{(\frac{\delta k_{\text{cat}}}{k_{\text{cat}}})^2 + (\frac{\delta K_{\text{M}}}{K_{\text{M}}})^2} \tag{14}$$

#### 1.11 Calculation of the expected rate constant of uncatalyzed conversion of fluorescein disulfate 1a

We derived the rate constant for spontaneous hydrolysis of fluorescein disulfate **1b** ($k_{\text{uncat}}$) from a linear free energy relationship (log[$k_{\text{uncat}}$] *vs* p$K_a^{\text{leaving group}}$) for the uncatalyzed hydrolysis of arylsulfates (Figure S22B), assuming a p$K_a^{\text{leaving group}}$ for fluorescein disulfate **1b** of 6.4. The rates of conversion for each of the arylsulfates used to construct the LFER was calculated according to equation 15.

$$k_{\text{uncat}} = \frac{k_B \times T}{h} \times e^{-\Delta G^{\ddagger}/R \times T} \tag{15}$$

In which $k_B$ is the boltzmann constant (1.381×10$^{-23}$ J K$^{-1}$), $h$ is the planck constant (6.63×10$^{-34}$ J s$^{-1}$), $R$ is the gas constant (8.314 J K$^{-1}$ mol$^{-1}$) and $T$ is the temperature at which the reaction takes place (at 30 °C T = 303 K). The Gibbs energy of activation ($\Delta G^{\ddagger}$ in J mol$^{-1}$) was calculated from the published data[6] for the enthalpy and entropy of activation ($\Delta H^{\ddagger}$ in cal mol$^{-1}$ and $\Delta S^{\ddagger}$ in cal mol$^{-1}$ K$^{-1}$), according to equation 16.

$$\Delta G^{\ddagger} = 4.184 \times (\Delta H^{\ddagger} - T \times \Delta S^{\ddagger}) \tag{16}$$

#### 1.12 Conversion of transformation efficiencies to suitable units for comparison between plasmids of different sizes

The efficiency of transformation of plasmid DNA molecules is typically expressed as the number of colony forming units per µg plasmid DNA ($TE_{cfu/\mu g}$). In order to compare between plasmids of differing sizes, this number has to be converted into the fraction of DNA molecules that yields a colony after transformation ($TE_{fraction}$) by dividing the $TE_{cfu/\mu g}$ through the number of plasmid DNA molecules in a µg (= 10$^{-6}$ g) of plasmid DNA according to equation 17,

$$TE_{fraction} = \frac{TE_{cfu/\mu g}}{(10^{-6}/M_W^{plasmid}) \times N_{av}} \tag{17}$$

in which $N_{av}$ = 6.02×10$^{23}$ molecules mol$^{-1}$. Equation 15 was used to convert the reported efficiency for the transformation of plasmid pUC19[7] (GenBank accession number M77789) to *E. cloni* 10G ELITE electrocompetent cells ($TE_{cfu/\mu g}$ = 2×10$^{10}$ cfu µg$^{-1}$) to its $TE_{fraction}$. The molecular weight of pUC19 ($M_W^{plasmid}$=1660926.62 g mol$^{-1}$) was calculated using the online calculator from molbiotools (http://www.molbiotools.com/dnacalculator.html). The $TE_{fraction}$ for pUC19 = 0.055, i.e. 1 in 18 plasmid molecules results in the formation of a colony.

For the pBAD-AT-His$_6$-*Sp*AS1 construct we determined the $TE_{fraction}$ using equation 18,

$$TE_{fraction} = \frac{N_{cf}}{[plasmid] \times V_{transformed} \times N_{av}} \tag{18}$$

in which $N_{cf}$ is the number of colonies formed and $V_{transformed}$ is the total volume (in L) of DNA that was transformed to 25 µL *E. cloni* 10G ELITE electrocompetent cells. The concentration of plasmid was calculated from the A$_{260}$ measurement using an extinction coefficient at λ= 260 nm calculated form the pBAD-AT-His$_6$*Sp*AS1 sequence using the online calculator from molbiotools ($\varepsilon_{260}$ = 116508208 $M^{-1}$ $cm^{-1}$). The transformation of 1 µL pBAD-AT-His$_6$-*Sp*AS1 ($[plasmid]$ = 2.08×$10^{-12}$ M), resulted in 342 colonies. As a result the $TE_{fraction}$ = 2.73×$10^{-4}$, i.e. 1 in 3661 plasmid molecules results in the formation of a colony. When the transformed cells were plated directly onto a nitrocellulose filter sitting on top of the growth medium, only 105 colonies formed. For the latter the $TE_{fraction}$ = 8.39×$10^{-5}$, i.e. 1 in 11925 plasmid molecules results in the formation of a colony.

### 2 Expanded discussion

#### 2.1 Comparison cell lysis *vs.* autodisplay

##### 2.1.1 Recovery of genetic diversity after microdroplet sorting

In order to compare the recovery of genetic diversity for our lysis-free method with current lysis-based methods, it has to be established what the maximum possible recovery for the lysis-based methods is.

When plasmid DNA is recovered from single cell lysates in microdroplets, the amount of DNA molecules that can maximally be recovered from a single droplet depends on the copy number of that particular plasmid. Additionally the DNA purification procedure can be expected to result in a ∼20% loss.[1] In order to recover the diversity for further testing, it has to be re-transformed to *E. coli*. For high-copy-number plasmids (300-400 copies/cell) such as pUC19[7] and pRSFDuet,[1] this means a single sorted droplet (=1 cell) will yield ∼300 plasmid molecules. For transformation of pUC19 (2.7 kb) to commercially available competent cells such as electrocompetent *E. cloni* 10G ELITE, 1 in 18 plasmid DNA molecules will result in a transformant (see expanded methods for detailed calculation), which means 300/18×100% = 16700% is the maximum recovery that can be achieved. However, for larger plasmids, transformation is considerably less efficient, e.g. for pBAD-AT-His$_6$-*Sp*AS1 (7.1 kb) only 1 in 3661 plasmid DNA molecules results in a transformant (see expanded methods for detailed calculation). Additionally pBAD-AT, as well as many other overexpression plasmids, has a copy number of ∼20 copies/cell. In combination this means 20/3661×100% = 0.55% of genotypic diversity can maximally be recovered using re-transformation (assuming an unlikely 100% efficiency for the DNA isolation procedure). As a result the diversity would need to be oversampled at least 100/0.55 = 181-fold to cover the total diversity just once. As a result only a limited number of overexpression systems, i.e. based on efficiently transfecting, high copy-number plasmids, are currently suitable for reliable recovery of genetic diversity from single cell-lysates in micro-droplets.

During the lysis-free autodisplay-based procedure, the cells displaying the selected *Sp*AS1 variants stay intact during the screening procedure. As a consequence, direct plating of the sorted cells onto LB agar could result in full recovery of the sorted diversity. However, when we disrupt the droplets containing the sorted cells and than plated the resulting solution directly onto LB-agar plates containing ampicillin followed by overnight incubation at 37 °C, no colonies were observed after overnight growth. This means that the cells are either not viable or are viable but not culturable (VBNC). In case of the latter the cells are present and metabolically active, but lack the ability to grow.[8,9] The VBNC state is typical for bacterial cells that are stressed, which can be expected to be the case in the presence of toxic compounds or starvation. Based on the conditions our cells are exposed to: induction of overexpression at high cell densities, overexpression of a 'useless' protein on their cell surface and sitting alone in a water-in-oil droplet, sorted cells are very likely to be stressed. In particular protein overexpression, in which almost all cellular resources are depleted (as with starvation), can be expected to induce the VBNC state. We decided to alter the conditions to favor cell recovery. The sorting procedure was done in buffers containing glycerol (0.5% (v/v) final concentration in droplets), which could act both as a general stabilizer of cellular integrity and as a carbon source for maintenance of cell viability. Furthermore, we directly collected sorted droplets into a 1:9:20 mix of a mild surfactant (PFO) : fluorous oil (HFE7500) : Lucigen recovery medium. The latter is normally used to recover *E. coli* cells after electroporation, also a highly stressful event. Directly following the end of droplet collection, we added additional recovery medium supplemented with sodium pyruvate (0.5% (w/v) final concentration), which was previously reported to resuscitate *E. coli* cells that are in the VBNC state,[10] and allowed the cells to recover similar to the post-transfection recovery typically done for electroporated *E. coli* cells. The medium phase (recovery medium + cells) was collected and plated directly onto LB-agar containing ampicillin covered with a nitrocellulose filter and incubated overnight at 37 °C. This resulted in approximately 12% of the sorted cells to grow into individual colonies. Given that the maximum recovery that can be achieved with cell lysis followed by isolation of plasmid DNA and re-transformation of the plasmid DNA is 0.55% (see above), our direct live-cell regrowth method is >20-fold more efficient at recovering genetic diversity than the alternative. Further optimization of the recovery conditions can probably increase the percentage of recovered cells.

##### 2.1.2 Recovery and number of experimental steps during agar plate-based screening

Methods that rely on cell lysis during plate-based screening methods, analogous to the microdroplet-based procedures, require recovery of the genotype. For agar plate-based screening procedures that require cell lysis, genotype recovery is achieved by keeping a so-called reference plate. In short, the colonies containing the library variant growing on standard growth medium are replicated onto a nitrocellulose membrane. The colonies on the original plate are subsequently regrown and serve as a 'genotype' reference from which the plasmid encoding the positive variants can be recovered. The replicated colonies are incubated in the presence of a compound that induces expression of the library variant. Actual cell lysis for the activity measurements is done by repeated freeze thaw cycles with the colonies on the nitrocellulose membrane. For the autodisplay system the cells can be grown directly on a nitrocellulose membrane. Exposure to substrate can be simply done by transferring the filter with the live growing cells to medium containing substrate. Positive colonies can be directly used to inoculate the reference microtiter plate for the next step, thereby avoiding any mix-up during recovery of the correct genotype.

##### 2.1.3 Number of experimental steps during microtiterplate screening

When cell lysis is needed for activity measurements in microtiterplate-based screening methods, cells are typically lysed with a chemical agent, such as BugBuster®, often in combination with enzymatic cell lysis using lysozyme. In many cases these procedures also involve harvesting of the cells and/or removal of cell debris after lysis, which both require centrifugation.

In order to normalize the activity data, either a cell density ($OD_{600}$) or protein content (Bradford) measurement is required. When using cell density as reference, this has to be determined prior to cell lysis, which means that the variability in completeness of the lytic process can cause under and overestimation of activity levels. Direct measurement of protein is done post-lysis, but can only be done with cleared lysates and has to be done in a separate microtiter plate, which both require additional liquid handling.

In short: the cell lysis methods require at least 3 steps: *i*) separate measurement of cell density/protein content, *ii*) addition of lytic agent and *iii*) mixing of cell lysate and substrate. *E. coli* autodisplay requires in essence only one liquid handling step: mixing of cells and substrate.

#### 3 Supporting Figures

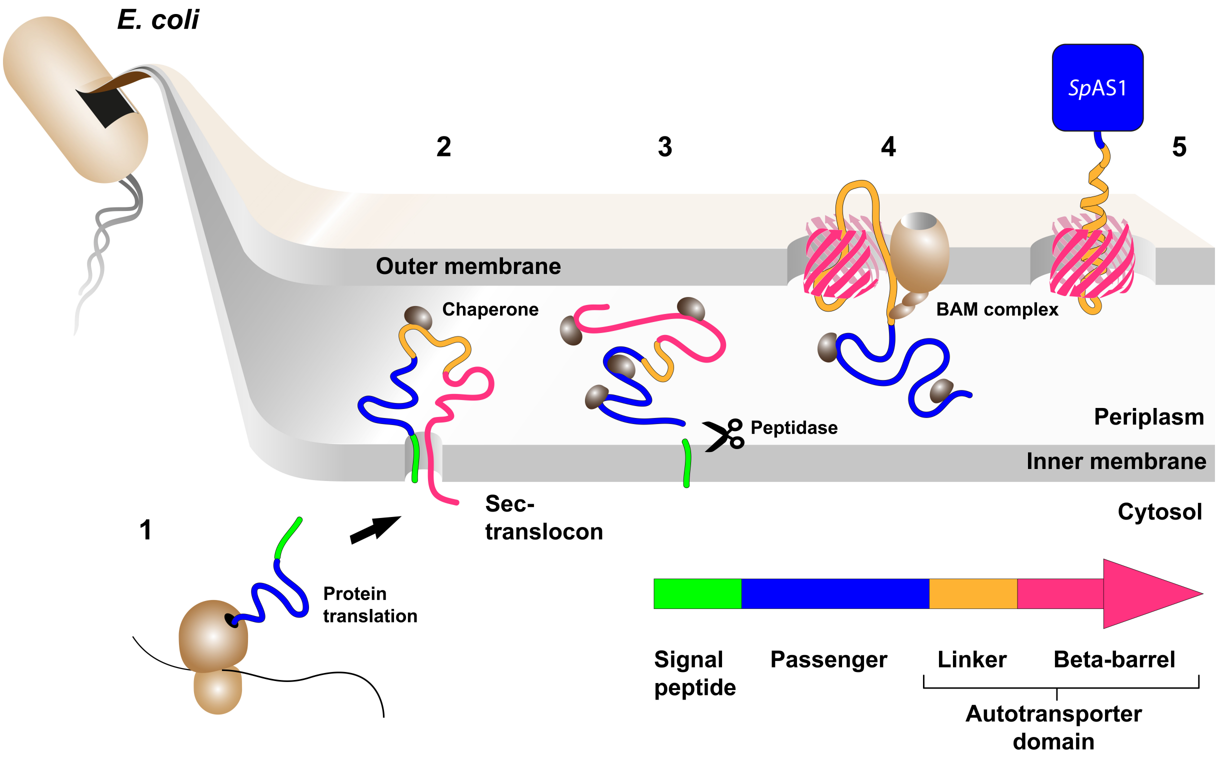


**Figure S1:** Schematic representation of autodisplay, based on the autotransporter type Va secretion system as used in this study. By the aid of an N-terminal signal peptide the precursor is translocated across the cytoplasmic membrane into the periplasm, where several chaperones are involved in maintaining an unfolded status. The C terminal β-sheet of the precursor interacts with the *E. coli* β-barrel assembly machinery (BAM), which inserts the β-barrel into the outer membrane and keeps it open until the passenger (natural or recombinant) is translocated to the cell surface. The Figure was adapted from Quehl *et al.*[11]

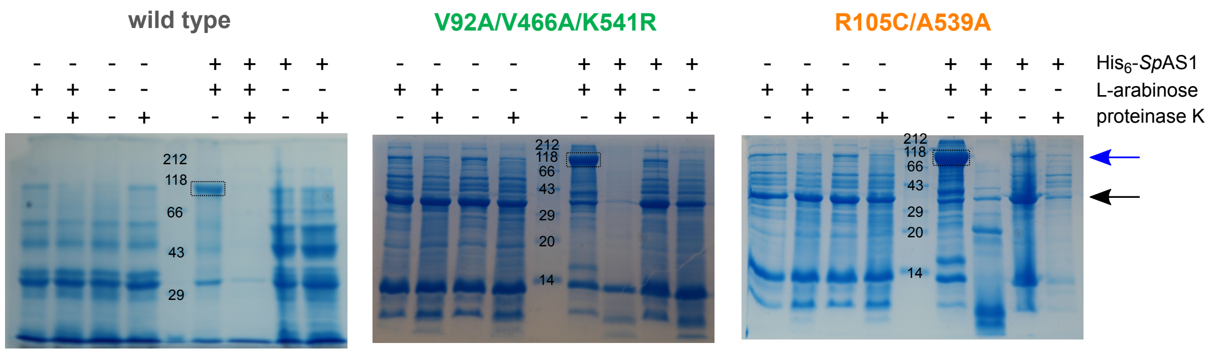


**Figure S2:** SDS-PAGE analysis of *Sp*AS1 autodisplay expression in the cell membrane of *E. cloni* 10G. The protein indicated with the blue arrow corresponds to the expected molecular mass of the *Sp*AS1-autotransporter fusion protein (113 kDa). The black arrow indicates natural outer membrane proteins (Omp) A, C, and F as internal standard. Proteinase K treatment of whole cells prior results in the disappearance of the *Sp*AS1-autotransporter fusion protein, indicating that *Sp*AS1 is displayed on the outside of *E. coli*.

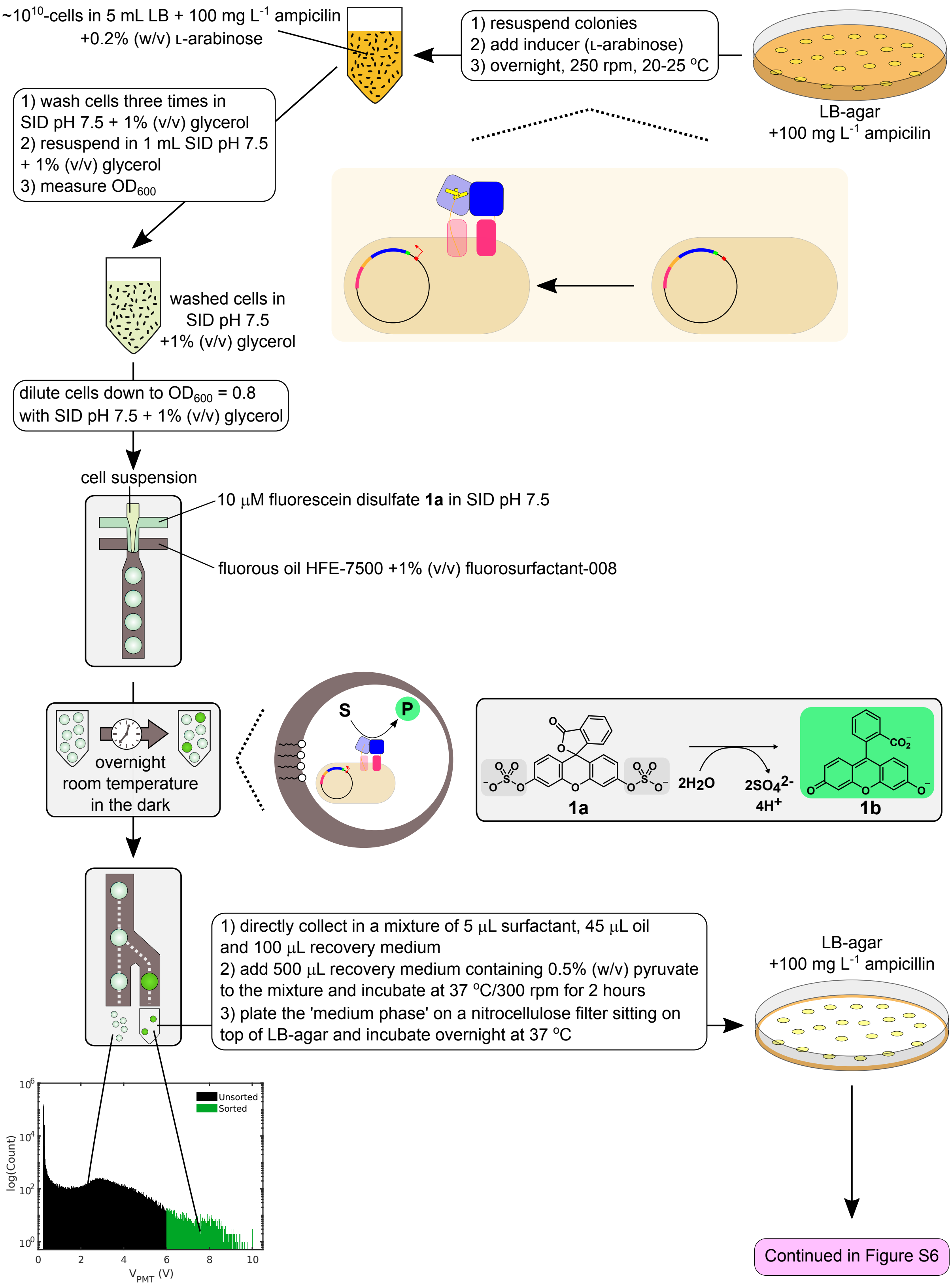


**Figure S3:** Procedure for testing *Sp*AS1-variants for activity toward fluorescein disulfate (sulfate monoester **1a**) (step 1 of the screening procedure). Experimental details regarding bulk-expression of the autodisplayed *Sp*AS1 variants, encapsulation of single cells into water-in-oil microdroplet, subsequent sorting of the microdroplets with >2-fold higher fluorescence signal after overnight incubation and subsequent recovery of the positive variants are indicated.

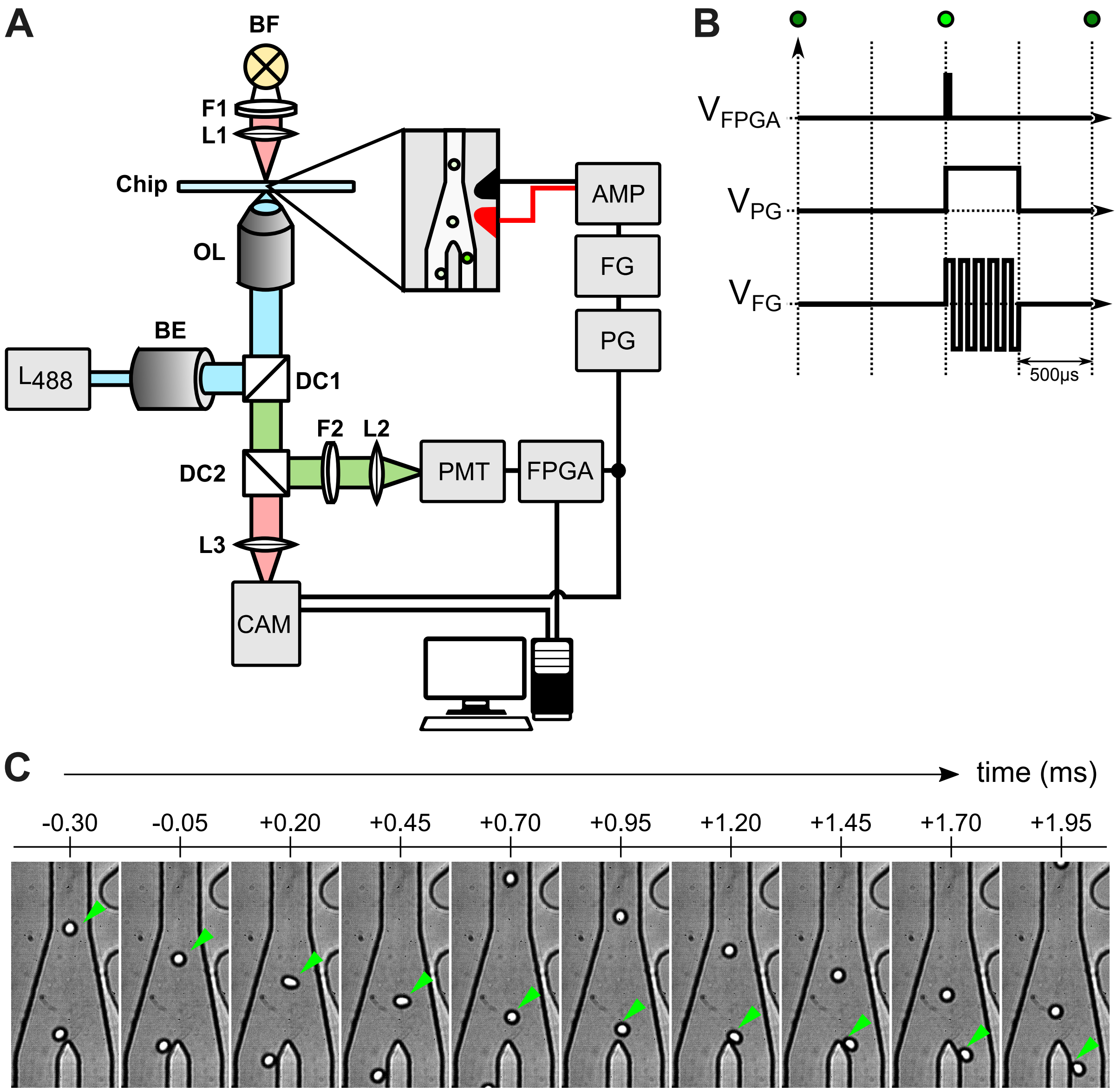


**Figure S4:** Fluorescence activated droplet sorter (FADS). **A**) Schematic representation of the set-up of the droplet sorter. PMT: photo multiplier tube; FPGA: field programmable gate array; PG: pulse generator, FG: function generator, CAM: high-speed camera; AMP: amplifier; BE: beam expander (10×); OL: objective lense; L488: 488nm laser; F1: 593nm longpass filter; DC1: 495nm dichroic mirror; DC2 555nm dichroic mirror; F2: 525/28 bandpass filter. **B**) Voltage pulse series used to sort droplets. The FPGA triggers the PG. The PG opens a gate for the FG which in turn applies a 10kHz square wave to the amplifier. The resulting AC field causes dielectrophoretic sorting of the selected droplet. **C**) A positive sorting event depicted in real time.

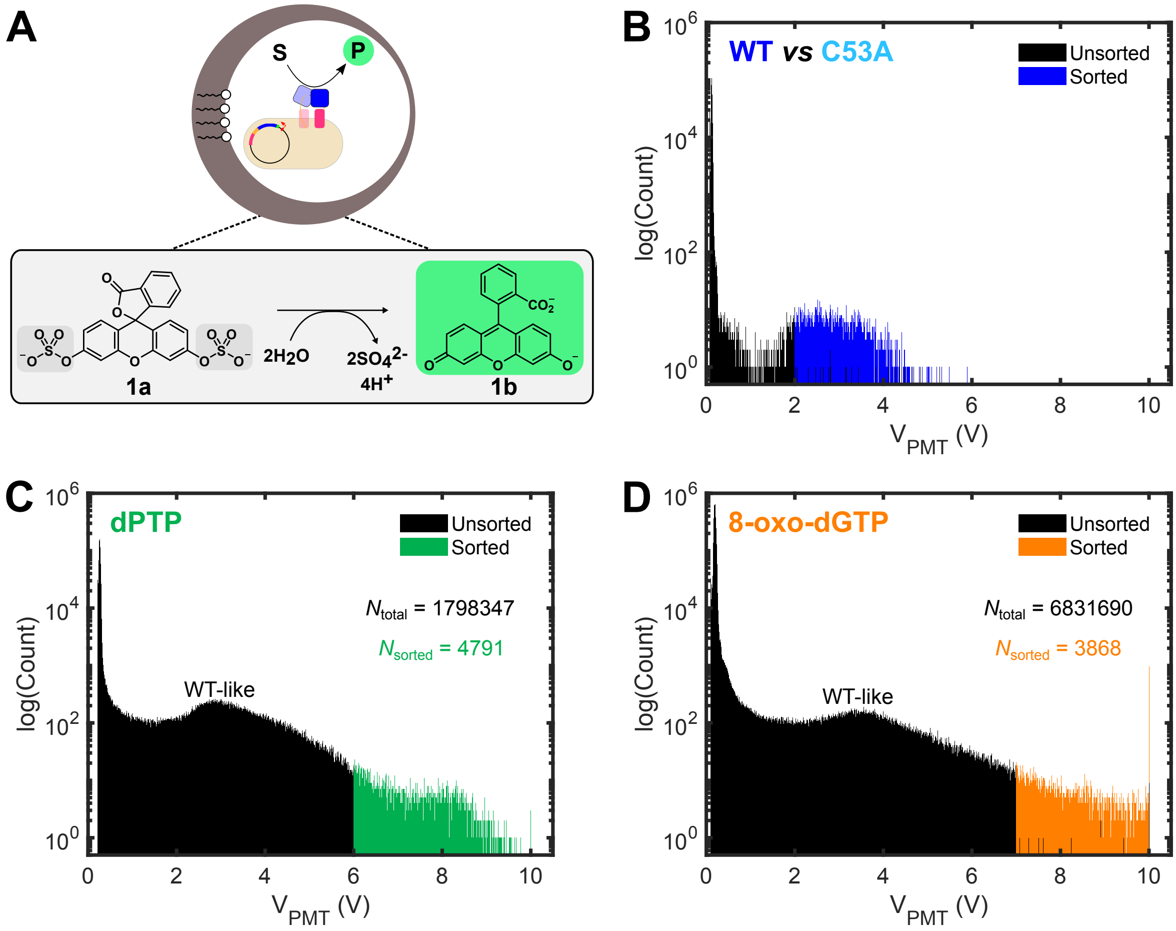


**Figure S5:** Histograms of droplets containing reaction buffer with 5 µM fluorescein disulfate **1a** and *E. cloni* cells displaying *Sp*AS1 variants. According to the Poisson distribution with $\lambda = 0.35$, ∼70% of the droplets is expected to contain no cells, ∼25% of the droplets contain a single cell and ∼5% contain 2 cells.The total number indicated ($N_{total}$) is the total number of droplets tested.

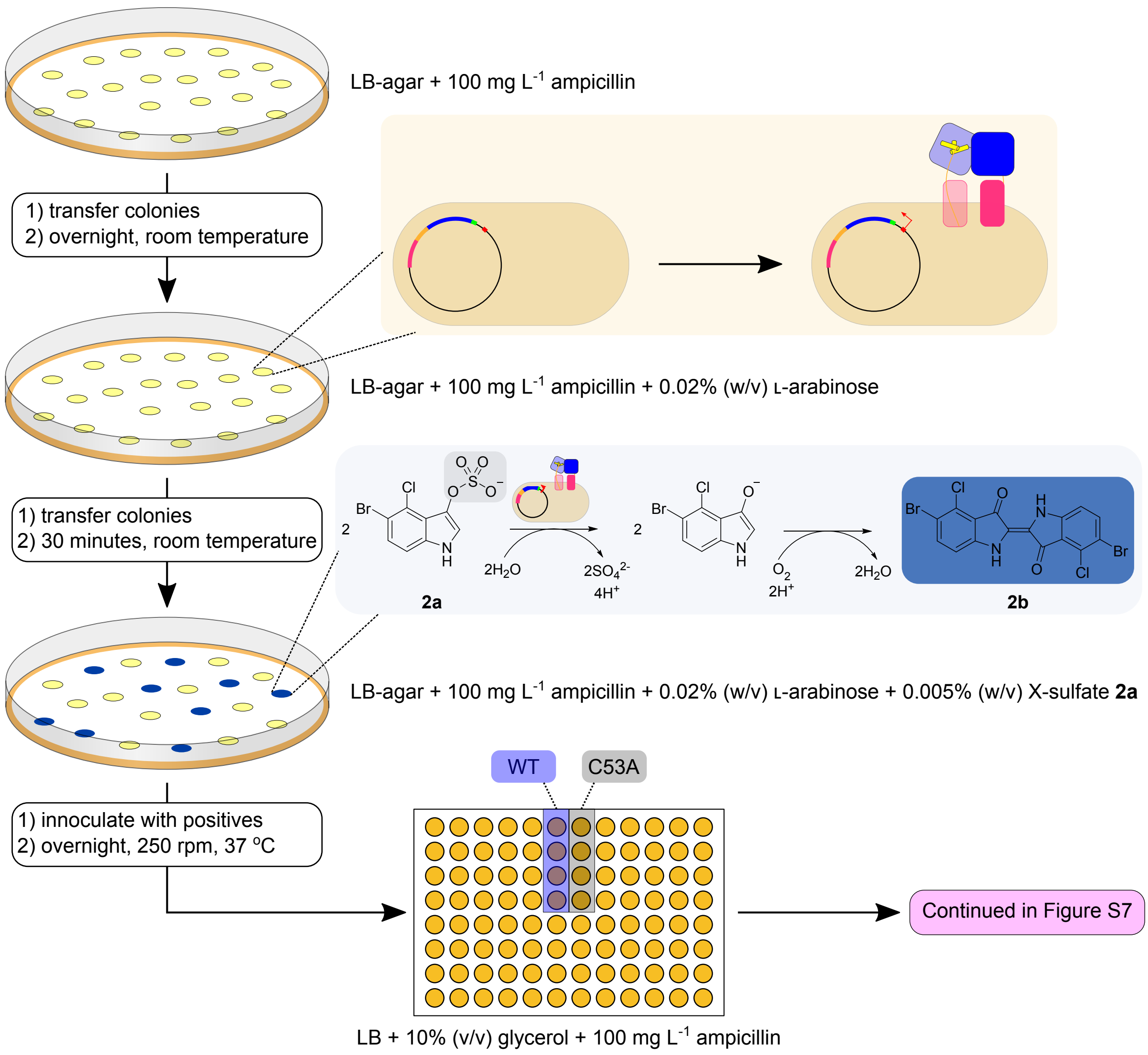


**Figure S6:** Procedure for testing *Sp*AS1-variants for activity toward sulfate monoester **2a** (step 2 of the screening procedure). Experimental details regarding expression of the autodisplayed *Sp*AS1 variants and criteria for scoring activity toward sulfate monoester **2a** are indicated.

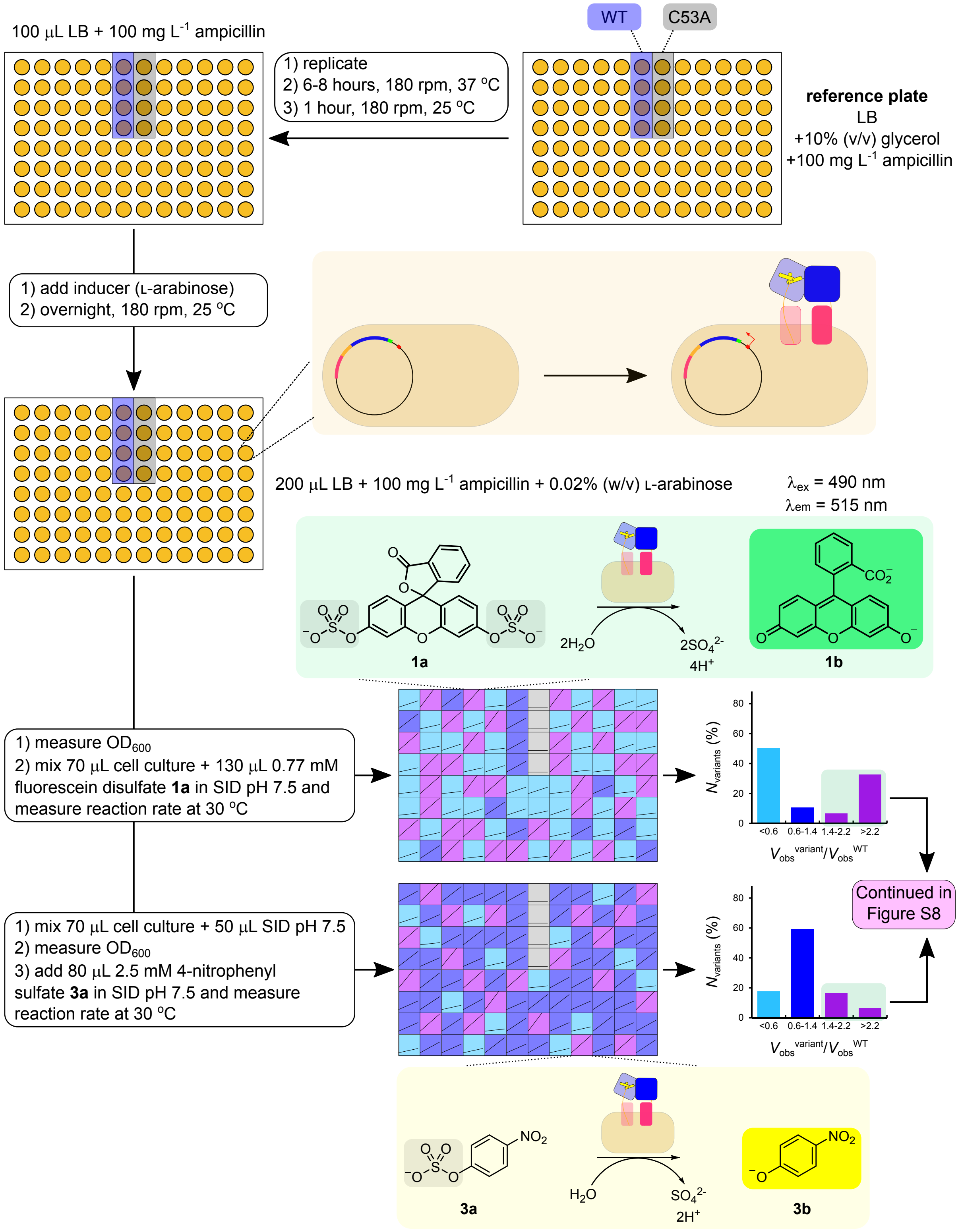


**Figure S7:** Procedure for side-by-side testing of *Sp*AS1-variants for activity toward fluorescein disulfate **1a** and 4-nitrophenyl sulfate **3a** (step 3 of the screening procedure). Experimental details regarding expression of the autodisplayed *Sp*AS1 variants and measurement of the initial rates toward fluorescein disulfate **1a** ([**1a**]$_{final}$ = 0.5 mM) and 4-nitrophenyl sulfate **3a** ([**3a**]$_{final}$ = 1.0 mM) are indicated.

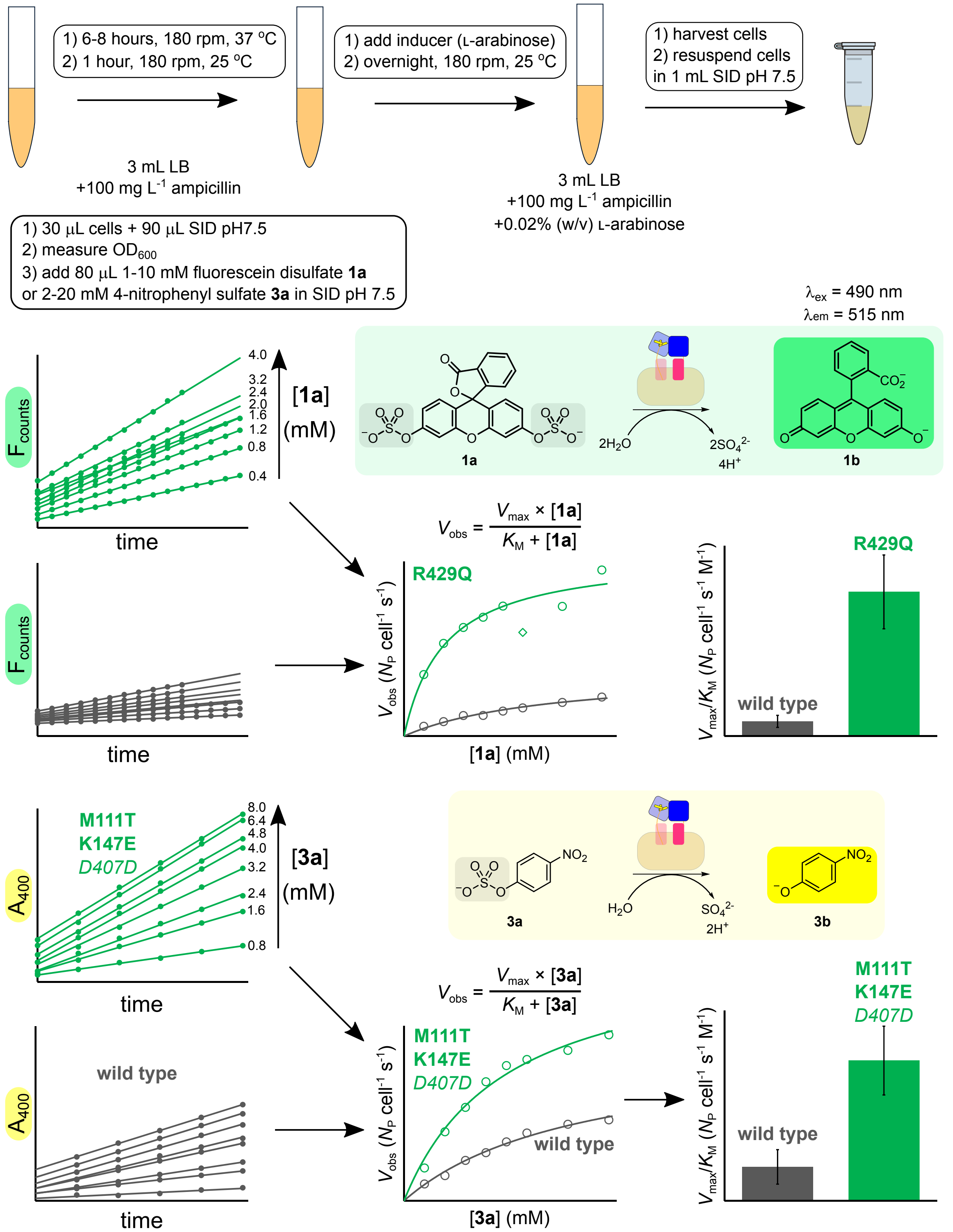


**Figure S8:** Procedure for side-by-side determination of a proxy for Michaelis-Menten kinetics toward fluorescein disulfate **1a** and 4-nitrophenyl sulfate **3a** for autodisplayed *Sp*AS1-variants (step 4 of the screening procedure). Experimental details regarding expression of the autodisplayed *Sp*AS1 variants and measurement of the initial rates for 8 different fluorescein disulfate **1a** and 4-nitrophenyl sulfate **3a** concentrations are indicated.

without droplet sorting | with droplet sorting

**A**

| $V_{obs}^{variant}/V_{obs}^{WT}$ **1a** \ $V_{obs}^{variant}/V_{obs}^{WT}$ **3a** | <0.6 | 0.6-1.4 | 1.4-2.2 | >2.2 |
|---|---|---|---|---|
| <0.6 | 70 | 31 | 9 | 2 |
| 0.6-1.4 | 8 | 28 | 9 | 1 |
| 1.4-2.2 | 3 | 8 | 1 | 1 |
| >2.2 | 1 | 4 | 0 | 0 |

**B**

| $V_{obs}^{variant}/V_{obs}^{WT}$ **1a** \ $V_{obs}^{variant}/V_{obs}^{WT}$ **3a** | <0.6 | 0.6-1.4 | 1.4-2.2 | >2.2 |
|---|---|---|---|---|
| <0.6 | 23 | 65 | 9 | 3 |
| 0.6-1.4 | 5 | 12 | 4 | 0 |
| 1.4-2.2 | 2 | 10 | 1 | 0 |
| >2.2 | 5 | 31 | 19 | 10 |

$\Sigma_{\text{to step 4}} = 94$

**C**

| $V_{obs}^{variant}/V_{obs}^{WT}$ **1a** \ $V_{obs}^{variant}/V_{obs}^{WT}$ **3a** | <0.6 | 0.6-1.4 | 1.4-2.2 | >2.2 |
|---|---|---|---|---|
| <0.6 | 162 | 23 | 0 | 0 |
| 0.6-1.4 | 7 | 8 | 1 | 1 |
| 1.4-2.2 | 2 | 1 | 0 | 0 |
| >2.2 | 5 | 3 | 0 | 0 |

**D**

| $V_{obs}^{variant}/V_{obs}^{WT}$ **1a** \ $V_{obs}^{variant}/V_{obs}^{WT}$ **3a** | <0.6 | 0.6-1.4 | 1.4-2.2 | >2.2 |
|---|---|---|---|---|
| <0.6 | 142 | 23 | 5 | 6 |
| 0.6-1.4 | 2 | 12 | 4 | 6 |
| 1.4-2.2 | 1 | 2 | 8 | 10 |
| >2.2 | 1 | 2 | 8 | 19 |

$\Sigma_{\text{to step 4}} = 72$

**Figure S9:** Sorting of autodisplayed *Sp*AS1 variants according to the combination of relative improvements (compared to autodisplayed *Sp*AS1$^{WT}$) for fluorescein disulfate **1a** and 4-nitrophenyl sulfate **3a** measured during screening step 3. All purple variants in panel **B** and **D** were selected for further testing in screening step 4 ($\Sigma_{\text{to step 4}}$).

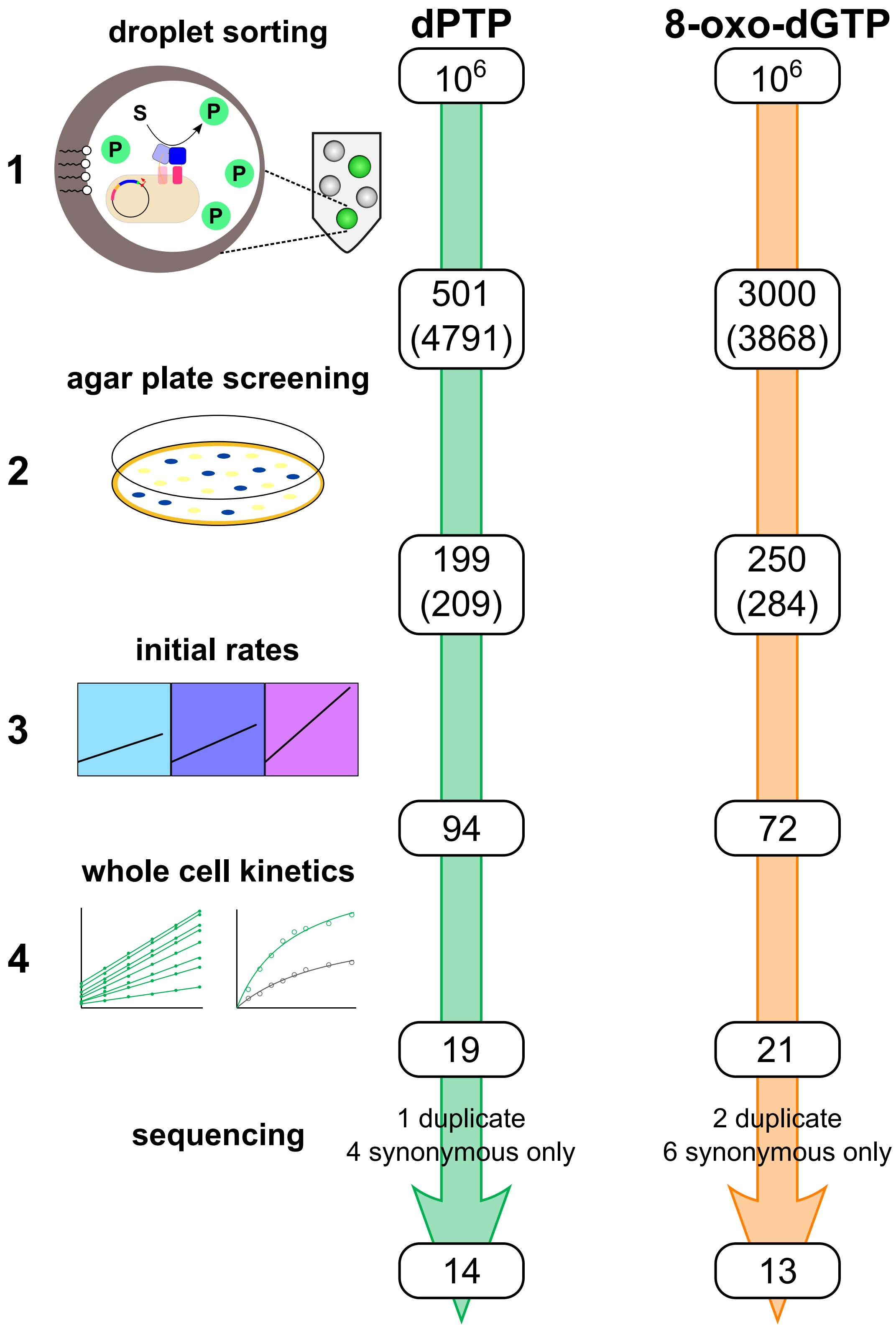


**Figure S10:** Numbers of variants selected after each step in the screening procedure. Sequence analysis after the final screening step ('whole cell kinetics') revealed duplicate hits (i.e. the same mutant found twice) and several variants that only contain synonymous mutations. The final numbers indicated are the variants for which the coding sequences were cloned into a vector for cytosolic expression in *E. coli* for which the catalytic efficiency ($k_{cat}/K_M$) of the corresponding purified proteins was determined for fluorescein disulfate **1a** and 4-nitrophenyl sulfate **3a**.

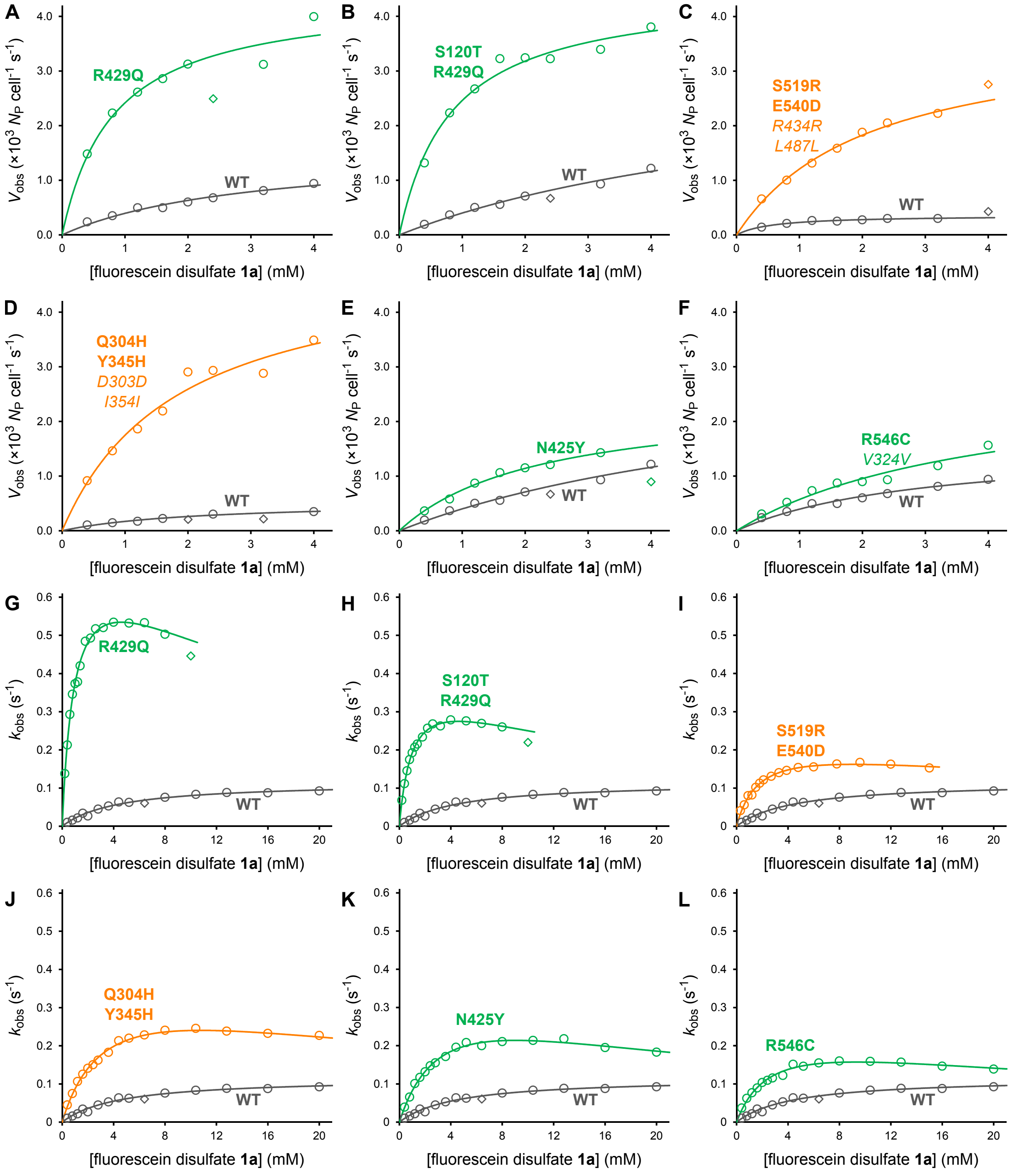


**Figure S11:** Michaelis-Menten curves for *Sp*AS1-catalyzed hydrolysis of fluorescein disulfate **1a** for the six variants with the highest $k_{cat}/K_M$ toward fluorescein disulfate **1a** for the autodisplayed (**A**-**F**) and purified (**G**-**L**) forms of the *Sp*AS1 variants. Data for the individual catalytic parameters toward fluorescein disulfate **1a** for all selected *Sp*AS1 variants are listed in Tables S2 and S3 (autodisplayed version) and Table S6 (purified protein).

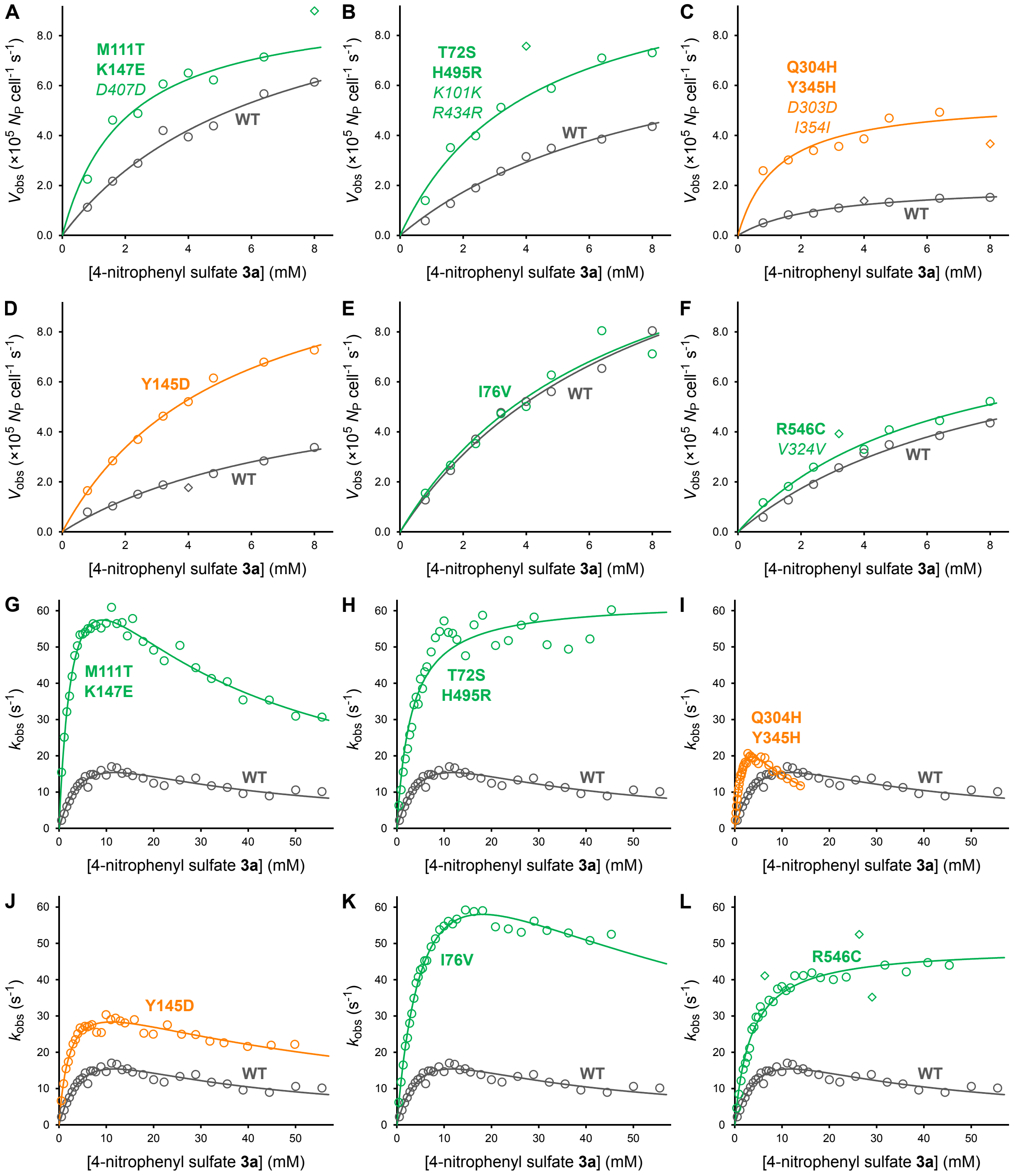


**Figure S12:** Michaelis-Menten curves for *Sp*AS1-catalyzed hydrolysis of 4-nitrophenyl sulfate **3a** for the six variants with the highest $k_{cat}/K_M$ toward 4-nitrophenyl sulfate **3a** for the autodisplayed (**A**-**F**) and purified (**G**-**L**) forms of the *Sp*AS1 variants. Data for the individual catalytic parameters toward 4-nitrophenyl sulfate **3a** for all selected *Sp*AS1-variants are listed in Tables S4 and S5 (autodisplayed version) and Table S7 (purified protein).

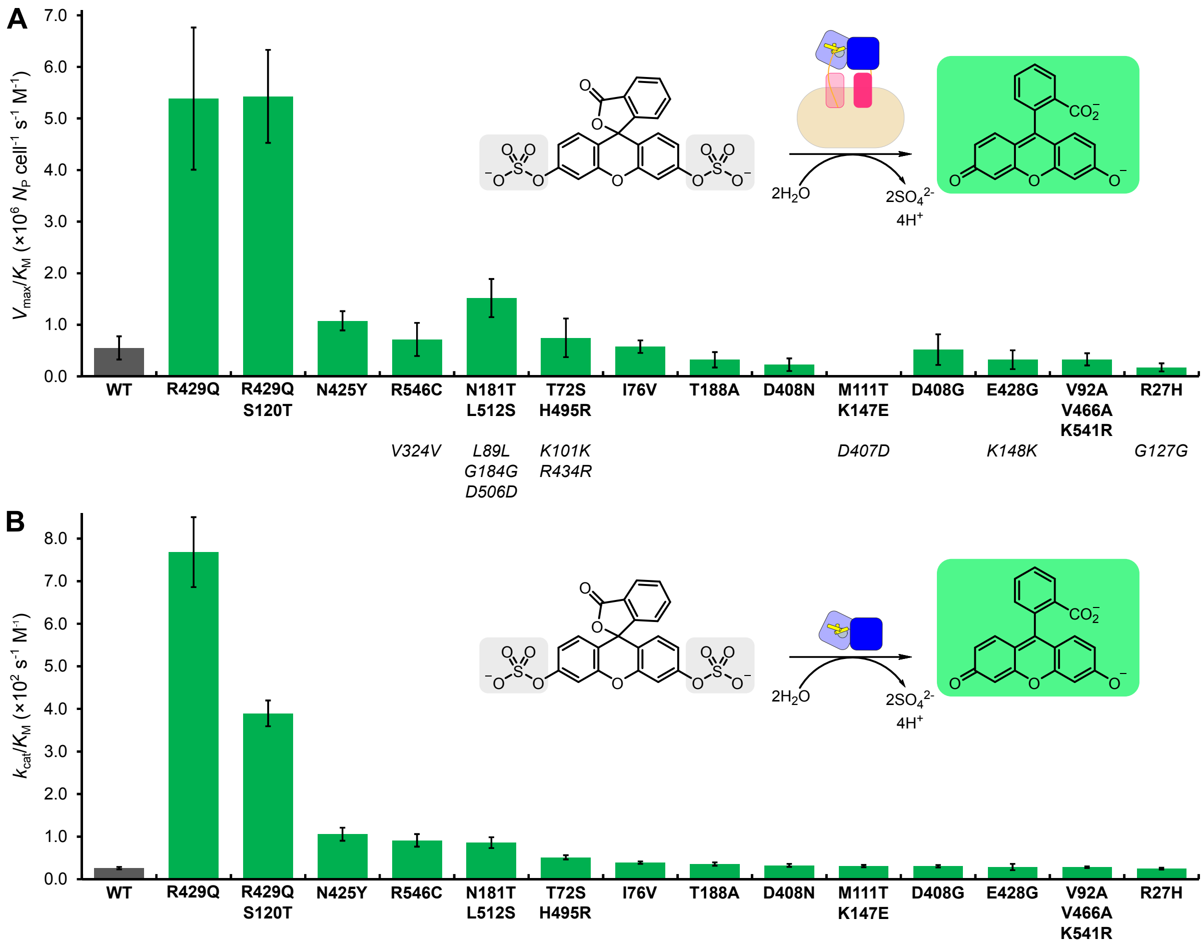


**Figure S13:** Catalytic performance toward fluorescein disulfate **1a** for all *Sp*AS1 variants selected from the **dPTP**-generated library. Data for the kinetic parameters for the autodisplayed (panel **A**) and purified (**B**) *Sp*AS1 variants are listed in Table S2 and S6 respectively.

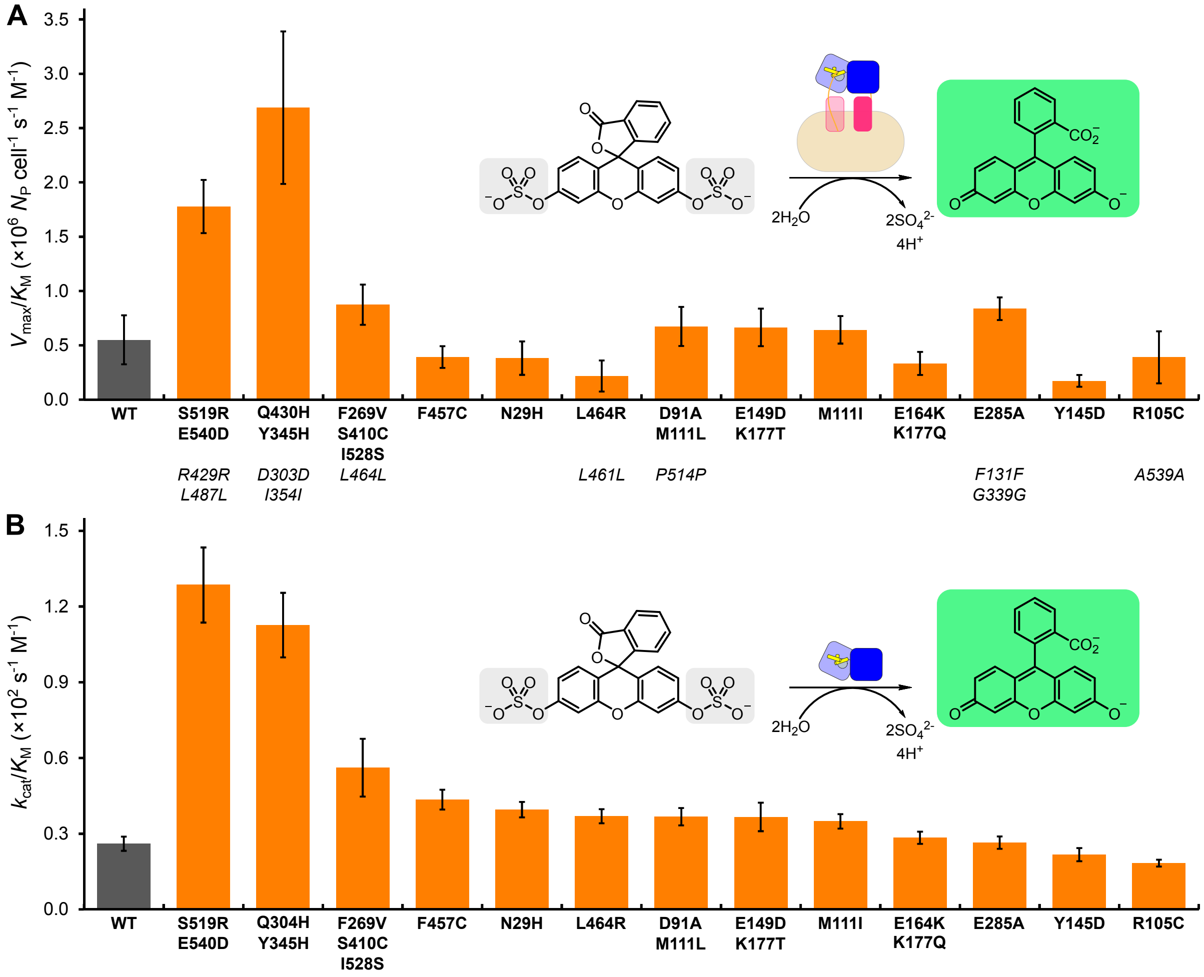


**Figure S14:** Catalytic performance toward fluorescein disulfate **1a** for all *Sp*AS1 variants selected from the **8-oxo-dGTP**-generated library. Data for the kinetic parameters for the autodisplayed (panel **A**) and purified (**B**) *Sp*AS1 variants are listed in Table S3 and S6 respectively.

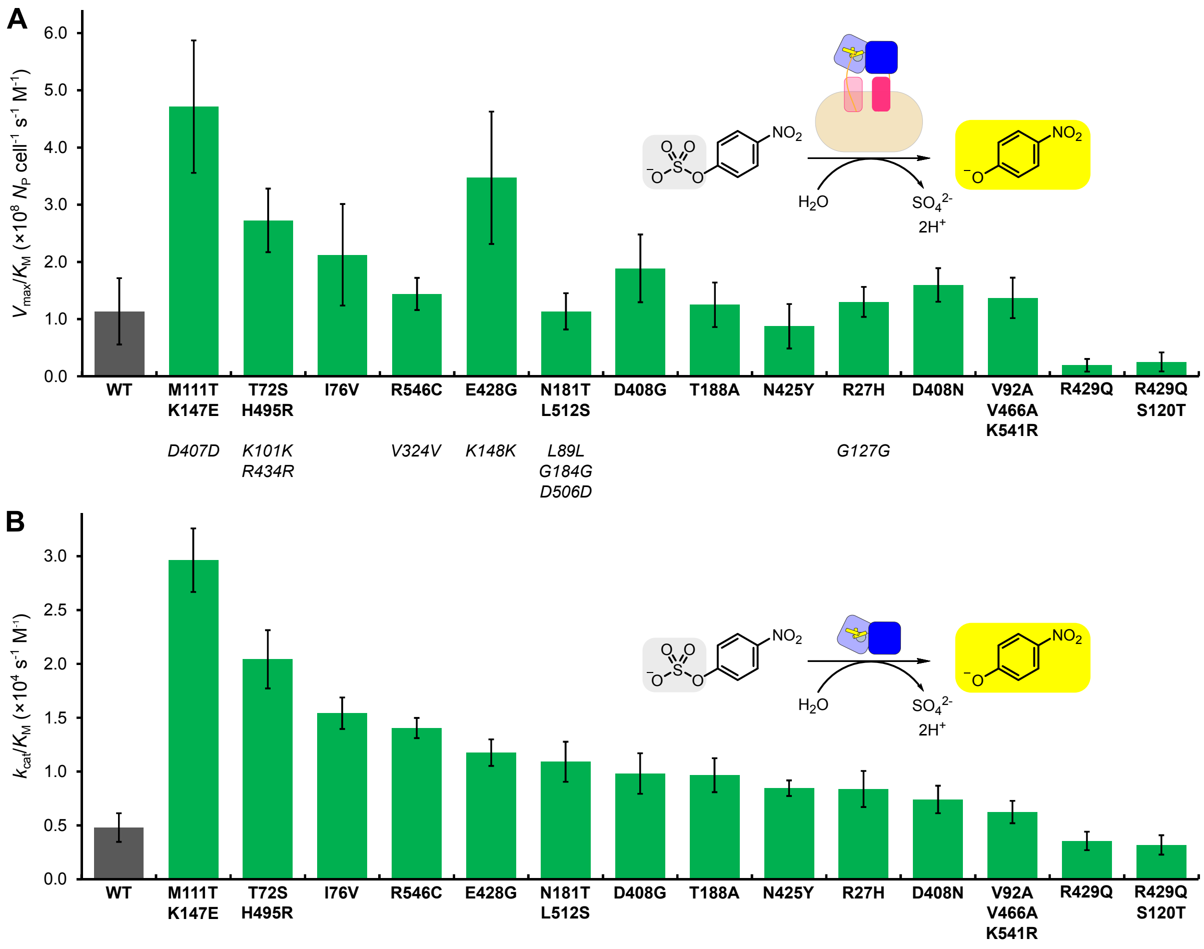


**Figure S15:** Catalytic performance toward 4-nitrophenyl sulfate **3a** for all *Sp*AS1 variants selected from the **dPTP**-generated library. Data for the kinetic parameters for the autodisplayed (panel **A**) and purified (**B**) *Sp*AS1 variants are listed in Table S4 and S7 respectively.

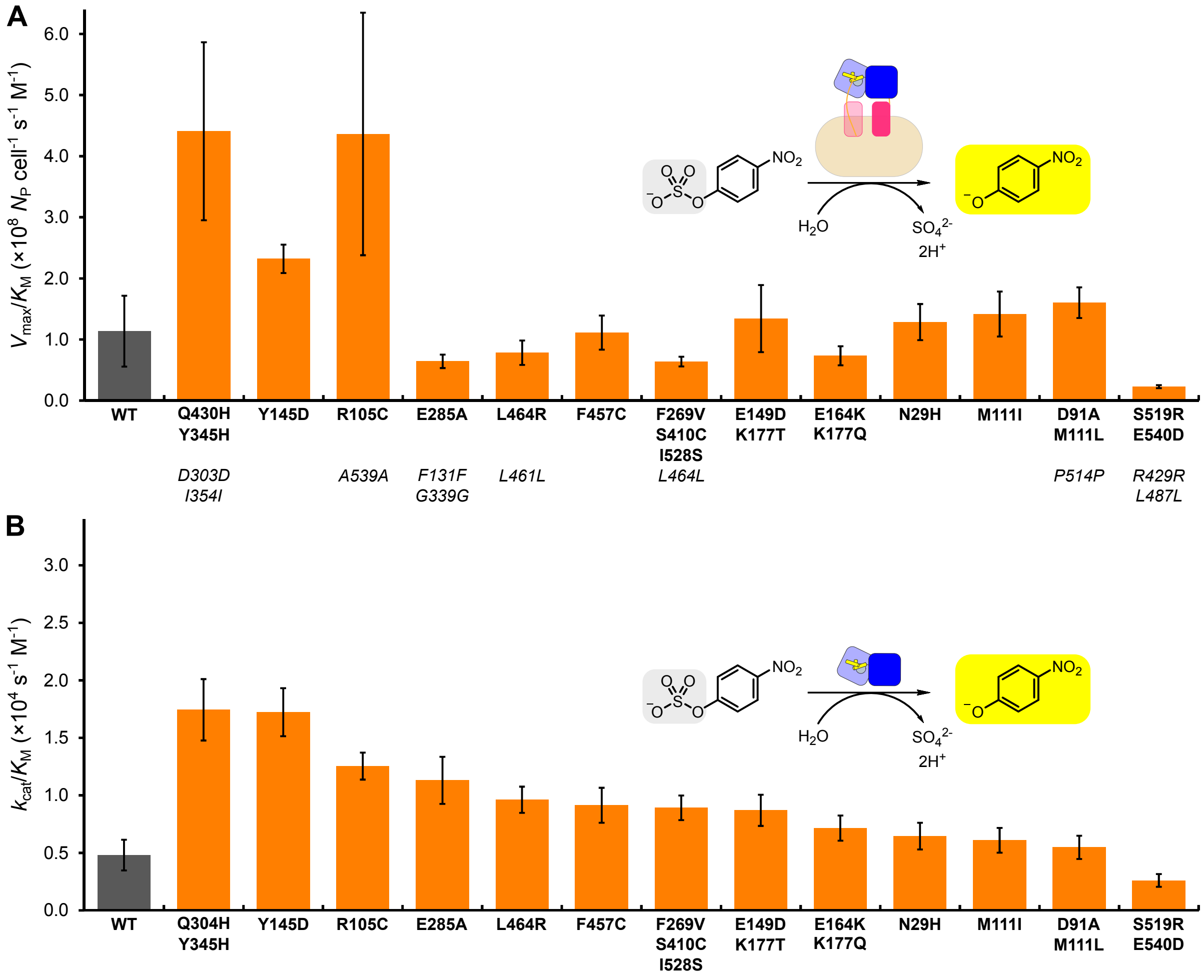


**Figure S16:** Catalytic performance toward 4-nitrophenyl sulfate **3a** for all *Sp*AS1 variants selected from the **8-oxo-dGTP**-generated library. Data for the kinetic parameters for the autodisplayed (panel **A**) and purified (**B**) *Sp*AS1 variants are listed in Table S5 and S7 respectively.

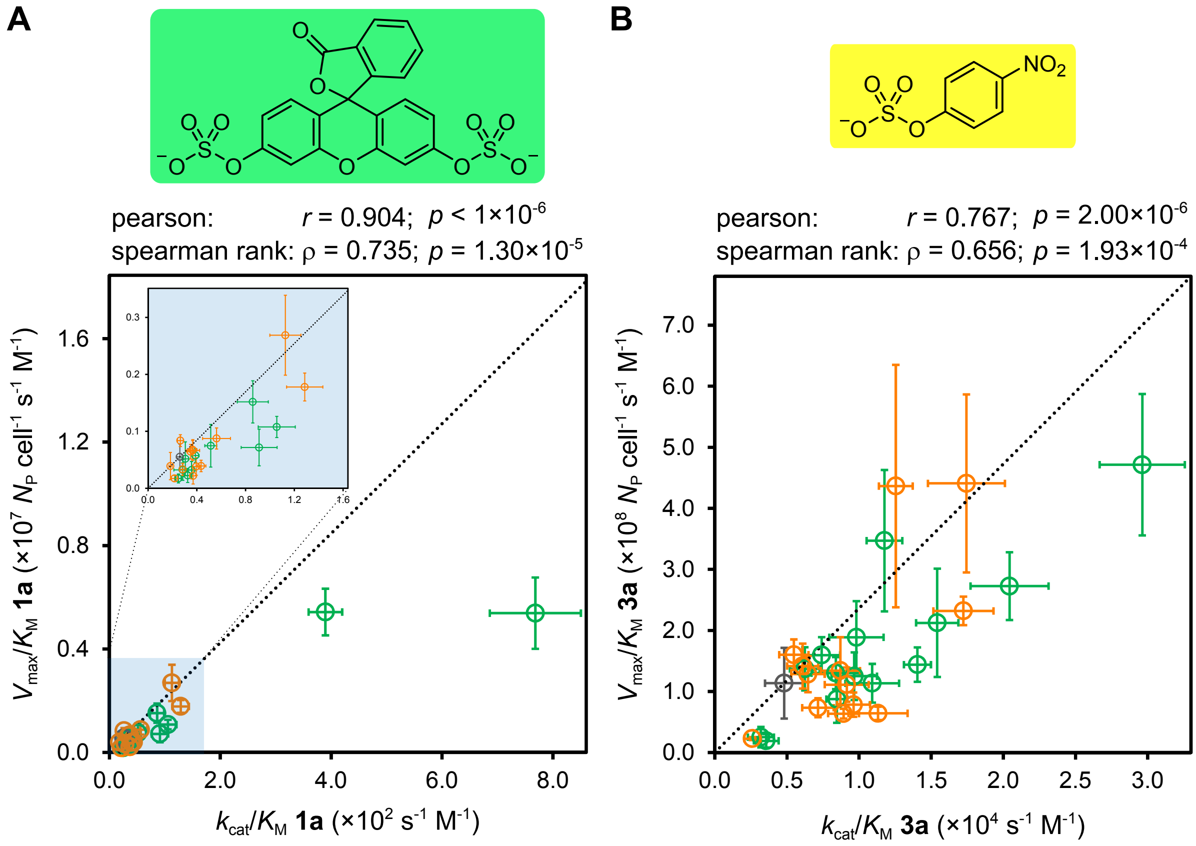


**Figure S17:** Correlation analysis of catalytic performance for autodisplayed ($V_{max}/K_M$) and purified ($k_{cat}/K_M$) *Sp*AS1 variants toward fluorescein disulfate (**A**) and 4-nitrophenyl sulfate (**B**). The dotted line is the correlation that assumes that all *Sp*AS1-variants have the same expression levels as the wild type.

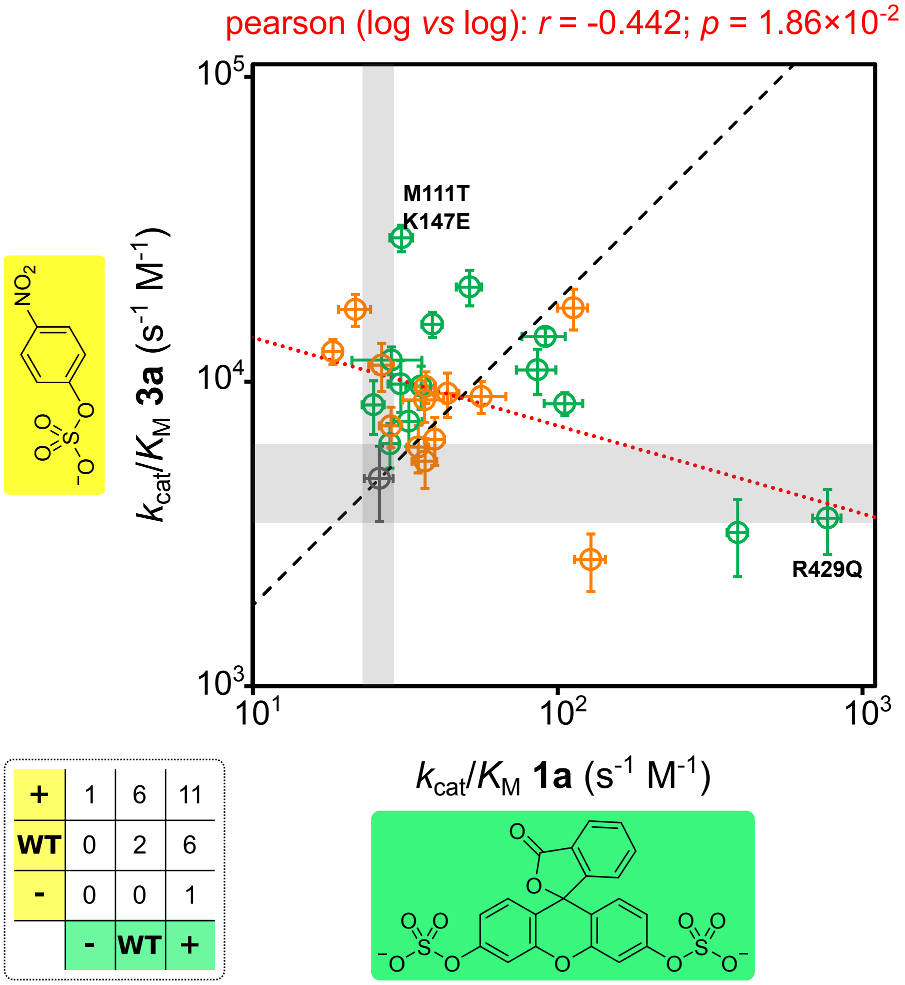


**Figure S18:** Trade-off between the catalytic performance of *Sp*AS1 variants toward 4-nitrophenyl sulfate **3a** *vs* fluorescein disulfate **1a**. Correlation analysis of the log-values of $k_{cat}/K_M$ for both substrates for each *Sp*AS1 variant shows a significant ($p < 5\times10^{-2}$) negative correlation ($r$ = -0.442). This observation is highlighted by the fact that *Sp*AS1$^{R429Q}$, the top performer for fluorescein disulfate **1a**, shows no significant improvement for activity toward 4-nitrophenyl sulfate **3a** and *vice versa* (for *Sp*AS1$^{M111T/K147E}$). In addition only 11 out of the 27 selected variants showed significantly improved performance for both substrates (matrix inset in the lower left corner). The grey zones indicate the possible variation in activity levels toward both substrates for *Sp*AS1$^{WT}$.

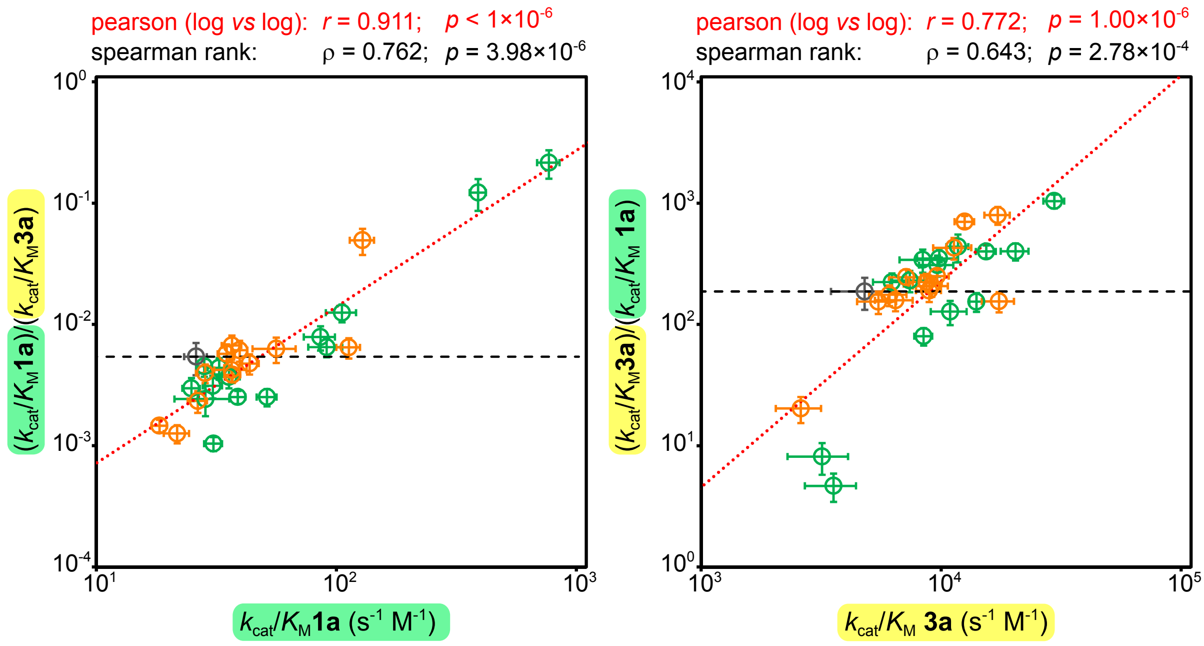


**Figure S19:** Trade-off between the catalytic performance of *Sp*AS1 variants toward fluorescein disulfate **1a** *vs.* 4-nitrophenyl sulfate **3a**. Increased catalytic efficiency ($k_{cat}/K_M$) toward fluorescein disulfate **1a** results in improved specificity toward fluorescein disulfate **1a** over 4-nitrophenyl sulfate **3a** (($k_{cat}/K_M$ **1a**)/($k_{cat}/K_M$ **3a**), relative to wild-type) (panel **A**) and and *vice versa* (**B**). The black dashed line is the hypothetical situation in which the ratio between the catalytic efficiencies toward both substrates ($k_{cat}/K_M$ **1a**)/($k_{cat}/K_M$ **3a**), as observed for *Sp*AS1$^{WT}$, is constant.

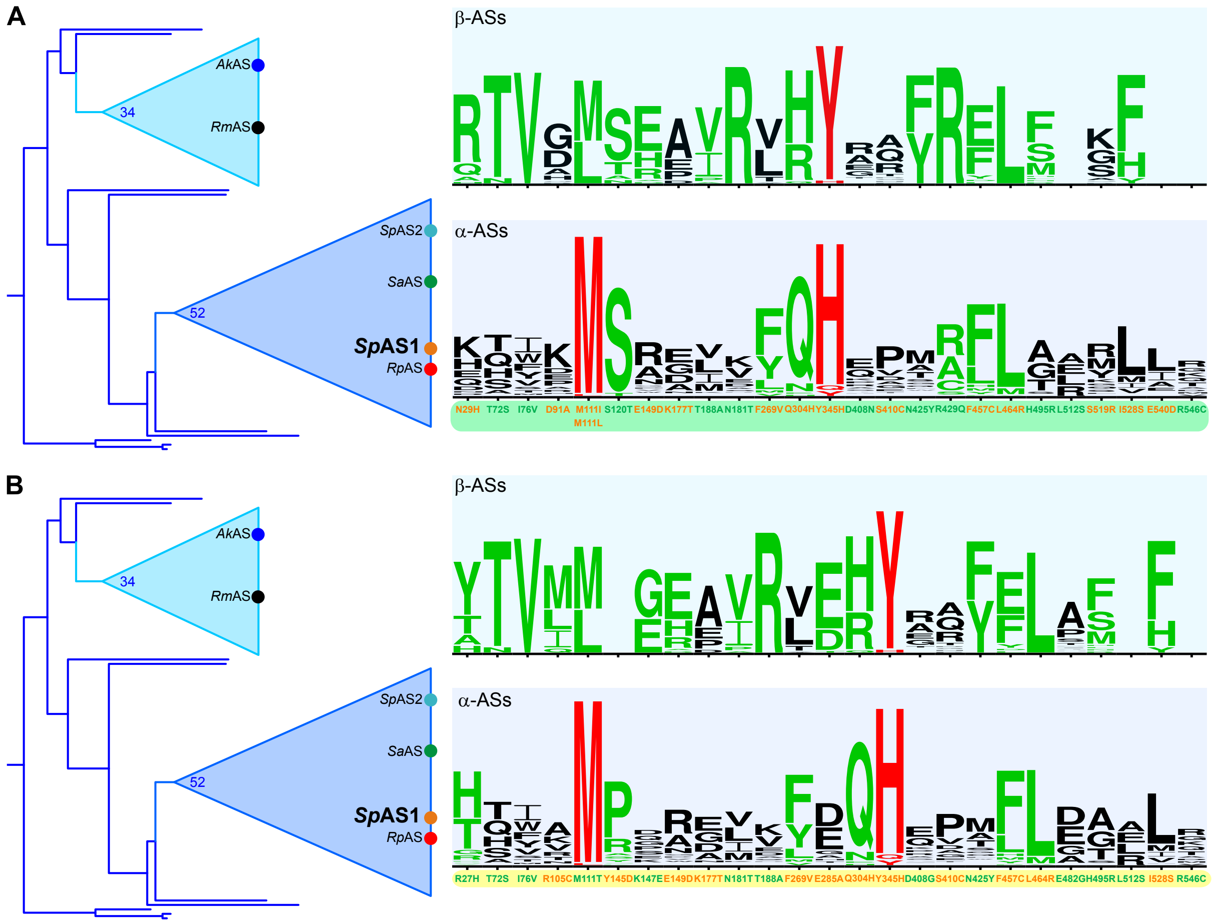


**Figure S20:** Phylogenetic relationships within the clade of the recently described dimeric arylsulfatases[3,12] to which *Sp*AS1 belongs. The logo representation of the amino acid conservation in the respective clades is displayed for the mutated positions in *Sp*AS1 as found in variants with improved activity toward fluorescein disulfate **1a** (panel **A**) and 4-nitrophenyl sulfate **3a** (**B**). Among the α-ASs there is essentially only strong conservation (red) position M111 and Y345 and for the most part moderate (green) to no (black) conservation for all other 29 positions. For M111 the amino acid found in *Sp*AS1[WT] is identical to the conserved residue. For the analogous position to Y345 the conserved residue is a histidine. As a result the Y345H amino acid substitution can be classified as a back-to-consensus mutation,[13] a phenomenon observed[14,15] and exploited[14,16–18] during directed evolution campaigns.

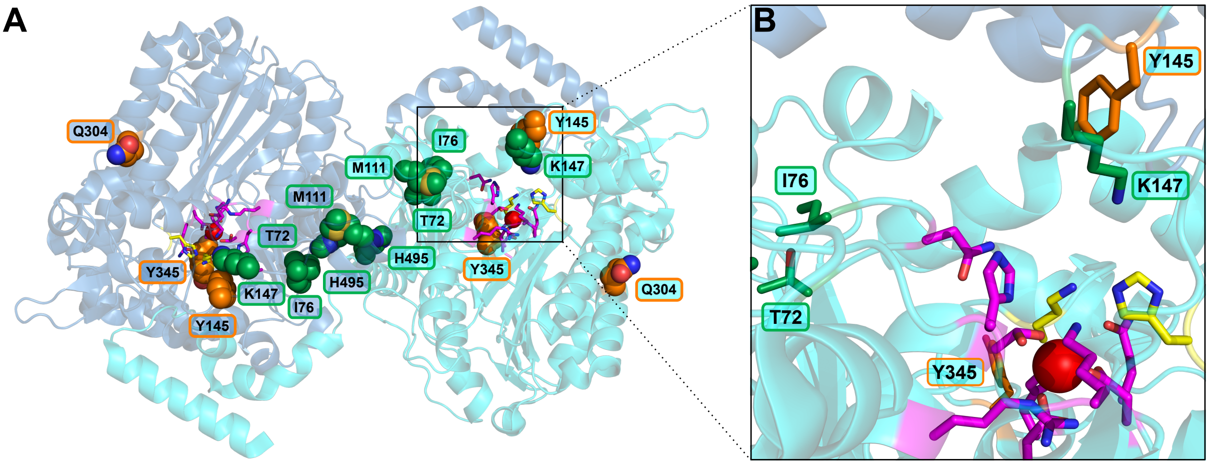


**Figure S21:** Positions of mutations in *Sp*AS1 variants with >3-fold improved activity toward 4-nitrophenylsulfate **3a** (PDB ID for monomer: 4UPI). Mutations indicated in green and orange were introduced during mutagenic PCRs with nucleotide analogs dPTP and 8-oxo-dGTP respectively. The conserved active site groups[3,4] are indicated in yellow (putative leaving group stabilizing residues H234 and K341), magenta (all other putative conserved active site residues), and red (active site metal ion). (**A**) Positions of all residues that are mutated in the five *Sp*AS1 variants that have a >3-fold increased $k_{cat}/K_M$ toward 4-nitrophenyl sulfate **3a**. (**B**) Enlarged view of the region indicated in panel A, highlighting mutated positions in the loop that contains conserved active site residue N74 (T72 and I76) and a loop at the edge of the active site in which multiple mutated positions were observed (Y145, K147, E149 (the latter not shown in this Figure)).

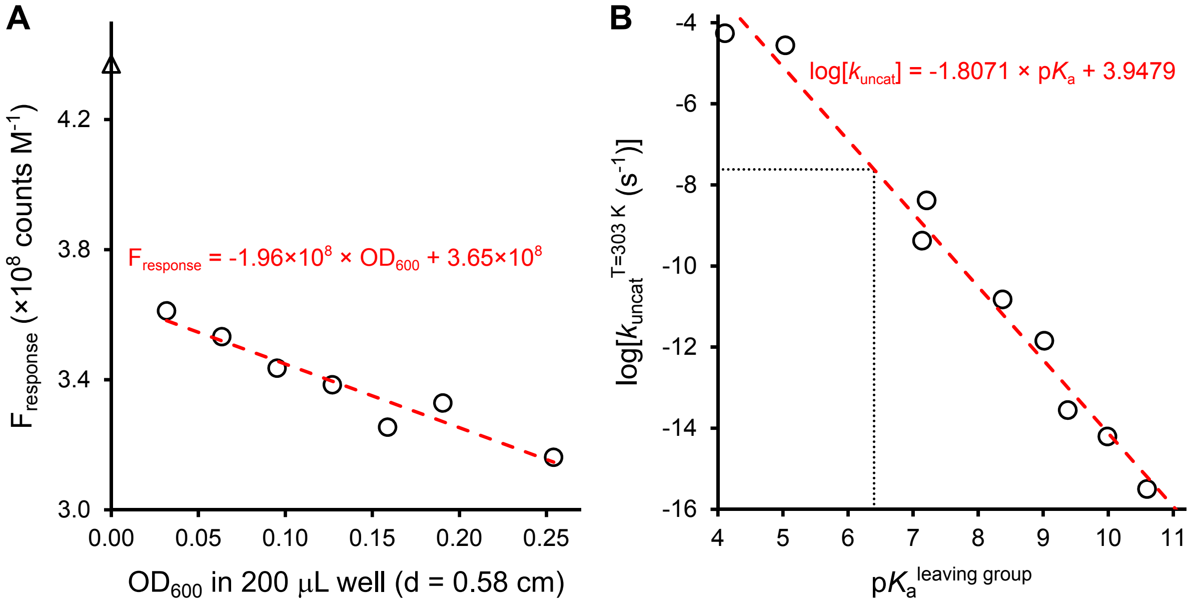


**Figure S22:** Calibration and correction of experimentally determined initial rates of fluorescein **1a** hydrolysis. (**A**) Correlation of fluorescence response of fluorescein **1b** ($F_{response}$ in counts $M^{-1}$) with the amount of *E. cloni* 10G cells expressing autodisplayed *Sp*AS1$^{WT}$ present (represented as the background corrected $OD_{600}$ in a microtiterplate well filled with 200 μL reaction mix). (**B**) Linear free energy relationship (LFER) for the rate constant of uncatalyzed hydrolysis of arylsulfate esters at 30 °C ($k_{uncat}^{T=303\ K}$), calculated using the published enthalpy ($\Delta H$) and entropy ($\Delta S$) values for the uncatalyzed hydrolysis of 9 arylsulfate monoesters[6] according to equation 15 and 16. From this LFER The uncatalyzed rate constant ($k_{uncat}$) for fluorescein disulfate **1a** (p$K_a^{leaving\ group}$ = 6.4, dotted line) was determined to be $10^{-1.8071\times6.4+3.9479}$ = $2.4\times10^{-8}$ $s^{-1}$. Initial rate of uncatalyzed fluorescein disulfate **1a** hydrolyis at [**1a**] = 0.5 mM was experimentally observed to be $1.2\times10^{-11}$ M $s^{-1}$. The expected rate would be $k_{uncat}\times$[S] = $2.4\times10^{-8}\times0.0005 = 1.2\times10^{-11}$ M $s^{-1}$.

#### 4 Supporting Tables

**Table S1:** Library characteristics.

| mutagenic nucleotide | library size[a] | $\lambda_{mutation}$[b] | droplets sorted[c] | fold oversampling |
|---|---|---|---|---|
| dPTP | $1.8\times10^5$ | $2.3\pm1.7$ | $1.8\times10^6$ | 3.5 |
| 8-oxo-dGTP | $4.9\times10^5$ | $3.7\pm2.6$ | $6.8\times10^6$ | 4.9 |

[a]Calculated from the transformation efficiency (in cfu $\mu g^{-1}$) and the amount of DNA transformed.

[b]Calculated from sequencing data obtained for 12 randomly picked variants for each library.

[c]Includes all droplets. Cells are encapsulated into the droplets according to a Poisson distribution with $\lambda$ = 0.35. As a result the total number of variants tested = 0.35×droplets sorted.

**Table S2:** Activity parameters for enzyme-catalyzed hydrolysis of sulfate monoester **1a** by *E. coli* cells displaying *Sp*AS1-variants generated using mutagenic oligonucleotide **dPTP**

| variant | mutations (non-synonymous) | (synonymous) | $V_{max}$[b] ($N_P$ cell$^{-1}$ s$^{-1}$) | $K_M$ (mM) | $V_{max}/K_M$ ($N_P$ cell$^{-1}$ s$^{-1}$ M$^{-1}$) | ratio[c] |
|---|---|---|---|---|---|---|
| wild type[a] | - | - | $(1.4\pm0.8)\times10^3$ | 2.9±1.7 | $(5.5\pm2.3)\times10^5$ | - |
| 1C3 | N425Y | - | $(2.4\pm0.2)\times10^3$ | 2.3±0.3 | $(1.1\pm0.2)\times10^6$ | 2.0±0.9 |
| 1E4 | I76V | - | $(2.9\pm0.3)\times10^3$ | 5.0±0.9 | $(5.7\pm1.2)\times10^5$ | 1.0±0.5 |
| 1B5 | N181T, L512S | *L89L, G184G, D506D* | $(3.6\pm0.4)\times10^3$ | 2.4±0.5 | $(1.5\pm0.4)\times10^6$ | 2.8±1.3 |
| 1C5 | S120T, R429Q | - | $(4.5\pm0.2)\times10^3$ | 0.83±0.13 | $(5.4\pm0.9)\times10^6$ | 9.8±4.3 |
| 1C8 | T72S, H495R | *K101K, R434R* | $(4.3\pm1.2)\times10^3$ | 5.7±2.4 | $(7.5\pm3.7)\times10^5$ | 1.4±0.9 |
| 1D9, 1E9 | M111T, K147E | *D407D* | n.d. | n.d | n.d. | n.d. |
| 1D10 | T188A | - | $(1.8\pm0.5)\times10^3$ | 5.6±2.2 | $(3.2\pm1.5)\times10^5$ | 0.58±0.36 |
| 1E10 | E482G | *K148K* | $(2.1\pm0.7)\times10^3$ | 6.5±3.0 | $(3.2\pm1.8)\times10^5$ | 0.59±0.41 |
| 1H10 | R429Q | - | $(4.4\pm0.3)\times10^3$ | 0.82±0.20 | $(5.4\pm1.4)\times10^6$ | 9.8±4.7 |
| 1F11 | D408N | - | $(1.6\pm0.5)\times10^3$ | 7.1±3.2 | $(2.2\pm1.2)\times10^5$ | 0.41±0.28 |
| 1F12 | R27H | *G127G* | $(6.0\pm1.4)\times10^2$ | 3.5±1.4 | $(1.7\pm0.8)\times10^5$ | 0.31±0.19 |
| 2B4 | R546C | *V324V* | $(2.9\pm0.7)\times10^3$ | 4.0±1.5 | $(7.2\pm3.2)\times10^6$ | 1.3±0.8 |
| 2D8 | D408G | - | $(2.0\pm0.6)\times10^3$ | 3.8±1.9 | $(5.2\pm3.0)\times10^5$ | 0.94±0.66 |
| 3C1 | V92A, V466A, K541R | - | $(7.0\pm1.1)\times10^2$ | 2.1±0.7 | $(3.3\pm1.2)\times10^5$ | 0.60±0.32 |

[a] Average of xx separate measurements. Due to co-variance of $V_{max}$ and $K_M$, $V_{max}/K_M$ is more reliable than would be expected from the values for $V_{max}$ and $K_M$.

[b] Expressed as the number of fluorescein molecules ($N_P$) formed per *E. coli* cell per second.

[c] $(V_{max}/K_M)_{mutant}/(V_{max}/K_M)_{WT}$.

**Table S3:** Activity parameters for enzyme-catalyzed hydrolysis of sulfate monoester **1a** by *E. coli* cells displaying *Sp*AS1-variants generated using mutagenic oligonucleotide **8-oxo-dGTP**

| variant | mutations (non-synonymous) | (synonymous) | $V_{max}$[b] ($N_P$ cell$^{-1}$ s$^{-1}$) | $K_M$ (mM) | $V_{max}/K_M$ ($N_P$ cell$^{-1}$ s$^{-1}$ M$^{-1}$) | ratio[c] |
|---|---|---|---|---|---|---|
| wild type[a] | - | - | $(1.5\pm0.8)\times10^3$ | $2.9\pm1.7$ | $(5.5\pm2.3)\times10^5$ | - |
| 1B8 | Q304H, Y345H | *D303D*, *I354I* | $(5.0\pm0.5)\times10^3$ | $1.9\pm0.4$ | $(2.7\pm0.7)\times10^6$ | $4.9\pm2.4$ |
| 1E9, 3C12 | F269V, S410C, I528S | *L464L* | $(2.3\pm0.2)\times10^3$ | $2.6\pm0.5$ | $(8.7\pm1.9)\times10^5$ | $1.6\pm0.7$ |
| 1A10 | S519R, E540D | *R429R, L487L* | $(3.8\pm0.2)\times10^3$ | $2.1\pm0.3$ | $(1.8\pm0.2)\times10^6$ | $3.2\pm1.4$ |
| 1B11 | Y145D | - | $(5.7\pm0.9)\times10^2$ | $3.3\pm0.9$ | $(1.7\pm0.5)\times10^5$ | $0.31\pm0.16$ |
| 1C12 | D91A, M111L | *P514P* | $(2.4\pm0.3)\times10^3$ | $3.6\pm0.8$ | $(6.7\pm1.8)\times10^5$ | $1.2\pm0.6$ |
| 2E2 | R105C | *A539A* | $(8.3\pm2.2)\times10^2$ | $2.1\pm1.2$ | $(3.9\pm2.4)\times10^5$ | $0.71\pm0.52$ |
| 2F6, 3B11 | E149D, K177T | - | $(1.4\pm0.2)\times10^3$ | $2.2\pm0.5$ | $(6.6\pm1.7)\times10^5$ | $1.2\pm0.6$ |
| 2F11 | N29H | - | $(1.7\pm0.4)\times10^3$ | $4.4\pm1.5$ | $(3.8\pm1.5)\times10^5$ | $0.69\pm0.40$ |
| 3E2 | M111I | - | $(1.6\pm0.1)\times10^3$ | $2.5\pm0.5$ | $(6.4\pm1.3)\times10^6$ | $1.2\pm0.5$ |
| 3G5 | E285A | *F131F, G339G* | $(1.58\pm0.08)\times10^3$ | $1.9\pm0.2$ | $(8.4\pm1.0)\times10^5$ | $1.5\pm0.6$ |
| 3B10 | E164K, K177Q | (M1R) | $(1.2\pm0.2)\times10^3$ | $3.5\pm1.0$ | $(3.3\pm1.1)\times10^5$ | $0.60\pm0.31$ |
| 3F11 | L464R | *L461L* | $(6.0\pm1.9)\times10^2$ | $2.8\pm1.6$ | $(2.2\pm1.4)\times10^5$ | $0.40\pm0.31$ |
| 3A12 | F457C | - | $(1.5\pm0.2)\times10^3$ | $3.8\pm0.8$ | $(3.9\pm1.0)\times10^5$ | $0.71\pm0.34$ |

[a] Average of xx separate measurements. Due to co-variance of $V_{max}$ and $K_M$, $V_{max}/K_M$ is more reliable than would be expected from the values for $V_{max}$ and $K_M$.

[b] Expressed as the number of fluorescein molecules ($N_P$) formed per *E. coli* cell per second.

[c] $(V_{max}/K_M)_{mutant}/(V_{max}/K_M)_{WT}$.

**Table S4:** Activity parameters for enzyme-catalyzed hydrolysis of sulfate monoester **3a** by *E. coli* cells displaying *Sp*AS1-variants generated using mutagenic oligonucleotide **dPTP**

| variant | mutations (non-synonymous) | (synonymous) | $V_{max}$[b] ($N_P$ cell$^{-1}$ s$^{-1}$) | $K_M$ (mM) | $V_{max}/K_M$ ($N_P$ cell$^{-1}$ s$^{-1}$ M$^{-1}$) | ratio[c] |
|---|---|---|---|---|---|---|
| wild type[a] | - | - | $(8.2\pm3.5)\times10^5$ | $8.2\pm4.0$ | $(1.1\pm0.6)\times10^8$ | - |
| 1C3 | N425Y | - | $(3.3\pm0.6)\times10^5$ | $3.8\pm1.5$ | $(8.8\pm3.9)\times10^7$ | $0.8\pm0.5$ |
| 1E4 | I76V | - | $(1.5\pm0.3)\times10^6$ | $6.7\pm2.5$ | $(2.1\pm0.9)\times10^8$ | $1.9\pm1.2$ |
| 1B5 | N181T, L512S | *L89L, G184G, D506D* | $(6.9\pm0.9)\times10^5$ | $6.1\pm1.5$ | $(1.1\pm0.3)\times10^8$ | $1.0\pm0.6$ |
| 1C5 | S120T, R429Q | - | $(4.3\pm1.7)\times10^5$ | $17\pm9$ | $(2.5\pm1.7)\times10^7$ | $0.22\pm0.19$ |
| 1C8 | T72S, H495R | *K101K, R434R* | $(1.1\pm0.1)\times10^6$ | $4.2\pm0.8$ | $(2.7\pm0.6)\times10^8$ | $2.4\pm1.3$ |
| 1D9, 1E9 | M111T, K147E | *D407D* | $(9.4\pm0.8)\times10^5$ | $2.0\pm0.5$ | $(4.7\pm1.2)\times10^8$ | $4.2\pm2.4$ |
| 1D10 | T188A | - | $(7.8\pm1.2)\times10^5$ | $6.3\pm1.7$ | $(1.3\pm0.4)\times10^8$ | $1.1\pm0.7$ |
| 1E10 | E482G | *K148K* | $(4.7\pm0.4)\times10^5$ | $1.4\pm0.4$ | $(3.5\pm1.2)\times10^8$ | $3.1\pm1.9$ |
| 1H10 | R429Q | - | $(2.5\pm0.9)\times10^5$ | $13\pm6$ | $(1.9\pm1.1)\times10^7$ | $0.17\pm0.13$ |
| 1F11 | D408N | - | $(6.3\pm0.5)\times10^5$ | $4.07\pm0.7$ | $(1.6\pm0.3)\times10^8$ | $1.4\pm0.8$ |
| 1F12 | R27H | *G127G* | $(1.4\pm0.2)\times10^6$ | $11\pm2$ | $(1.3\pm0.3)\times10^8$ | $1.1\pm0.6$ |
| 2B4 | R546C | *V324V* | $(9.1\pm0.9)\times10^5$ | $6.4\pm1.1$ | $(1.4\pm0.3)\times10^8$ | $1.3\pm0.7$ |
| 2D8 | D408G | - | $(1.9\pm0.3)\times10^6$ | $10\pm3$ | $(1.9\pm0.6)\times10^8$ | $1.7\pm1.0$ |
| 3C1 | V92A, V466A, K541R | - | $(1.0\pm0.1)\times10^6$ | $7.3\pm1.6$ | $(1.4\pm0.4)\times10^8$ | $1.2\pm0.7$ |

[a] Average of 14 separate measurements. Due to co-variance of $V_{max}$ and $K_M$, $V_{max}/K_M$ is more reliable than would be expected from the values for $V_{max}$ and $K_M$.

[b] Expressed as the number of 4-nitrophenolate molecules ($N_P$) formed per *E. coli* cell per second.

[c] $(V_{max}/K_M)_{mutant}/(V_{max}/K_M)_{WT}$.

**Table S5:** Activity parameters for enzyme-catalyzed hydrolysis of sulfate monoester **3a** by *E. coli* cells displaying *Sp*AS1-variants generated using mutagenic oligonucleotide **8-oxo-dGTP**

| variant | mutations (non-synonymous) | (synonymous) | $V_{max}$[b] ($N_P$ cell$^{-1}$ s$^{-1}$) | $K_M$ (mM) | $V_{max}/K_M$ ($N_P$ cell$^{-1}$ s$^{-1}$ M$^{-1}$) | ratio[c] |
|---|---|---|---|---|---|---|
| wild type[a] | - | - | $(8.2\pm3.5)\times10^5$ | 8.2±4.0 | $(1.1\pm0.6)\times10^8$ | - |
| 1B8 | Q304H, Y345H | *D303D*, *I354I* | $(5.5\pm0.5)\times10^5$ | 1.2±0.4 | $(4.4\pm1.5)\times10^8$ | 3.9±2.4 |
| 1E9, 3C12 | F269V, S410C, I528S | *L464L* | $(4.4\pm0.3)\times10^5$ | 6.9±0.8 | $(6.4\pm0.8)\times10^7$ | 0.56±0.30 |
| 1A10 | S519R, E540D | *R429R, L487L* | $(1.9\pm0.1)\times10^5$ | 8.5±0.8 | $(2.3\pm0.2)\times10^7$ | 0.20±0.10 |
| 1B11 | Y145D | - | $(1.23\pm0.06)\times10^6$ | 5.3±0.5 | $(2.3\pm0.2)\times10^8$ | 2.0±1.1 |
| 1C12 | D91A, M111L | *P514P* | $(1.09\pm0.09)\times10^6$ | 6.8±0.9 | $(1.6\pm0.3)\times10^8$ | 1.4±0.8 |
| 2E2 | R105C | *A539A* | $(1.6\pm0.3)\times10^6$ | 3.7±1.5 | $(4.4\pm2.0)\times10^8$ | 3.8±2.6 |
| 2F6, 3B11 | E149D, K177T | - | $(1.2\pm0.3)\times10^6$ | 9.0±3.1 | $(1.3\pm0.5)\times10^8$ | 1.2±0.8 |
| 2F11 | N29H | - | $(6.9\pm0.7)\times10^5$ | 5.4±1.1 | $(1.3\pm0.3)\times10^8$ | 1.1±0.6 |
| 3E2 | M111I | - | $(1.3\pm0.2)\times10^6$ | 9.5±2.0 | $(1.4\pm0.4)\times10^8$ | 1.2±0.7 |
| 3G5 | E285A | *F131F, G339G* | $(4.3\pm0.4)\times10^5$ | 6.7±1.0 | $(6.4\pm1.1)\times10^7$ | 0.57±0.30 |
| 3B10 | E164K, K177Q | (M1R) | $(7.2\pm0.8)\times10^5$ | 10±2 | $(7.3\pm1.6)\times10^7$ | 0.65±0.36 |
| 3F11 | L464R | *L461L* | $(3.7\pm0.4)\times10^5$ | 4.8±1.1 | $(7.8\pm2.0)\times10^7$ | 0.69±0.39 |
| 3A12 | F457C | - | $(9.1\pm1.2)\times10^5$ | 8.2±1.8 | $(1.1\pm0.3)\times10^8$ | 1.0±0.6 |

[a] Average of 14 separate measurements. Due to co-variance of $V_{max}$ and $K_M$, $V_{max}/K_M$ is more reliable than would be expected from the values for $V_{max}$ and $K_M$.

[b] Expressed as the number of 4-nitrophenolate molecules ($N_P$) formed per *E. coli* cell per second.

[c] $(V_{max}/K_M)_{mutant}/(V_{max}/K_M)_{WT}$.

**Table S6:** Kinetic parameters for enzyme-catalyzed hydrolysis of sulfate monoester **1a** by purified *Sp*AS1-variants[a]

| mutations | mutagenic nucleotide | $k_{cat}$ ($s^{-1}$) | $K_M$ (mM) | $K_{SI}$ (mM) | $k_{cat}/K_M$ ($s^{-1}$ $M^{-1}$) |
|---|---|---|---|---|---|
| wild type | - | 0.116±0.004 | 4.5±0.5 | n/a | 26±3 |
| R429Q | dPTP | 0.77±0.04 | 1.0±0.1 | 21±4 | 768±82 |
| S120T, R429Q | dPTP | 0.40±0.01 | 1.02±0.07 | 21±3 | 390±30 |
| S519R, E540D | 8-oxo-dGTP | 0.22±0.01 | 1.7±0.2 | 47±11 | 129±15 |
| Q304H, Y345H | 8-oxo-dGTP | 0.40±0.02 | 3.5±0.3 | 33±6 | 113±13 |
| N425Y | dPTP | 0.39±0.03 | 3.7±0.5 | 22±4 | 106±15 |
| R546C | dPTP | 0.25±0.02 | 2.7±0.4 | 32±8 | 91±15 |
| N181T, L512S | dPTP | 0.25±0.02 | 2.9±0.4 | 36±8 | 86±13 |
| F269V, S410C, I528S | 8-oxo-dGTP | 0.14±0.01 | 2.5±0.5 | 58±25 | 56±11 |
| T72S, H495R | dPTP | 0.159±0.005 | 3.1±0.3 | n/a | 51±5 |
| F457C | 8-oxo-dGTP | 0.145±0.005 | 3.3±0.3 | n/a | 44±4 |
| N29H | 8-oxo-dGTP | 0.127±0.003 | 3.2±0.2 | n/a | 40±3 |
| I76V | dPTP | 0.102±0.003 | 2.6±0.2 | n/a | 39±3 |
| L464R | 8-oxo-dGTP | 0.150±0.005 | 4.1±0.3 | n/a | 37±3 |
| D91A, M111L | 8-oxo-dGTP | 0.144±0.005 | 3.9±0.3 | n/a | 37±3 |
| E149D, K177T | 8-oxo-dGTP | 0.14±0.01 | 3.9±0.5 | 76±31 | 37±7 |
| T188A | dPTP | 0.116±0.004 | 3.3±0.3 | n/a | 36±4 |
| M111I | 8-oxo-dGTP | 0.132±0.004 | 3.8±0.3 | n/a | 35±3 |
| D408N | dPTP | 0.109±0.004 | 3.4±0.3 | n/a | 32±3 |
| M111T, K147E | dPTP | 0.074±0.002 | 2.4±0.2 | n/a | 31±3 |
| D408G | dPTP | 0.115±0.004 | 3.8±0.3 | n/a | 31±3 |
| E482G | dPTP | 0.14±0.02 | 4.8±1.1 | 56±32 | 28±7 |
| E164K, K177Q | 8-oxo-dGTP | 0.113±0.003 | 4.0±0.3 | n/a | 28±2 |
| V92A, V466A, K541R | dPTP | 0.118±0.003 | 4.2±0.3 | n/a | 28±2 |
| E285A | 8-oxo-dGTP | 0.092±0.003 | 3.5±0.3 | n/a | 26±2 |
| R27H | dPTP | 0.131±0.004 | 5.3±0.4 | n/a | 25±2 |
| Y145D | 8-oxo-dGTP | 0.121±0.006 | 5.6±0.6 | n/a | 22±3 |
| R105C | 8-oxo-dGTP | 0.0393±0.0009 | 2.1±0.2 | n/a | 18±1 |

**Table S7:** Kinetic parameters for enzyme-catalyzed hydrolysis of sulfate monoester **3a** by purified *Sp*AS1-variants

| mutations | mutagenic nucleotide | $k_{cat}$ ($s^{-1}$) | $K_M$ (mM) | $K_{SI}$ (mM) | $k_{cat}/K_M$ ($s^{-1}$ $M^{-1}$) |
|---|---|---|---|---|---|
| wild type | - | 34±5 | 7.1±1.7 | 19±5 | (4.8±1.3)×$10^3$ |
| M111T, K147E | dPTP | 100±4 | 3.4±0.3 | 25±2 | (3.0±0.3)×$10^4$ |
| T72S, H495R | dPTP | 63±2 | 3.1±0.4 | n/a | (2.0±0.3)×$10^4$ |
| Q304H, Y345H | 8-oxo-dGTP | 52±5 | 3.0±0.4 | 4.5±0.6 | (1.7±0.4)×$10^4$ |
| Y145D | 8-oxo-dGTP | 40±2 | 2.4±0.3 | 53±7 | (1.7±0.2)×$10^4$ |
| I76V | dPTP | 99±5 | 6.5±0.5 | 51±6 | (1.5±0.1)×$10^4$ |
| R546C | dPTP | 49±1 | 3.5±0.2 | n/a | (1.40±0.09)×$10^4$ |
| R105C | 8-oxo-dGTP | 34±1 | 2.7±0.2 | 239±56 | (1.3±0.1)×$10^4$ |
| E482G | dPTP | 45±1 | 3.8±0.4 | n/a | (1.2±0.1)×$10^4$ |
| E285A | 8-oxo-dGTP | 68±6 | 6.0±0.9 | n/a | (1.1±0.2)×$10^4$ |
| N181T, L512S | dPTP | 50±4 | 4.6±0.7 | 76±21 | (1.1±0.2)×$10^4$ |
| D408G | dPTP | 51±5 | 5.2±0.9 | 23±4 | (9.8±1.9)×$10^3$ |
| T188A | dPTP | 49±4 | 5.1±0.7 | 32±5 | (9.7±1.6)×$10^3$ |
| L464R | 8-oxo-dGTP | 42±2 | 4.3±0.5 | 29±3 | (9.6±1.1)×$10^3$ |
| F457C | 8-oxo-dGTP | 46±4 | 5.0±0.7 | 33±5 | (9.1±1.5)×$10^3$ |
| F269V, S410C, I528S | 8-oxo-dGTP | 18.0±0.5 | 2.0±0.2 | n/a | (8.9±1.1)×$10^3$ |
| E149D, K177T | 8-oxo-dGTP | 33±2 | 3.8±0.5 | 57±10 | (8.7±1.4)×$10^3$ |
| N425Y | dPTP | 30.1±0.7 | 3.6±0.3 | n/a | (8.5±0.7)×$10^3$ |
| R27H | dPTP | 44±4 | 5.2±0.9 | 26±5 | (8.4±1.7)×$10^3$ |
| D408N | dPTP | 49±4 | 6.7±1.0 | 20±3 | (7.4±1.3)×$10^3$ |
| E164K, K177Q | 8-oxo-dGTP | 29±2 | 4.1±0.6 | 42±7 | (7.2±1.1)×$10^3$ |
| N29H | 8-oxo-dGTP | 35±3 | 5.4±0.9 | 42±7 | (6.5±1.2)×$10^3$ |
| V92A, V466A, K541R | dPTP | 37±3 | 5.9±0.8 | 16±2 | (6.2±1.0)×$10^3$ |
| M111I | 8-oxo-dGTP | 28±2 | 4.7±0.7 | 24±4 | (6.1±1.1)×$10^3$ |
| D91A, M111L | 8-oxo-dGTP | 39±4 | 7.0±1.1 | 20±3 | (5.5±1.0)×$10^3$ |
| R429Q | dPTP | 26±3 | 7.3±1.5 | 30±7 | (3.6±0.9)×$10^3$ |
| S120T, R429Q | dPTP | 11.3±0.8 | 3.5±1.0 | n/a | (3.2±0.9)×$10^3$ |
| S519R, E540D | 8-oxo-dGTP | 12±1 | 4.4±0.8 | 48±12 | (2.6±0.6)×$10^3$ |

**Table S8:** Synthetic oligonucleotides[a] used in this study.

| | | |
|---|---|---|
| pBAD-AT | cloning | |
| forward[b] | 5’-gcgcgcctcgag(**cat**)$_6$(gca)$_3$ATGACCCAAAACCGCAACATTCTGTGG-3’ | *Xho*I |
| reverse | 5’-cgcgcgggtaccCTACGAGTTTTCACCAAGAATGCGC-3’ | *Kpn*I |
| pBAD-AT | error-prone PCR | |
| forward | 5’-CATAGCACCATGGAGCTCGAGCAT-3’ | *Xho*I |
| reverse | 5’-CACTTCTTTTGTAGGATTAAGGGTACC-3’ | *Kpn*I |
| pASKIBA5$^+$ | cloning | |
| forward | 5’-tacgccctcgagcATGACCCAAAACCGCAACATTCTGTGG-3’ | *Xho*I |
| reverse | 5’-cgcgcgctgcagCTACGAGTTTTCACCAAGAATGCGC-3’ | *Pst*I |
| reverse R546C | 5’-cgcgcgctgcagCTACGAGTTTTCACCAAGAATGCaC-3’ | *Pst*I |

[a] Indicated restriction sites are underlined, non-matching base-pairs in lower case.

[b] His$_6$-tag coded by bases indicated in bold.
